# Broadscale evolutionary analysis of eukaryotic DDE transposons

**DOI:** 10.1101/2021.09.26.461848

**Authors:** Mathilde Dupeyron, Tobias Baril, Alexander Hayward

**Author notes:** These authors contributed equally (Mathilde Dupeyron, Tobias Baril). **Author Contributions:** MD curated sequence alignments and host data, inferred phylogenies, performed prokaryotic sequence analyses, and produced host range heatmaps, FigTree files, and cophylogeny tanglegrams. TB developed the scripts for automated BLAST, sequence extraction, taxonomic information extraction, and sequence alignment, performed sequence-clustering and principle components analyses, and produced graphs summarising host range and estimated horizontal transfers. AH conceived and coordinated the study, performed additional phylogenetic, horizontal transfer and cophylogeny analyses, produced the figures, and drafted the manuscript. All authors contributed to the final version of the manuscript. **Competing Interest Statement:** The authors declare that they have no competing interests.

## Abstract

DDE transposons are widespread selfish genetic elements, often comprising a large proportion of eukaryotic genomic content. DDE transposons have also made important contributions to varied host functions during eukaryotic evolution, and their transposases may be the most abundant and ubiquitous genes in nature. Yet much remains unknown about their basic biology. We employ a broadscale screen of DDE transposase diversity to characterise major evolutionary patterns for all 19 DDE transposon superfamilies. We identify considerable variation in DDE transposon superfamily size, and find a dominant association with animal hosts. While few DDE transposon superfamilies specialise in plants or fungi, the four largest superfamilies contain major plant-associated clades, at least partially underlying their relative success. We recover a pattern of host conservation among DDE transposon lineages, punctuated by occasional horizontal transfer to distantly related hosts. Host range and horizontal transfer are strongly positively correlated with DDE transposon superfamily size, arguing against variation in the capacity for generalism. We find that rates of horizontal transfer decrease sharply with increasing levels of host taxonomy, supporting the existence of host-associated barriers to DDE transposon spread. Overall, despite their relatively simple genetic structure, our results imply that trade-offs in host adaptation are important in defining DDE transposon-host relationships and evolution. In addition, our study provides a phylogenetic framework to facilitate the identification and further analysis of DDE transposons.

## Introduction

Transposons are mobile selfish genetic elements that can proliferate in host genomes (1). Transposons may disrupt host genome function if they jump into coding regions, and as a consequence of their repetitive nature, they can mediate structural changes such as deletions, translocations, duplications, and ectopic recombination (2–5). Consequently, many transposon insertions are deleterious to the host, and their activity is implicated in various diseases (6–9). However, transposon sequences also contain genetic diversity that may be utilized during host genome evolution, including regulatory motifs and protein domains, while the structural rearrangements that transposons facilitate can result in further novelty for selection to act upon (4, 5, 10). Thus, while transposons evolved to promote their own selfish replication, they also offer host benefits, and there is growing evidence of their recruitment for diverse functions during host genomic evolution (2–5, 10–12).

The most diverse and widespread group of transposons in eukaryotic genomes are DDE transposons (2, 13–15). DDE transposons are relatively simple genetic elements, at their most basic consisting of: (i) an intronless transposase gene, which codes for the enzyme that catalyzes transposition, and, (ii) a pair of terminal inverted repeats that act as transposase binding sites (16, 17). DDE transposons are named after the catalytic triad, a coordinated set of three acidic amino acids within the transposase enzyme, which of key importance for catalyzing transposition, and is composed of either two aspartic acids and a glutamic acid (DDE), or in some cases three aspartic acids (DDD).

In addition to vertical transmission via the germline, DDE transposons can spread to new hosts by horizontal transfer, resulting in a flow of novel genetic material among eukaryotes, some of which may be utilised by host genomes (14, 16, 18–20). Indeed, there is an increasing realization of the important contributions made by DDE transposons in particular, to varied and fundamental host functions including adaptive immunity (21), centromere formation (22), and development (23), among others (2, 12).

Most eukaryotic genomes include a proption of DDE transposon content. However, relatively little is known about how host range is structured in DDE transposons. Pronounced differences in host associations have been observed within other major groups of selfish genetic elements. For example, there is great variability in host range and rates of horizontal transfer among retroviral genera (i.e. enveloped LTR retrotransposons) (24). Addressing these aspects in DDE transposons is crucial to elucidate the mechanisms responsible for promoting their spread among hosts, understanding their impact on eukaryotic evolution, and elucidating the factors that drive the varying host interactions of selfish genetic elements more generally.

DDE transposons were the first group of transposon discovered, and they have a comparatively long history of study, and of application in the laboratory as genetic tools for transgenesis and insertional mutagenesis (25, 26). Collectively DDE transposons represent a considerable proportion of sum genetic diversity present within the eukaryotic tree of life, and their transposases may be the most abundant and most ubiquitous genes in nature (15). Yet, much of the evolutionary biology of this important group of selfish genetic elements remains unclear. Here we employ a broadscale screen of DDE transposon diversity to infer phylogeny and characterise major evolutionary patterns for all 19 DDE transposon superfamilies. Specifically, we address the five major outstanding questions discussed below:

1. How does phylogeny and diversity vary among DDE transposon superfamilies?: DDE transposons are grouped into distinct superfamilies that share a common evolutionary origin, but relationships among and within superfamilies are not well understood (2, 13), and little is known about their comparative diversity, or the mechanisms responsible for driving variation among superfamilies.
2. How does host range vary among DDE transposon superfamilies?: DDE transposons may be generalists able to exploit a wide range of hosts across eukaryotic diversity, in line with suggestions that compared to other transposons they are genetically streamlined opportunists (14, 16, 20). Alternatively, groups of DDE transposons may specialise on subsets of eukaryotic diversity (either overlapping or non-overlapping), indicating that DDE transposons undergo some degree of host adaptation in order to successfully exploit a given set of hosts.
3. What is the pattern of horizontal transfer among and within DDE transposon superfamilies?: The frequency and magnitude of horizontal transfer events undertaken by DDE transposons is unclear. Horizontal transfer may occur frequently between distantly related hosts. Alternatively, transfers may occur primarily between closely related hosts, consistent with the existence of barriers to spread, either ecological (i.e. constrained by opportunity) or physiological (i.e. constrained by compatibility). Additionally, the frequency of horizontal transfer may vary among superfamilies, especially if certain superfamilies possess adaptations that predispose them to spread by vectors, or which ensure proliferation in a wide variety of host cellular backgrounds.
4. To what extent is host-DDE transposon cophylogeny congruent? Horizontal transfer to novel hosts is proposed to be an important strategy for transposons to avoid extinction as a consequence of host genomic defences (16, 20, 27). However, the impact of horizontal transfer on host-transposon cophylogeny is untested in DDE transposons, and the relative importance of horizontal versus vertical transmission remains unclear (20). A strong signal of cophylogeny would imply a limited impact of horizontal transfer on host-transposon codiversification. This is expected if vertical transmission is the dominant means by which DDE transposons persist, and horizontal transfer events rarely result in the establishment of successful descendent transposon lineages. Meanwhile, a lack of cophylogenetic signal would suggest a pervasive influence of horizontal transfer in shaping DDE transposon evolution.
5. Do eukaryotic DDE transposon superfamilies include prokaryotic elements? DDE transposons share several characteristics with prokaryotic insertion sequences (2, 28–33). However, the relationship between similar transposons from eukaryote and prokaryote genomes remains confusing. Elucidating this question is important to clarify host relationships among DDE transposons, their evoution, and ultimate origin.

Addressing the above questions offers fundamental insights into the evolutionary dynamics of a major group of selfish genetic element, the DDE transposons, which are a key contributor to eukaryotic genome content, size, and evolvability. Alongside our study, we present a novel resource consisting of a set of annotated phylogenies and corresponding transposase alignments for all DDE transposon superfamilies. This framework offers a detailed context for the future characterisation and identification of DDE transposons, and further analyses into their evolution and host relationships.

## Results and Discussion

### Phylogeny and diversity of DDE transposons

Using queries of online genetic databases we isolated ~29,000 transposase sequences from 1,129 diverse species of eukaryote, after removal of identical transposases from the same host species. Retrieved transposase sequences were analysed together with reference sequences to infer phylogeny for all 19 DDE transposon superfamilies (Figures S1-19). Substantial variation was observed in the number of transposase sequences recovered for each DDE transposon superfamily, with the smallest superfamily, *Novosib,* containing just 10 transposases, while the largest four superfamilies (*PHIS, Mutator, hAT, Tc1/mariner*) contained over 3,000 transposases each (Figure 1, Table S1).

**Figure 1.**
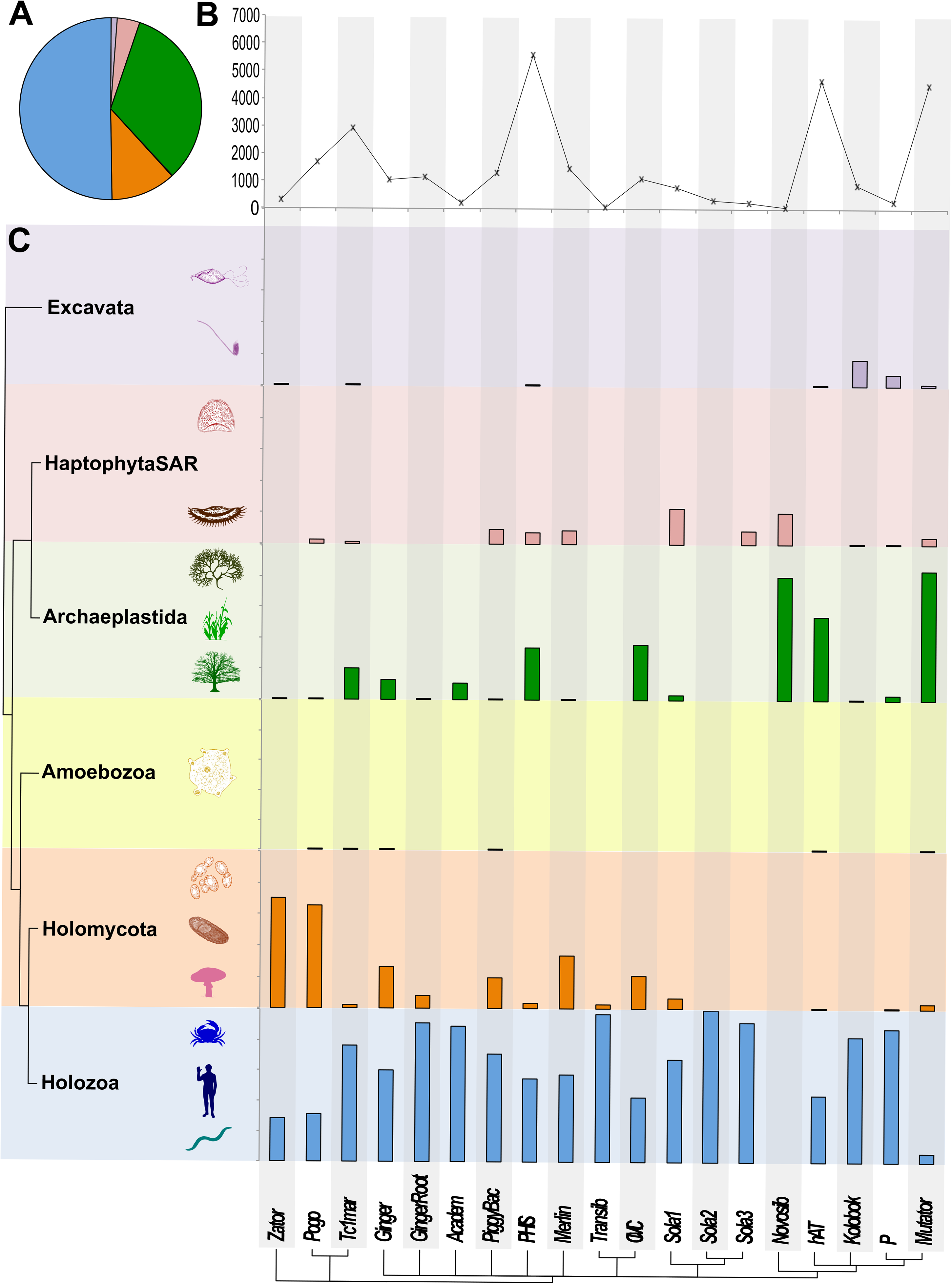
Size and host distribution of DDE transposon superfamilies. **A** Pie chart illustrating the proportion of elements recovered from each major group of eukaryotic diversity. **B** Line graph illustrating the variation in the number of elements recovered for each DDE superfamily. **C** The proportion of transposases from each DDE superfamily recovered from each major group of eukaryotic diversity (each vertical adds up to 100%). DDE superfamily phylogeny adapted from Yuan & Wessler (2011).

DDE transposons share a common evolutionary origin and are considered a deeply ancient group (14, 15). However, relationships among superfamilies and their respective dates of origin are unresolved (13). Little is known about the rate of DDE transposase sequence evolution, but if relatively constant over time, the substantial variation we observed in tree height (Table S2), suggests that DDE superfamilies may differ considerably in age. Yet, superfamily age alone is unlikely to underly observed differences in DDE transposon superfamily size; tree height was not correlated with superfamily size (Spearman’s rank correlation: *R* = 0.16, *p* = 0.517), and it is evident that superfamilies can be deeply ancient, but still contain relatively few transposases. For example, *Recombination-activating gene 1* (*RAG1*) is a key component of the vertebrate adaptive immune system which originated from a *Transib* element that underwent molecular domestication in an ancestor of jawed vertebrates (34). Since the earliest definitive fossils of jawed fishes date to the Silurian period (419-444 million years ago), the *Transib* superfamily is at least 400 million years old (35). Yet, excluding domesticated *RAG1*-like elements, we recovered a total of just 35 *Transib* transposases (all with open reading frames, Figure S18). Thus, rather than reflecting the age of DDE superfamilies, the considerable variation observed in superfamily size suggests that relative rates of transposon proliferation and extinction differ considerably among DDE transposon superfamilies.

### Host range of DDE transposons

At higher levels of host diversity, we find that DDE transposon superfamilies are predominantly associated with hosts from just one or two major eukaryotic divisions (Figure 1C): Holozoa is the primary host for 14 DDE transposon superfamilies and is a major host for almost all superfamilies; Archaeplastida (kingdom Plantae *sensu lato*) is the primary host for three superfamilies (*hAT, Mutator, Novosib*); while Holomycota (fungi and their closest relatives) is the primary host group for just two superfamilies (*Pogo, Zator*) (Figure 1C). These findings suggest that animals are by far the single most important host group for DDE transposon superfamilies collectively, and individually in the majority of cases, while plants and fungi are important hosts for fewer DDE superfamilies. In comparison, haptophytes, stramenopiles, alveolates, and rhizarians (Haptophyta-SAR), excavates, and amoeba, are minor hosts of limited overall importance. Collectively, the number of transposases recovered from each major division of eukarotic host taxonomy correlates with the number of described species each host division contains (Spearman’s rank correlation: *R* = 0.94, *p* = 0.017, Table S3). Thus, our observations deviate considerably from the expected pattern if all DDE transposons were highly generalist and equally able to exploit hosts from across eukaryotic diversity, whereby each superfamily would be display a host range in proportion to the diversity of available hosts (i.e. 50% Holozoa, 11% Archaeplastida, 3% Holomycota, 1% HaptophytaSAR, <1% Ameobozoa and Excavata).

A finer-scale examination of host range also revealed consistent patterns among DDE transposon superfamilies, with a particular diversity and abundance of transposases recovered from deuterostome (chordates, hemichordates, echinoderms), arthropod, and cnidarian genomes (Figure 2). In contrast, dicotyledonous plants, and fungi from Mucoromycota and Ascomycota, were hotspots of abundance for fewer DDE transposon superfamilies (Figure 2). Again, these patterns demonstrate that DDE transposon superfamilies show considerable overlap in their primary hosts overall, but there is strong evidence for differenes in host specialization for certain DDE transposon superfamilies.

**Figure 2.**
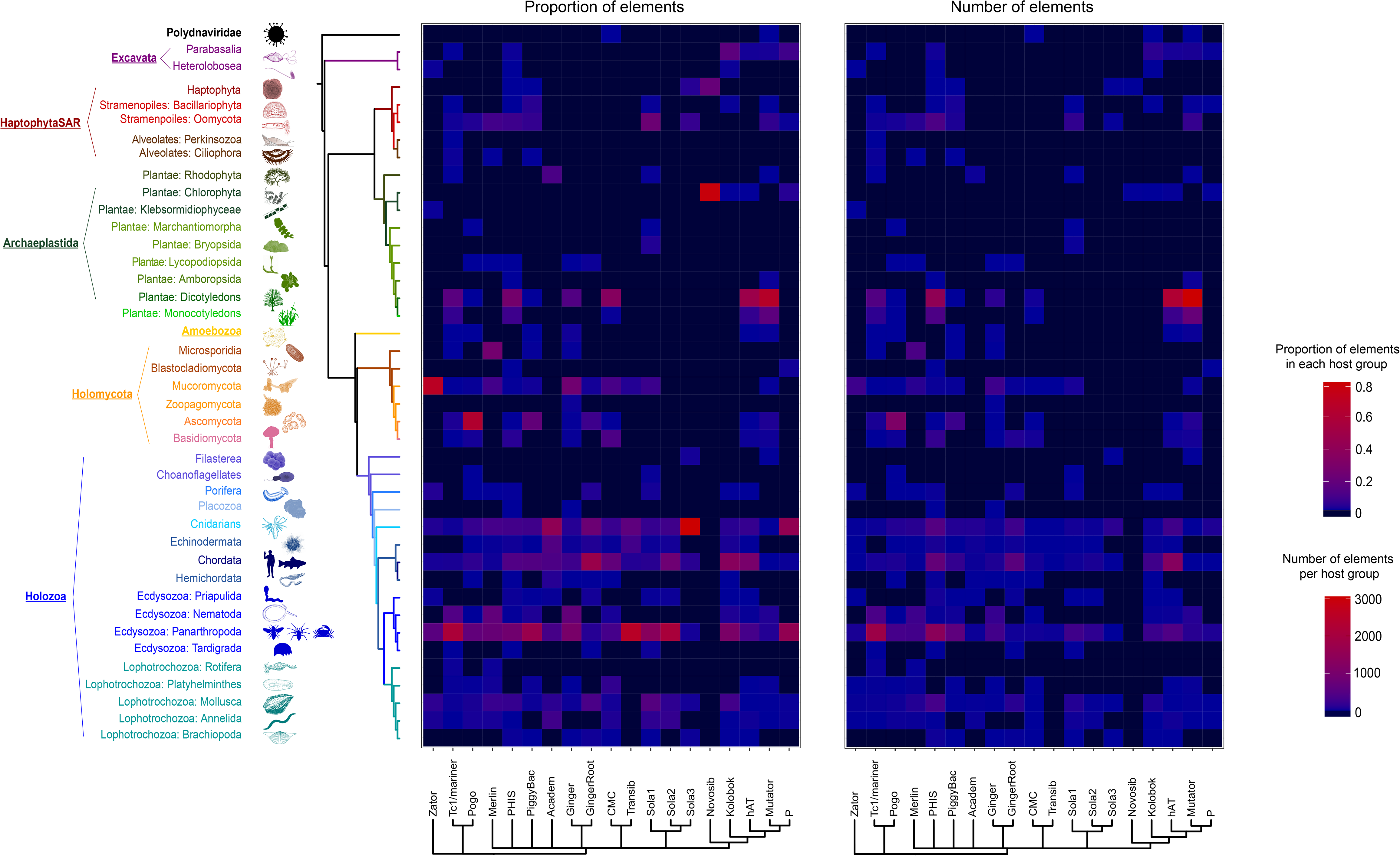
Heatmaps illustrating the proportion of elements (left) and number of elements (right) recovered at a lower level of host eukaryotic diversity for each DDE superfamily. DDE superfamily phylogeny adapted from Yuan & Wessler (2011).

Host range was strongly positively correlated with DDE transposon superfamily size at all levels of host taxonomic rank from kingdom to family (*P* < 0.01, *R^2^* = 50.3%, excluding *Transib* and *Novosib* which contain very few elements, Figures 3A and 3B, Table S4). No superfamily was found to be more generalist than its size suggests (which would be represented by outliers to the right-hand side of the trendlines in Figure 3A). The only strong outlier was the *Mutator* superfamily at the family and order levels of host taxonomy, and the direction was towards greater specificity rather than generalism (Figure 3A).

**Figure 3.**
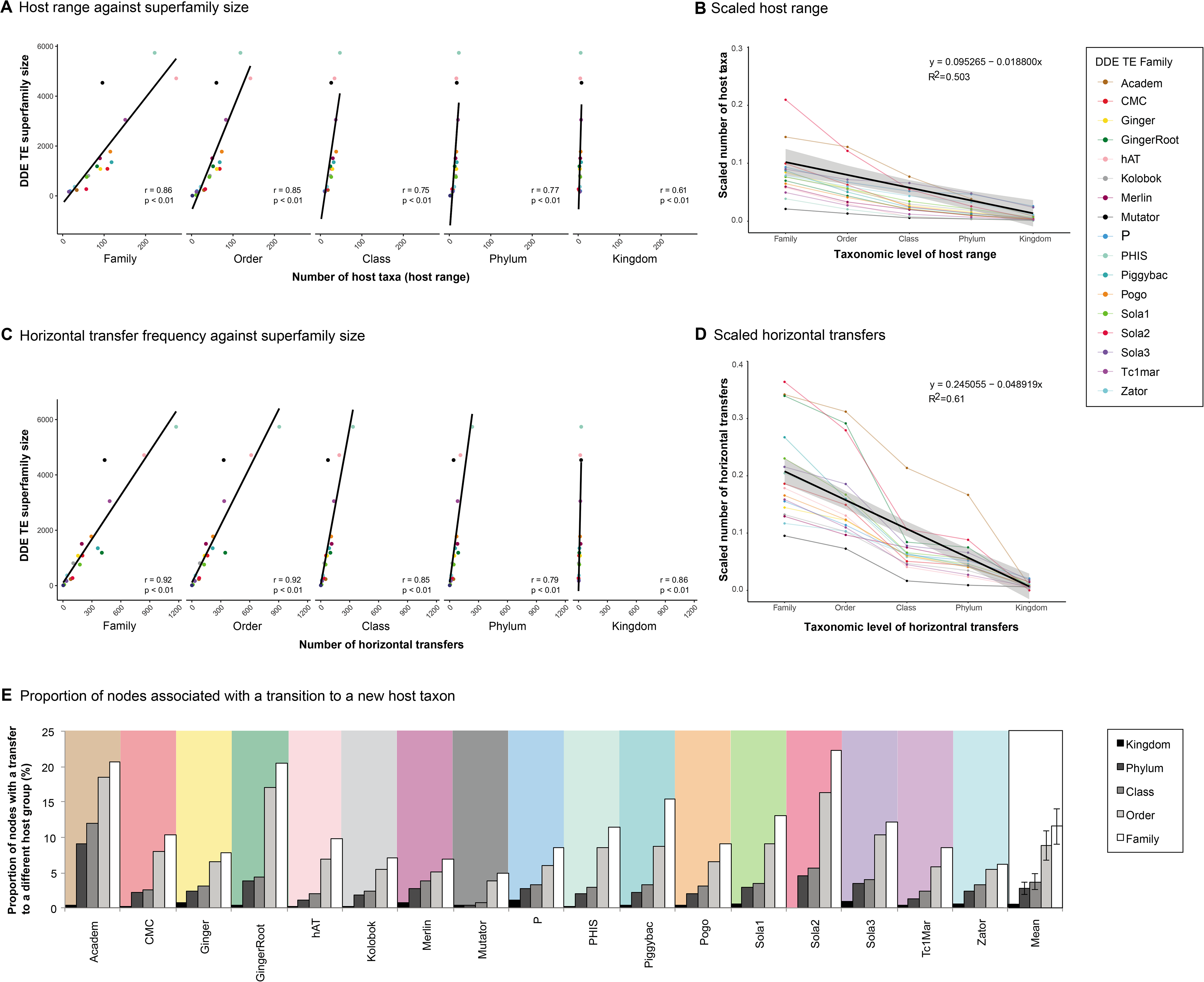
Host range and horizontal transfer of DDE transposons. **A** Scatter plots illustrating host range defined as number of host taxa, for each level of host diversity from family to kingdom. Individual DDE superfamilies are plotted in different colours, and a line of best fit is illustrated at each taxonomic level. **B** Functions summarising host range at each level of host diversity for each DDE superfamily, with a line of best fit. **C** Scatter plots illustrating the estimated frequency of horizontal transfers in numbers of taxa, for each level of host diversity from family to kingdom. Inidividual DDE superfamilies are plotted in different colours, with a line of best fit for each taxonomic level. **D** Functions summarising horizontal transfer at each level of host diversity for all DDE superfamilies with a line of best fit. **E** Bar chart illustrating the proportion of phylogenetic nodes associated with a transition to a new host within each DDE transposon superfamily phylogeny.

Collectively, observations for host range argue against an innate propensity of individual DDE transposon lineages for host generalism across all divisions of eukaryotic taxonomic diversity, instead supporting the existence of barriers that constrain exploitation of very different host environments. However, the nature of the relationship between superfamily size and host range remains unclear. On one hand, variability in host range among superfamilies may drive differences in DDE superfamily size, with more generalist lineages becoming more successful. Conversely, larger superfamilies may have a wider host range because their size increases the likelihood of exposure to a greater diversity of hosts. Thus, to examine the factors responsible for driving host associations in DDE transposons further, we next considered the pattern of horizontal transfer events across host diversity.

### Horizontal transfer of DDE transposons

Variation in the frequency of horizontal transfer among DDE transposon superfamilies would imply intrinsic differences in the capacity for host generalism. To estimate the frequency of higher-level horizontal transfer events (i.e. host family and above) for each DDE transposon superfamily, we applied a phylogeny-based approach previously applied to retroviruses and endogenous retroviruses (24). This assigns host associations to nodes from tip to root within transposase phylogeny, and counts the frequency of transitions in host association between nodes, from root to tip (24). We found a significant correlation between the estimated frequency of horizontal transfer events and superfamily size across all host taxonomic levels (*P* < 0.01, *R^2^* = 61%, excluding *Transib* and *Novosib*, Figure 3C). Thus, the largest DDE transposon superfamilies identified, *PHIS*, *hAT, Tc1/mariner* and *Mutator,* had particularly high numbers of estimated horizontal transfer events. The strong positive correlation identified between DDE superfamily size and frequency of horizontral transfer suggests that superfamilies share a similar propensity for horizontal transfer. Superfamilies with an increased tendency for horizontral transfer would be represented by outliers to the right hand side of the lines of best fit in Figure 3C, indicating a greater number of transfer events than their size suggested relative to other superfamilies, and no such outliers were apparent. Thus, our findings argue against: (i) the existence of adaptations that predispose spread in certain DDE transposon superfamilies, and, (ii) intrinsic differences in generalism being responsible for driving variation in DDE superfamily size, since all superfamilies show a similar propensity for horizontal transfer across eukaryotic host diversity.

Although the *hAT* and *Tc1/mariner* superfamilies are well known to include many confirmed cases of horizontral transfer, our results suggest that the frequency of horizontral transfer events for the *Mutator* and *PHIS* superfamilies are currently underappreciated. For example, out of 211 records of confirmed horizontal transfer events from the ‘Horizontally transferred transposable elements database’ (36), 88 records are for *Tc1/mariner* and 53 are for *hAT*, while just 5 records are for *PHIS* and only 1 is for *Mutator*.

Estimated rates of horizontal transfer decreased sharply with increasing levels of host taxonomy (e.g. mean number of horizontal transfer events per DDE transposon superfamily: family-level 301, order-level 227, class-level 87, phylum-level 61, kingdom-level 11; excluding *Transib* and *Novosib*, which contain very few sequences, Figures 3C and 3D), suggesting that horizontal transfer events across higher-levels of host diversity are rare. For example, assuming a date of origin for DDE transposon superfamilies of 400 million years (my) ago (the minimum age estimate for *Transib,* see above), kingdom-level horizontal transfer events have occurred at an approximate rate of 400my / 11.4 transfers (the mean number of estimated kingdom-level transfers, Table S5), equivalent to one kingdom-level horizontal transfer event every 35.1my. In contrast, transfer across lower levels of host taxonomy are much more frequent (Figure 3C). For example, applying the same parameters suggests that family-level transfers occur at a rate of 400my / 300.6 transfers (Table S5), equivalent to one family-level transfer every 1.33my. Host genomic sampling remains patchy at lower taxonomic levels (e.g. family-level and below), and inferrences of the number of host switches at lower taxonomic levels are likely to be underestimates. However, it is also likely that most DDE transposon superfamilies are much older than 400my, and rates of horizontal transfer may be considerably lower than estimated, particularly for higher levels of host taxonomy.

We next considered the proportion of phylogenetic nodes associated with a transition to a new host taxon, verses nodes where host associations remained constant within DDE transposon superfamily phylogenies. A large proportion of transitions to new host taxa, relative to nodes where associations remained constant, would imply few or no barriers to horizontal transfer at that level of host taxonomy. In contrast, we identify a tendency for host conservatism, with a relatively low proportion of transitions, suggesting that barriers exist which act to constrain horizontal transfer (Figure 3E). The frequency of host transitions decreases with increasing host taxonomic rank (mean proportion of transitions to a new host taxon: family 11.5%, order 8.7%, class 3.7%, phylum 2.8%, kingdom 0.4%, Figure 3E).

Phylogenetic clustering in host relationsips is also evident from direct consideration of DDE transposon superfamily phylogenies, where large subclades are frequently associated with a particular host group (Figure 4, Figures S20-S38). For example, the phylogenies of *CMC, Ginger, hAT, Mutator, PHIS,* and *Tc1/mariner* all contain very large clades exclusively or predominantly composed of transposases recovered from dicotyledonous and monocotyledonous plant hosts. The phylogenetic distance between transposase clusters from plant hosts and other hosts is particularly apparent from principle component analysis (PCA) plots, where plant-host clusters are typically isolated at the extreme of one or both principle component axes, and similar patterns are also apparent for other host groups (Figures S39-S57).

**Figure 4.**
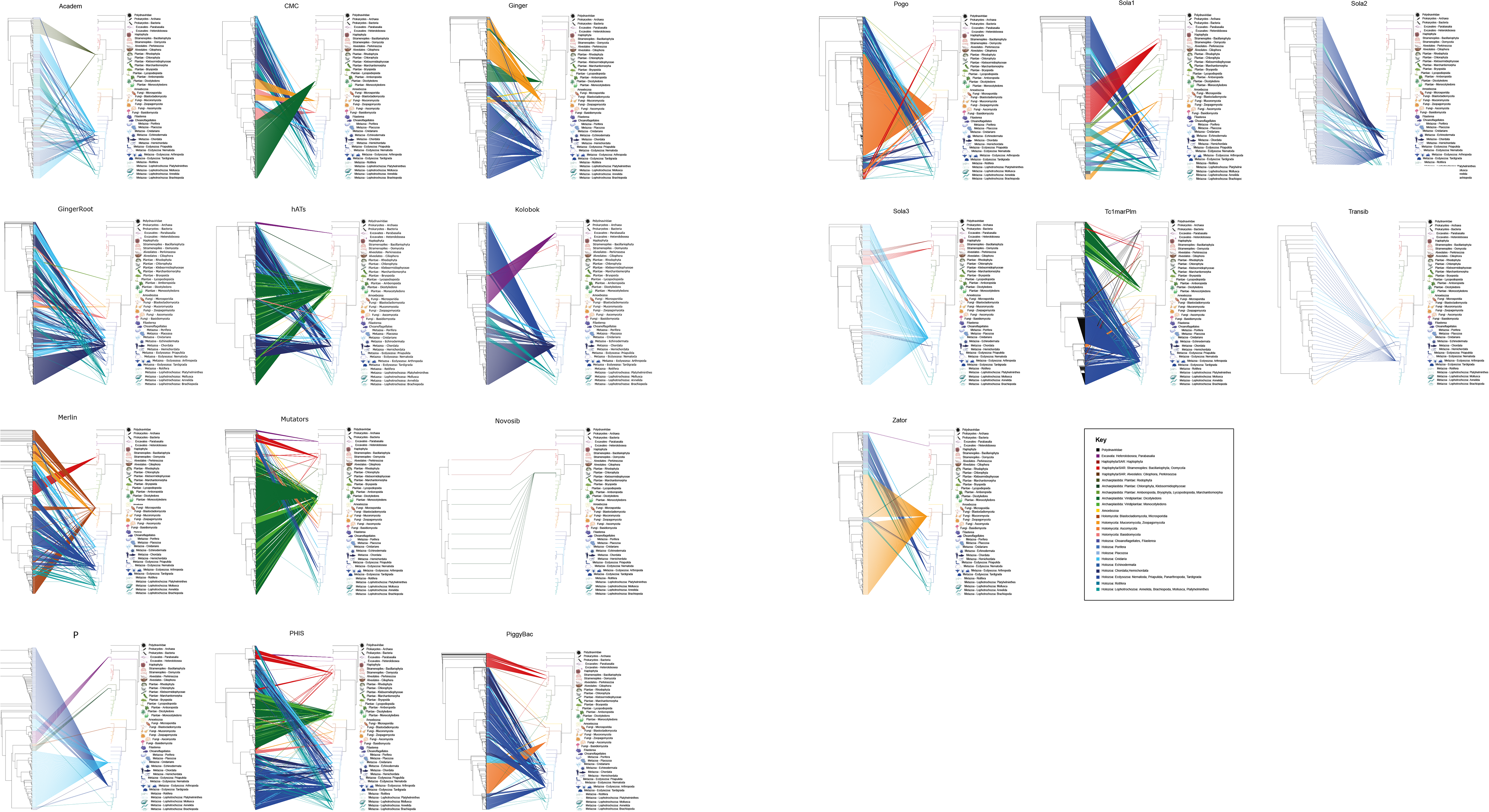
Host-transposon tanglegrams for all 19 DDE superfamilies, illustrating incongruence between higher levels of host eukaryotic phylogeny and DDE superfamily phylogeny.

Our findings for horizontal transfer are consistent with the ‘phylogenetic distance effect’ reported for plant and insect transposons, whereby the number of horizontally acquired transposons shared between host species is negatively correlated with genetic distance between hosts (20). Overall, our findings suggest a pattern of host conservatism for DDE transposons, due to host-associated barriers to horizontal transfer, which increase in strength as the relationship between hosts becomes more distant.

### Host-transposon cophylogenetic congruence

Across biological diversity, tests of host-symbiont cophylogeny have revealed a general pattern of phylogenetic congruence, although this is typically weaker for parasites than for mutualists (37). To examine host-transposon phylogenetic congruence, we generated tanglegrams for all DDE transposon superfamilies and their higher-level eukaryotic host taxa (Figures S20-S38), and tested their fit quantitatively by calculating cophylogenetic signal (38) (Figure 4). If lineages within DDE transposon superfamilies display fidelity to the same host groups (or closely related sets of hosts), a strong signature of host-transposon phylogenetic congruence is expected. Conversely, if horizontal transfer to distantly related hosts is an important feature of DDE transposon evolution, phylogenetic incongruence is expected. We found that phylogenetic patterns between hosts and transposons were incongruent (Figure 4), with no evidence of cophylogenetic signal for any DDE transposon superfamily (Figure 5A). These findings indicate a pervasive influence of horizontal transfer during DDE transposon superfamily evolution across higher levels of host diversity. Thus, while horizontal transfer events are comparatively rare (particularly at higher levels of host taxonomy, see above), they occassionally result in the establishment of successful new lineages of elements, and crucially, the proliferation and persistence of these lineages has been sufficient to disrupt host-transposon cophylogeny over the course of DDE transposon evolutionary history.

**Figure 5.**
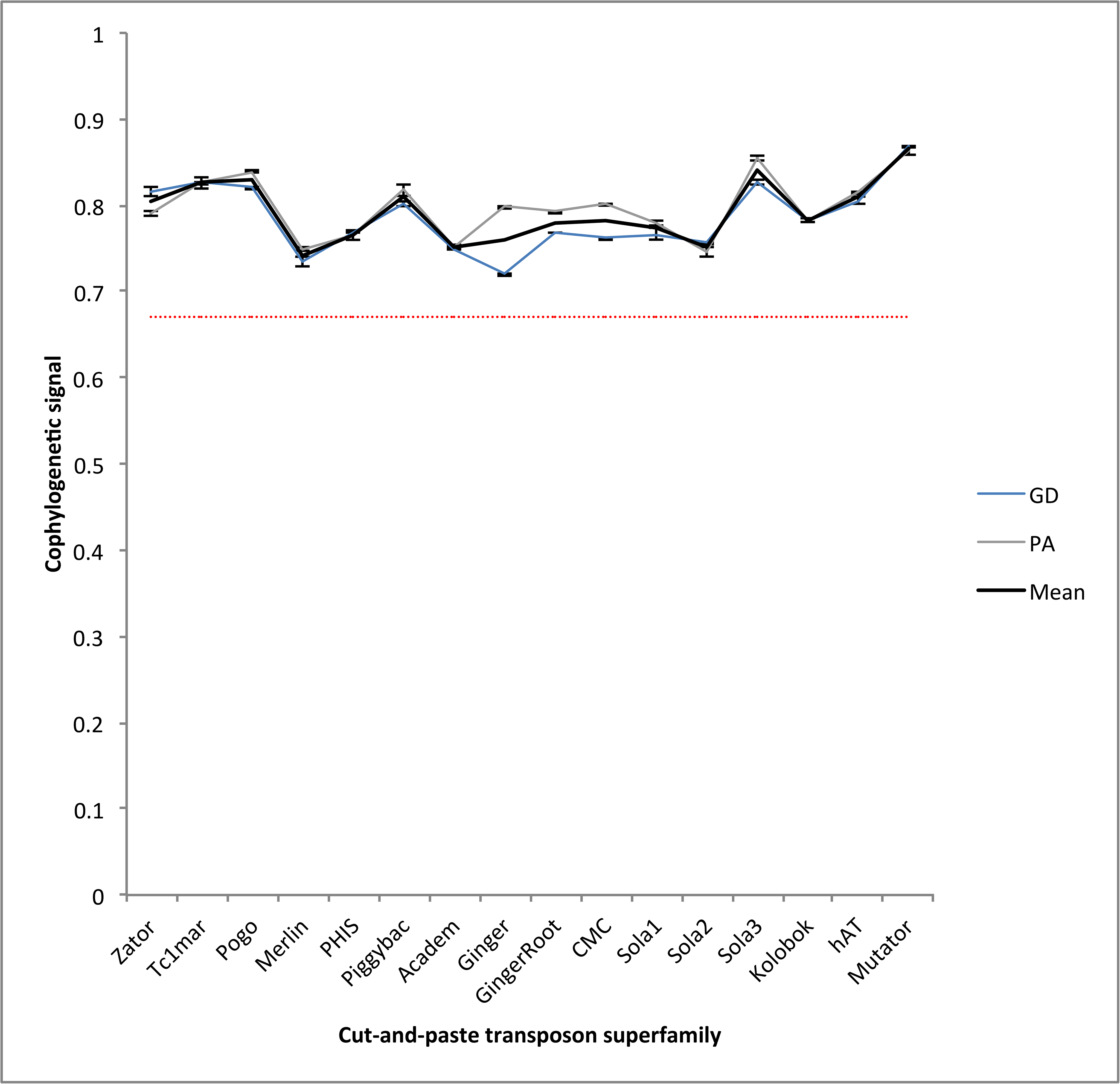
Line graph summarising the lack of cophylogenetic signal between host and transposon phylogeny for all 19 DDE superfamilies. Cophylogenetic signal was quantified as the normalized Gini coefficient (*G\**) computed with both Geodesic Distances (GD) and Procrustes Approach to Cophylogeny (PA) in the program RandomTapas. Cophylogenetic signal is implicated when the normalized Gini coefficient is below 0.67.

### DDE transposon superfamilies and prokaryotic IS elements

The catalytic triad of eukaryotic DDE transposons is shared with prokaryotic insertion sequences (IS elements), and the integrases of LTR elements and retroviruses (32). Furthermore, similarities in sequence identity, protein topology, and transpositional mechanisms between the DDE domains of certain DDE transposons and prokaryotic IS elements are widely recognised (e.g. *Tc1/mariner* / IS630, *Mutator* / IS256, *PiggyBac* / IS1380, *PHIS* / IS5, *Merlin* / IS1016) (2, 28–33). Across all searches, we recovered just 65 DDE transposases originating from bacteria according to Genbank and RefSeq taxonomic descriptions. Considering the vast diversity of prokaryotes and their associated IS elements, this extremely low number suggested that recovered prokaryotic hits were spurious, and that authentic IS elements are too distantly related to DDE transposons to be recovered by our searches. Indeed, multiple independent lines of evidence demonstrated that the identity of these sequences could not be confirmed (see Methods), and we excluded them from analyses. The lack of conclusive hits to prokaryotic DDE transposons suggests that IS elements are not closely related to eukaryotic DDE transposons. Thus, we concur with previous suggestions that while several prokaryotic IS families may share origins with eukaryotic DDE transposon superfamilies, divergence events presumably occurred very anciently, potentially even prior to the split of eukaryotes and prokaryotes (2, 28).

Although we found no conclusive evidence of horizontal transfer events of DDE transposons to prokaryotes, occasional transfers of prokaryotic IS elements to eukaryotes are documented in the literature, suggesting that transfers in this direction are more permissive. These include the repeated transfer of IS607 family elements to the genomes of several eukaryotic hosts (39), and an IS5-like element to a bdelloid rotifer (40). However, the long-term success of elements that undergo such transfers is currently unclear.

## Conclusions

We identify considerable variation in size and phylogeny among DDE transposon superfamilies, with just four superfamilies accounting for the majority of transposases recovered. Considerable overlap in host range is revealed, with most superfamilies having a primary distribution in animal genomes, and just a few superfamilies specialising in plant or fungus genomes. Host range and horizontal transfer are both strongly positively correlated with DDE transposon superfamily size, implying a similar capacity for spread. Overall, DDE transposons show a dominant pattern of host conservatism, with horizontal transfer events being comparatively rare, particularly at higher levels of host taxonomy, but of sufficient frequency and magnitude to disrupt any signal of host-transposon cophylogeny.

In common with host-parasite interactions more broadly, host associations in DDE transposons are presumably shaped by a combination of opportunity filters (likelihood of encountering a new host) and compatibility filters (ability to survive and reproduce in a new host) (*sensu* 41). Regarding opportunity filters, DDE transposons lack the envelope/envelope-like glycoprotein genes encoded by viruses and certain LTR Transposons (42–44) and are not directly infectious. However, naked DNA remains viable in a wide range of animal fluids (45), and the intimate ecological relationships shared among diverse hosts presumably offer plentiful opportunities for the cross-taxon spread of DDE transposons. Indeed, there is strong evidence that host-symbiont interactions mediate the horizontal transfer of DDE transposons, including across large taxonomic divides, such as from invertebrate to vertebrate hosts (46–48). Horizontal transfer via viral vectors is particularly noteworthy, given that viruses inject their genome directly into host cells, and evidence of viral-assisted horizontal transfer of mobile DNA is accumulating (49, 50). Our findings suggest that DDE transposons from different superfamilies are vectored with similar efficiency, and are consistent with a hypothesis of DDE transposons as passive passengers during host spread.

Regarding compatibility filters, estimates of horizontal transfer and tests of host-transposon cophylogeny both suggest that DDE transposons posses an underlying capacity to cross divides between distantly related host taxa. These findings are in line with empirical studies demonstrating that some DDE transposons can function readily when experimentally transferred to distantly related hosts, including those from different host phyla and kingdoms, and the observation that transposases can be the only proteins required for successful transposition (39, 40, 51, 52). However, across DDE transposon diversity, we find that horizontal transfer events occur much less frequently as the taxonomic distance between hosts increases. Meanwhile frequencies of transfer events across DDE transposon superfamily phylogeny reveal a dominant overall pattern of host conservatism, suggesting that DDE transposon lineages often remain associated with taxonomically similar hosts for protracted periods of time, punctuacted by occasional successful horizontal transfer events. Thus, our results imply that trade-offs typically act to constrain the ability of DDE transposons to undergo horizontal transfer and successful proliferation in distantly related hosts.

Patterns for host range reinforce those identified for horizontal transfer, demonstrating the overall dominance of animal hosts, with only certain DDE transposon lineages displaying an ability to successfully exploit plant and fungal hosts. For example, while most DDE transposon superfamilies contain very few or no transposases recovered from plant genomes, the four largest superfamilies (*PHIS*, *hAT, Mutator, Tc1/mariner*), all contain major clades of transposases recovered almost exclusively from plant hosts. Consequently, rare shifts to plant hosts in certain DDE transposon lineages appear to have led to highly successful radiations of elements, leading to the diversification of large clades of plant-associated transposases, which at least partially explain the relative success of these superfamilies. Therefore, while our results support a passive mode of spread for DDE transposons, they are not fully inkeeping with suggestions that DDE transposons are genetically streamlined opportunists (14, 16, 20). Instead, the patterns described above suggest that a process of host adaptation is important in determining the host relationships of individual DDE transposon lineages.

Adaptations to novel distinct cellular environments could arise via several mechanisms. For example, DDE transposons can contain ORFs in addition to the transposase (e.g. 53), while considerable variation exists among and within DDE transposon superfamilies in transposase structure (e.g. spacing of the DDE motif), transposase target site preference, presence and type of promoter, and mode of replicative transposition (13, 26, 53–55). Detailed surveys of such variation are required across DDE transposon diversity to investigate this issue further. Of particular interest are the processes that occur following horizontral transfer between distantly related host taxa. For example, to investigate if adaptations to the new host are gradually evolved over successful waves of proliferation, or if horizontral transfer most frequently involves ‘hopeful monsters’, in which genes for host adaptation have become mutated or lost, potentially allowing a lower efficiency of proliferation across a wider set of hosts intially.

We identify considerable differences in DDE transposon superfamily size, and show that these are not a consequence of differences in the propensity for host spread among superfamilies. Consequently, elucidating the factors responsible for driving variation in DDE superfamily size, and the extent to which this fluctuates over time, represent major questions for future research. Transposon activity rates are associated with host virulence, and in response, host genomes have evolved adaptations to repress transposon proliferation (5, 14). In turn, this may put pressure on transposons to evolve counter-adaptations (56). Such antagonistic interactions could lead to the fluctuating success of DDE transposon lineages that share targets of host repression. However, while there is relatively little evidence of transposons having successfully evolved mechanisms to escape host silencing, some DDE transposons, such as certain *Tc1/mariner* elements, have evolved mechanisms to self-regulate their own proliferation (56). This presents the intriguing possibility that the most prolific DDE transposon lineages, such as *Tc1/mariner* elements, may be those that have evolved optimal self-regulation strategies, to balance the benefits of proliferation against costs to host fitness, rather than those which utilise maximal profliferation strategies, geared towards leaving as many copies as possible over short timeframes. Such ‘optimal proliferation’ strategies may be most effective when coupled with an insertion site preference for genomic safe havens, to further limit negative impacts on host fitness, and elements that manage proliferation optimally may leave more descent copies. Proliferation rates are known to vary greatly among DDE transposons (52), and according to host genomic background (53), and thus a variety of context-dependent genetic adaptations may underlie proliferation patterns in DDE transposons.

As DDE transposons proliferate, non-autonomous elements accumulate in the host genome, limiting the proliferation of autonomous elements, and eventually driving functional copies extinct (26). The observation that DDE transposons are now extinct in the human genome, but are represented by a large number of fossil elements, suggests considerable former activity (57). However, the extent to which patterns of fluctuating success among hosts are driven by the accumulation of non-autonomous elements represents a question for future study.

Finally, a major challenge facing DDE transposon research is managing systems to characterise and catalogue the vast numbers of elements continually being discovered. Until addressed more formally, the phylogenetic resource we provide, building on the earlier framework of Yuan and Wessler (13), offers a phylogenetic alternative to utilising sequence similarity thresholds to classify elements, such as BLAST e-value scores below 0.01 (13), or identity thresholds (58). It also offers a basis for further evolutionary analyses, by providing a context to examine phylogenetic relationships among DDE transposons isolated from genomes of interest, or for comparative phylogenetic analyses, such as to correlate genetic variation in DDE transposons with estimates of their host range.

## Materials and Methods

### Mining of DDE transposases

We focussed on the only shared gene among DDE transposons, the transposase domain. We included transposases from all DDE superfamilies in our analyses, updating the 17 superfamily framework of Yuan and Wessler (13), i.e. *Tc1/mariner, Merlin, PIF/Harbinger, MULE* (Mutator + Rehavkus), *P, hAT, Kolobok, Novosib, Sola1, Sola2, Sola3, PiggyBac, CMC* (CACTA, Mirage, Chapaev), *Transib, Academ, Ginger, Zator*, to include a total of 19 superfamilies, with the follow changes in line with recent research: (i) we split *Pogo* from *Tc1/mariner* to form its own distinct superfamily (59, 60) (ii); we grouped *PIF/Harbinger* with *Spy* as the superfamily *PHIS* (61); (iii) we included the newly described *GingerRoot* family (62).

We used the transposase amino acid sequences from Yuan and Wessler (13) for our database searches, except for *MULE, Pogo, Tc1/mariner*, and *GingerRoot,* where we used sequences from more recent analyses (53, 54, 62). We queried the NCBI nr database with BLASTp (63, 64), using a script to parse outputs, specifying table format output, including a column containing the subject sequence. Matches were filtered to retain sequences with >50% identity to the query over a minimum of 50% of query length. Fasta files were generated using the filtered BLAST tables. Matches were extracted and processed into fasta format using awk and sed, and the EMBOSS tool v6.6 (65). Headers were constructed using the following format: “>[query_seqid]—[subject_seqid]#[subject_taxid]”. Thus, each match was identified by its reference numbers and taxonomic origin identity (‘taxid’) (66, 67). Nucleotide queries such as TBLASTN were not performed, since the searches were computationally prohibitive given the large number of query sequences and database size, while BLASTN searches would require computationally prohibitive downstream reconstruction of transposase open-reading frames. The vast majority of mined transposases recovered were uninterrupted by stop codons (97.7%, Table S7).

Sequences for each superfamily were aligned in MAFFT v7 (68) using the --auto flag, and any duplicate sequences originating from the same species were removed. Alignments were checked to confirm that transposases were present in one superfamily only, using a script to cross-reference fasta headers, which confirmed this to be the case. While all transposases were confirmed as occurring in one superfamily only, we acknowledge that the evolution of *Ginger* and *GingerRoot* transposases is complex, and their relationships with the integrases of certain Gypsy LTR retrotransposons with which they share similarity, remains unclear (62, 69). Phylogeny was estimated using FastTree v2.1.11 (70), applying minimum-evolution subtree-pruning-regrafting (SPRs) and maximum-likelihood nearest-neighbour interchanges (NNIs). We performed 1,000 bootstrap repetitions, specifying the -spr 4 option to improve SPRs, and the ‘–mlacc 2’ and ‘-slownni’ options to increase accuracy. Following this, sequences representing the diversity of each superfamily were selected using a previously described phylogenetic taxon reduction protocol (24), and used to perform a second iteration of queries to retrieve additional sequences following the protocol described above.

### Phylogenetic analysis

To generate final phylogenies for each superfamily, we performed 1,000 ultra-fast bootstrap repetitions in IQ-TREE v1.6.12 (71), specifying the best fit amino acid model for each family as indicated by ModelFinder (72). Sequences from related transposon superfamilies were used as outgroups. Transposon sequences co-opted for host purposes are subject to host evolutionary patterns and processes rather than transposon evolutionary dynamics. Consequently, we removed domesticated elements from our phylogenies as identified with reference to the literature (Dataset S39: https://doi.org/10.6084/m9.figshare.14899350; https://figshare.com/s/f88085e8034904f648a0). Phylogenetic trees were visualised and rooted in FigTree v1.4.4 (http://tree.bio.ed.ac.uk/software/figtree/).

All DDE transposon superfamilies were strongly supported as monophyletic, with clade support values for the ancestral node in each superfamily between 98-100%, with the exception of *Zator* (73% support). Tree height, as measured in substitutions per site (averaged over all root-to-tip path lengths) was calculated using the function ‘calc_tree_height’ in the Perl package Bio::Phylo (73). ClusterPicker v1.2.3 (74) was used to generate clusters of sequences using a genetic distance threshold increment of 0.5%, from 1% - 50%. At each threshold the number of clades (i.e. ≥2 sequences) identified were summed with singleton sequences, to calculate the total number of discrete lineages identified at each genetic diversity cut-off. Results were analysed in R using tidyverse and ggplot2 (75, 76), illustrating a decrease in clusters with increasing cut-off threshold. Polynomial models were fitted to the data using the lm function in R, and the total area under the curve (AUC) was calculated using the ‘area_under_curve’ function in bayestestR (77). A 95% AUC value provides a threshold whereby the number of phylogenetic clusters inferred with increasing genetic distance plateaus in most cases for DDE transposon superfamilies, representing a standardised approach to inferring taxa across superfamilies. The 95% AUC genetic distance cut-off value was determined by iteratively adding the AUC for each data point, and used to divide sequences within each phylogeny into sets of clades, which are annotated for each superfamily phylogeny in FigTree format (Datasets S20-S38).

A eukaryote host tree was generated in ETE3 (78) with reference to recent phylogenetic analyses (79–81). Host-transposon tanglegrams were generated in the R package Ape v5.4.1 (82), and cophylogenetic signal was measured using the normalized Gini coefficient (*G\**), computed with both geodesic distances (GD)(83) and procrustes approach to cophylogeny (PA)(84) in RandomTapas (38). A threshold of 0.67 was applied to evaluate if cophylogenetic signal was higher or lower than expected by chance (38). Heatmaps were generated using ggplot2 v3.3.2 in Rstudio. The frequency of higher-level horizontal transfer events and quantification of host transitions for each DDE TE superfamily were estimated using a phylogeny-based approach previously applied for retroviruses, involving reconstruction of ancestral hosts based on contemporary host associations (24). PCA was performed using branch length pairwise distances, calculated with the ‘cophenetic.phylo’ command in Ape (82). Classical multidimensional scaling (MDS) was performed on each distance matrix using the ‘cmdscale’ command of the statistical system package in R, where a maximum of 2 dimensions was specified (*k* = 2). PCA plots were generated using ggplot2 (v3.3.2).

### Tests for the validity of prokaryotic sequences

To investigate the validity of the small number of transposase hits recovered that were labelled as originating from prokaryotic hosts, we considered several lines of evidence. Firstly, we examined their flanking sequences for evidence of TIRs bounded by genomic DNA from their labelled host species. In the vast majority of cases, we found no evidence even of the existence of TIRs (Table S6). In several cases where putative TIRs were identified, full transposon sequences did not match their labelled taxon in BLASTP or BLASTN searches, but instead matched DDE transposons from eukaryotic taxa or different prokaryotic species. No searches recovered a top hit for the prokaryotic species that the transposase was labelled as originating from. Additionally, transposases labelled as being of prokaryotic origin typically came from samples where contamination is more likely (environmental samples, metagenomes, bacterial symbionts of eukaryotic hosts). Meanwhile, previous phylogenetic analyses of individual DDE transposon superfamilies that included prokaryotic IS elements recovered eukaryotic elements as monophyletic to the exclusion of IS elements (52, 85–87), while in our analyses, transposases labelled as prokaryotic occurred within the ingroups of DDE transposon superfamilies.

## Acknowledgments

We thank Professor Edze Westra and Professor Angus Buckling for providing comments on the manuscript. MD and AH were supported by a Biotechnology and Biological Sciences Research Council (BBSRC) David Phillips Fellowship (BB/N020146/1) to AH. TB was supported by a studentship from the Biotechnology and Biological Sciences Research Council-funded South West Biosciences Doctoral Training Partnership (BB/M009122/1).

## Data Availability

Amino acid alignments are available at: doi.org/10.6084/m9.figshare.14899335 (https://figshare.com/s/ee047e33109c36d80db9). Accompanying annotated phylogenies are available at: doi.org/10.6084/m9.figshare.14899344; (https://figshare.com/s/9f9efee7c3227f4ac88e). Updated versions of alignments and phylogenies are maintained at: www.jumpinggenes.org/data/.

## Notes

https://figshare.com/s/ee047e33109c36d80db9

https://figshare.com/s/9f9efee7c3227f4ac88e

### Competing Interest Statement

The authors have declared no competing interest.

