## Supplementary figures and tables for "Broadscale evolutionary analysis of eukaryotic DDE transposons"

**Figures S1-S19.** DDE transposon superfamily phylogenies.

**Figures S20-S38.** DDE transposon-host cophylogenies.

**Figures S39-S57.** PCA plots of phylogenetic distance illustrating patterns in host clustering.

**Table S1.** Number of transposases recovered for each DDE TE superfamily.

**Table S2.** Mean tree height for each DDE TE superfamily.

**Table S3.** Species diversity of major host groups and corresponding numbers of transposases recovered

**Table S4.** Host range at different taxonomic levels for each DDE TE superfamily.

**Table S5.** Number of host taxa present at different taxonomic levels for each DDE TE superfamily.

**Table S6.** Number of host switches at different taxonomic levels for each DDE TE superfamily.

**Table S7.** Verification checks of transposases labelled as originating from bacteria.

**Table S8.** Numbers of stop codons present in transposases for each DDE transposon superfamily.

**Datasets S1-S19 ( https://figshare.com/s/ee047e33109c36d80db9 ).** DDE transposon superfamily transposase amino acid alignments.

**Datasets S20-S38 ( https://figshare.com/s/9f9efee7c3227f4ac88e ).** Annotated DDE transposon superfamily phylogenies in FigTree format, organised into inferred taxonomic clusters.

**Dataset S39 ( https://figshare.com/s/f88085e8034904f648a0 ).** Sequence reference numbers for domesticated transposases.

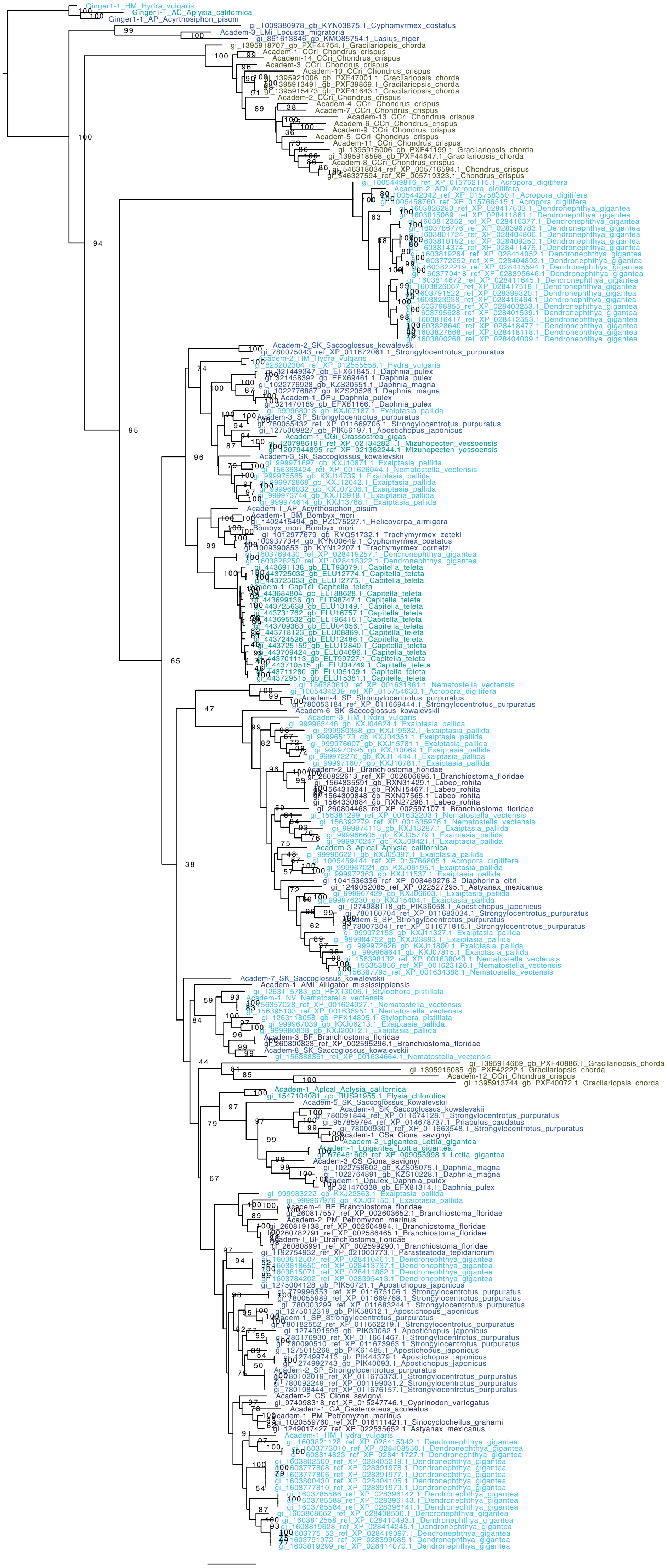
Ginger1-1_HM_Hydra_vulgaris
Ginger1-1_AC_Aplysia_californica
Ginger1-1_AP_Acyrthosiphon_pisum
gi_1009380978_gb_KYN03875.1_Cyphomyrmex_costatus
Academ-3_LMi_Locusta_migratoria
gi_861613846_gb_KMQ85754.1_Lasius_niger
gi_1395918707_gb_PXF44754.1_Gracilariopsis_chorda
Academ-1_CCri_Chondrus_crispus
Academ-14_CCri_Chondrus_crispus
Academ-3_CCri_Chondrus_crispus
Academ-10_CCri_Chondrus_crispus
gi_1395921006_gb_PXF47001.1_Gracilariopsis_chorda
gi_1395913491_gb_PXF39869.1_Gracilariopsis_chorda
gi_1395915473_gb_PXF41643.1_Gracilariopsis_chorda
Academ-2_CCri_Chondrus_crispus
Academ-4_CCri_Chondrus_crispus
Academ-7_CCri_Chondrus_crispus
Academ-13_CCri_Chondrus_crispus
Academ-6_CCri_Chondrus_crispus
Academ-9_CCri_Chondrus_crispus
Academ-5_CCri_Chondrus_crispus
Academ-11_CCri_Chondrus_crispus
gi_1395915006_gb_PXF41199.1_Gracilariopsis_chorda
gi_1395918598_gb_PXF44647.1_Gracilariopsis_chorda
Academ-8_CCri_Chondrus_crispus
gi_546318034_ref_XP_005716594.1_Chondrus_crispus
gi_546327594_ref_XP_005719323.1_Chondrus_crispus
gi_1005449816_ref_XP_015762115.1_Acropora_digitifera
Academ-2_ADi_Acropora_digitifera
gi_1005442042_ref_XP_015758350.1_Acropora_digitifera
gi_1005458760_ref_XP_015766515.1_Acropora_digitifera
gi_1603826280_ref_XP_028417603.1_Dendronephthya_gigantea
gi_1603815069_ref_XP_028411861.1_Dendronephthya_gigantea
gi_1603812352_ref_XP_028410377.1_Dendronephthya_gigantea
gi_1603786776_ref_XP_028396783.1_Dendronephthya_gigantea
gi_1603801724_ref_XP_028404806.1_Dendronephthya_gigantea
gi_1603810192_ref_XP_028409250.1_Dendronephthya_gigantea
gi_1603814374_ref_XP_028411476.1_Dendronephthya_gigantea
gi_1603819264_ref_XP_028414052.1_Dendronephthya_gigantea
gi_1603772252_ref_XP_028404892.1_Dendronephthya_gigantea
gi_1603822219_ref_XP_028415594.1_Dendronephthya_gigantea
gi_1603770418_ref_XP_028395646.1_Dendronephthya_gigantea
gi_1603814672_ref_XP_028411645.1_Dendronephthya_gigantea
gi_1603826067_ref_XP_028417518.1_Dendronephthya_gigantea
gi_1603791522_ref_XP_028399320.1_Dendronephthya_gigantea
gi_1603823938_ref_XP_028416464.1_Dendronephthya_gigantea
gi_1603798855_ref_XP_028403253.1_Dendronephthya_gigantea
gi_1603795626_ref_XP_028401539.1_Dendronephthya_gigantea
gi_1603816417_ref_XP_028412553.1_Dendronephthya_gigantea
gi_1603828640_ref_XP_028418477.1_Dendronephthya_gigantea
gi_1603827668_ref_XP_028418118.1_Dendronephthya_gigantea
gi_1603800268_ref_XP_028404009.1_Dendronephthya_gigantea
Academ-2_SK_Saccoglossus_kowalevskii
gi_780075043_ref_XP_011672061.1_Strongylocentrotus_purpuratus
Academ-2_HM_Hydra_vulgaris
gi_828202304_ref_XP_012555558.1_Hydra_vulgaris
gi_321449347_gb_EFX61845.1_Daphnia_pulex
gi_321458392_gb_EFX69461.1_Daphnia_pulex
gi_1022776928_gb_KZS20551.1_Daphnia_magna
gi_1022776887_gb_KZS20526.1_Daphnia_magna
Academ-1_DPu_Daphnia_pulex
gi_321470189_gb_EFX81166.1_Daphnia_pulex
gi_999968013_gb_KXJ07187.1_Exaiptasia_pallida
Academ-3_SP_Strongylocentrotus_purpuratus
gi_780055432_ref_XP_011669706.1_Strongylocentrotus_purpuratus
gi_1275009827_gb_PIK56197.1_Apostichopus_japonicus
Academ-1_CGi_Crassostrea_gigas
gi_1207986191_ref_XP_021342821.1_Mizuhopecten_yessoensis
gi_1207944895_ref_XP_021362244.1_Mizuhopecten_yessoensis
Academ-3_SK_Saccoglossus_kowalevskii
gi_999971697_gb_KXJ10871.1_Exaiptasia_pallida
gi_156363424_ref_XP_001626044.1_Nematostella_vectensis
gi_999975565_gb_KXJ14739.1_Exaiptasia_pallida
gi_999972868_gb_KXJ12042.1_Exaiptasia_pallida
gi_999968032_gb_KXJ07206.1_Exaiptasia_pallida
gi_999973744_gb_KXJ12918.1_Exaiptasia_pallida
gi_999974614_gb_KXJ13788.1_Exaiptasia_pallida
Academ-1_AP_Acyrthosiphon_pisum
Academ-1_BM_Bombyx_mori
gi_1402415494_gb_PZC75227.1_Helicoverpa_armigera
Bombyx_mori_Bombyx_mori
gi_1012977679_gb_KYQ51732.1_Trachymyrmex_zeteki
gi_1009377344_gb_KYN00649.1_Cyphomyrmex_costatus
gi_1009390853_gb_KYN12207.1_Trachymyrmex_cornetzi
gi_1603769430_ref_XP_028419257.1_Dendronephthya_gigantea
gi_1603828250_ref_XP_028418322.1_Dendronephthya_gigantea
gi_443691138_gb_ELT93079.1_Capitella_teleta
gi_443725032_gb_ELU12774.1_Capitella_teleta
gi_443725033_gb_ELU12775.1_Capitella_teleta
Academ-1_CapTel_Capitella_teleta
gi_443684804_gb_ELT88628.1_Capitella_teleta
gi_443699136_gb_ELT98747.1_Capitella_teleta
gi_443725638_gb_ELU13149.1_Capitella_teleta
gi_443731762_gb_ELU16757.1_Capitella_teleta
gi_443695532_gb_ELT96415.1_Capitella_teleta
gi_443709383_gb_ELU04056.1_Capitella_teleta
gi_443718123_gb_ELU08869.1_Capitella_teleta
gi_443724526_gb_ELU12486.1_Capitella_teleta
gi_443725159_gb_ELU12840.1_Capitella_teleta
gi_443709424_gb_ELU04096.1_Capitella_teleta
gi_443701113_gb_ELT99727.1_Capitella_teleta
gi_443710515_gb_ELU04749.1_Capitella_teleta
gi_443711280_gb_ELU05109.1_Capitella_teleta
gi_443729515_gb_ELU15381.1_Capitella_teleta
gi_156380610_ref_XP_001631861.1_Nematostella_vectensis
gi_1005434239_ref_XP_015754630.1_Acropora_digitifera
Academ-4_SP_Strongylocentrotus_purpuratus
gi_780053184_ref_XP_011669444.1_Strongylocentrotus_purpuratus
Academ-6_SK_Saccoglossus_kowalevskii
Academ-3_HM_Hydra_vulgaris
gi_999965446_gb_KXJ04624.1_Exaiptasia_pallida
gi_999980358_gb_KXJ19532.1_Exaiptasia_pallida
gi_999965173_gb_KXJ04351.1_Exaiptasia_pallida
gi_999976607_gb_KXJ15781.1_Exaiptasia_pallida
gi_999970895_gb_KXJ10069.1_Exaiptasia_pallida
gi_999972270_gb_KXJ11444.1_Exaiptasia_pallida
gi_999971607_gb_KXJ10781.1_Exaiptasia_pallida
Academ-2_BF_Branchiostoma_floridae
gi_260822613_ref_XP_002606696.1_Branchiostoma_floridae
gi_1564335591_gb_RXN31429.1_Labeo_rohita
gi_1564318241_gb_RXN15467.1_Labeo_rohita
gi_1564309848_gb_RXN07565.1_Labeo_rohita
gi_1564330884_gb_RXN27298.1_Labeo_rohita
gi_260804463_ref_XP_002597107.1_Branchiostoma_floridae
gi_156381299_ref_XP_001632203.1_Nematostella_vectensis
gi_156392279_ref_XP_001635976.1_Nematostella_vectensis
gi_999974113_gb_KXJ13287.1_Exaiptasia_pallida
gi_999966605_gb_KXJ05779.1_Exaiptasia_pallida
gi_999970247_gb_KXJ09421.1_Exaiptasia_pallida
Academ-3_Aplcal_Aplysia_californica
gi_999966221_gb_KXJ05397.1_Exaiptasia_pallida
gi_1005459444_ref_XP_015766805.1_Acropora_digitifera
gi_999967021_gb_KXJ06195.1_Exaiptasia_pallida
gi_999972363_gb_KXJ11537.1_Exaiptasia_pallida
gi_1041536336_ref_XP_008469276.2_Diaphorina_citri
gi_1249052085_ref_XP_022527295.1_Astyanax_mexicanus
gi_999967429_gb_KXJ06603.1_Exaiptasia_pallida
gi_999976230_gb_KXJ15404.1_Exaiptasia_pallida
gi_1274988118_gb_PIK36058.1_Apostichopus_japonicus
gi_780160704_ref_XP_011683034.1_Strongylocentrotus_purpuratus
Academ-5_SP_Strongylocentrotus_purpuratus
gi_780073041_ref_XP_011671815.1_Strongylocentrotus_purpuratus
gi_999972153_gb_KXJ11327.1_Exaiptasia_pallida
gi_999984752_gb_KXJ23893.1_Exaiptasia_pallida
gi_999972626_gb_KXJ11800.1_Exaiptasia_pallida
gi_999968641_gb_KXJ07815.1_Exaiptasia_pallida
gi_156398132_ref_XP_001638043.1_Nematostella_vectensis
gi_156353856_ref_XP_001623126.1_Nematostella_vectensis
gi_156387795_ref_XP_001634388.1_Nematostella_vectensis
Academ-7_SK_Saccoglossus_kowalevskii
Academ-1_AMi_Alligator_mississippiensis
gi_1263115783_gb_PFX13006.1_Stylophora_pistillata
Academ-1_NV_Nematostella_vectensis
gi_156357028_ref_XP_001624027.1_Nematostella_vectensis
gi_156395103_ref_XP_001636951.1_Nematostella_vectensis
gi_1263118058_gb_PFX14895.1_Stylophora_pistillata
gi_999967039_gb_KXJ06213.1_Exaiptasia_pallida
gi_999980838_gb_KXJ20012.1_Exaiptasia_pallida
Academ-3_BF_Branchiostoma_floridae
gi_260800823_ref_XP_002595296.1_Branchiostoma_floridae
Academ-8_SK_Saccoglossus_kowalevskii
gi_156388351_ref_XP_001634664.1_Nematostella_vectensis
gi_1395914669_gb_PXF40886.1_Gracilariopsis_chorda
gi_1395916085_gb_PXF42222.1_Gracilariopsis_chorda
Academ-12_CCri_Chondrus_crispus
gi_1395913744_gb_PXF40072.1_Gracilariopsis_chorda
Academ-1_Aplcal_Aplysia_californica
gi_1547104081_gb_RUS91955.1_Elysia_chlorotica
Academ-5_SK_Saccoglossus_kowalevskii
Academ-4_SK_Saccoglossus_kowalevskii
gi_780091844_ref_XP_011674128.1_Strongylocentrotus_purpuratus
gi_957859794_ref_XP_014678737.1_Priapulus_caudatus
gi_780009301_ref_XP_011663548.1_Strongylocentrotus_purpuratus
Academ-1_CSa_Ciona_savignyi
Academ-2_Lgigantea_Lottia_gigantea
Academ-1_Lgigantea_Lottia_gigantea
gi_676461609_ref_XP_009055998.1_Lottia_gigantea
Academ-3_CS_Ciona_savignyi
gi_1022758602_gb_KZS05075.1_Daphnia_magna
gi_1022764891_gb_KZS10228.1_Daphnia_magna
Academ-1_Dpulex_Daphnia_pulex
gi_321470338_gb_EFX81314.1_Daphnia_pulex
gi_999983222_gb_KXJ22363.1_Exaiptasia_pallida
gi_999967976_gb_KXJ07150.1_Exaiptasia_pallida
Academ-4_BF_Branchiostoma_floridae
gi_260817557_ref_XP_002603652.1_Branchiostoma_floridae
Academ-2_PM_Petromyzon_marinus
gi_260819138_ref_XP_002604894.1_Branchiostoma_floridae
gi_260782791_ref_XP_002586465.1_Branchiostoma_floridae
Academ-1_BF_Branchiostoma_floridae
gi_260808991_ref_XP_002599290.1_Branchiostoma_floridae
gi_1192754932_ref_XP_021000773.1_Parasteatoda_tepidariorum
gi_1603812507_ref_XP_028410461.1_Dendronephthya_gigantea
gi_1603818650_ref_XP_028413737.1_Dendronephthya_gigantea
gi_1603815071_ref_XP_028411862.1_Dendronephthya_gigantea
gi_1603784202_ref_XP_028395413.1_Dendronephthya_gigantea
gi_1275004128_gb_PIK50721.1_Apostichopus_japonicus
gi_779996353_ref_XP_011675106.1_Strongylocentrotus_purpuratus
gi_780055989_ref_XP_011669768.1_Strongylocentrotus_purpuratus
gi_780003299_ref_XP_011683244.1_Strongylocentrotus_purpuratus
gi_1275012319_gb_PIK58612.1_Apostichopus_japonicus
Academ-1_SP_Strongylocentrotus_purpuratus
gi_780182552_ref_XP_011662219.1_Strongylocentrotus_purpuratus
gi_1274991596_gb_PIK39062.1_Apostichopus_japonicus
gi_780176930_ref_XP_011661467.1_Strongylocentrotus_purpuratus
gi_780090510_ref_XP_011673963.1_Strongylocentrotus_purpuratus
gi_1275015268_gb_PIK61485.1_Apostichopus_japonicus
gi_1274997413_gb_PIK44379.1_Apostichopus_japonicus
gi_1274992743_gb_PIK40093.1_Apostichopus_japonicus
Academ-2_SP_Strongylocentrotus_purpuratus
gi_780102019_ref_XP_011675373.1_Strongylocentrotus_purpuratus
gi_780092249_ref_XP_001199031.2_Strongylocentrotus_purpuratus
gi_780108444_ref_XP_011676157.1_Strongylocentrotus_purpuratus
Academ-2_CS_Ciona_savignyi
gi_974098318_ref_XP_015247746.1_Cyprinodon_variegatus
Academ-1_GA_Gasterosteus_aculeatus
Academ-1_PM_Petromyzon_marinus
gi_1020559760_ref_XP_016111421.1_Sinocyclocheilus_grahami
gi_1249017427_ref_XP_022535652.1_Astyanax_mexicanus
Academ-1_HM_Hydra_vulgaris
gi_1603821128_ref_XP_028415042.1_Dendronephthya_gigantea
gi_1603773010_ref_XP_028408550.1_Dendronephthya_gigantea
gi_1603814823_ref_XP_028411727.1_Dendronephthya_gigantea
gi_1603802500_ref_XP_028405219.1_Dendronephthya_gigantea
gi_1603777808_ref_XP_028391978.1_Dendronephthya_gigantea
gi_1603777806_ref_XP_028391977.1_Dendronephthya_gigantea
gi_1603800430_ref_XP_028404105.1_Dendronephthya_gigantea
gi_1603777810_ref_XP_028391979.1_Dendronephthya_gigantea
gi_1603785586_ref_XP_028396142.1_Dendronephthya_gigantea
gi_1603785588_ref_XP_028396143.1_Dendronephthya_gigantea
gi_1603785584_ref_XP_028396141.1_Dendronephthya_gigantea
gi_1603808662_ref_XP_028408500.1_Dendronephthya_gigantea
gi_1603812558_ref_XP_028410493.1_Dendronephthya_gigantea
gi_1603819628_ref_XP_028414245.1_Dendronephthya_gigantea
gi_1603775153_ref_XP_028419097.1_Dendronephthya_gigantea
gi_1603791072_ref_XP_028399085.1_Dendronephthya_gigantea
gi_1603819299_ref_XP_028414070.1_Dendronephthya_gigantea
0.9

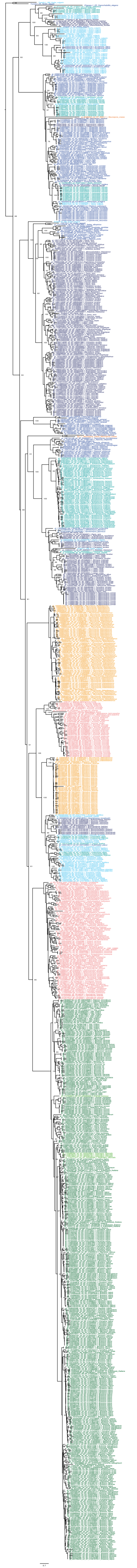

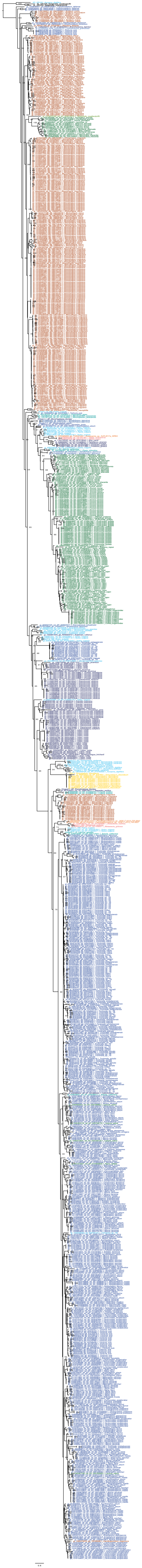

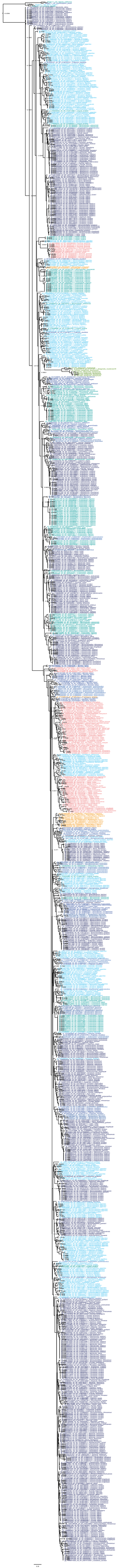

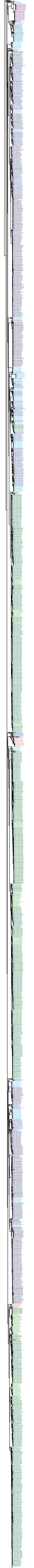

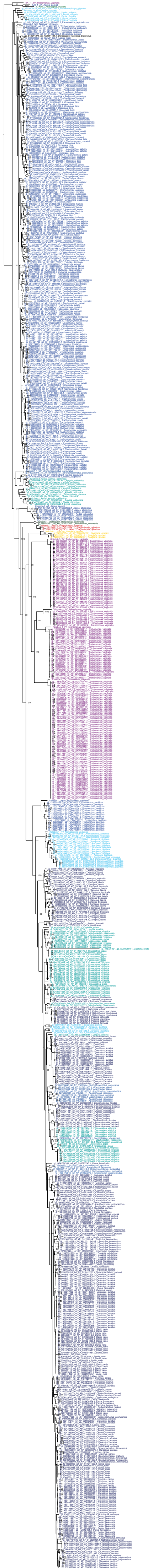

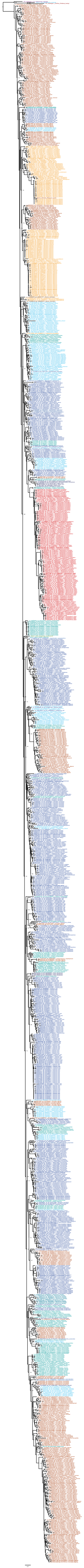

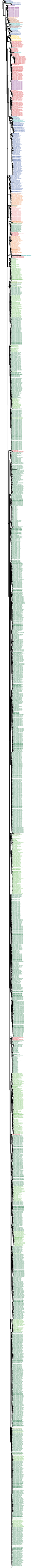

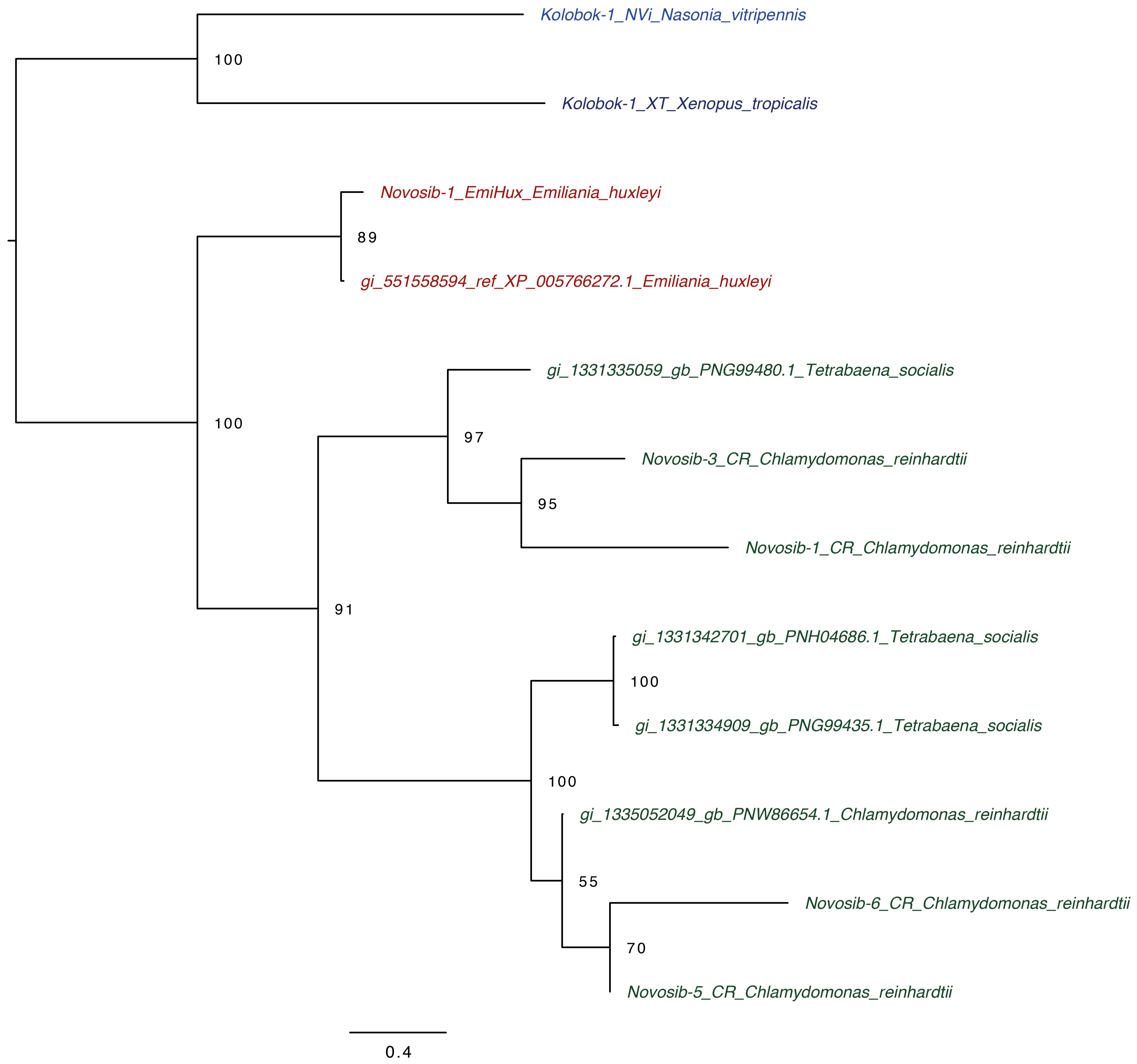
Kolobok-1_NVi_Nasonia_vitripennis
100
Kolobok-1_XT_Xenopus_tropicalis
Novosib-1_EmiHux_Emiliania_huxleyi
89
gi_551558594_ref_XP_005766272.1_Emiliania_huxleyi
gi_1331335059_gb_PNG99480.1_Tetrabaena_socialis
100
97
Novosib-3_CR_Chlamydomonas_reinhardtii
95
Novosib-1_CR_Chlamydomonas_reinhardtii
91
gi_1331342701_gb_PNH04686.1_Tetrabaena_socialis
100
gi_1331334909_gb_PNG99435.1_Tetrabaena_socialis
100
gi_1335052049_gb_PNW86654.1_Chlamydomonas_reinhardtii
55
Novosib-6_CR_Chlamydomonas_reinhardtii
70
Novosib-5_CR_Chlamydomonas_reinhardtii
0.4

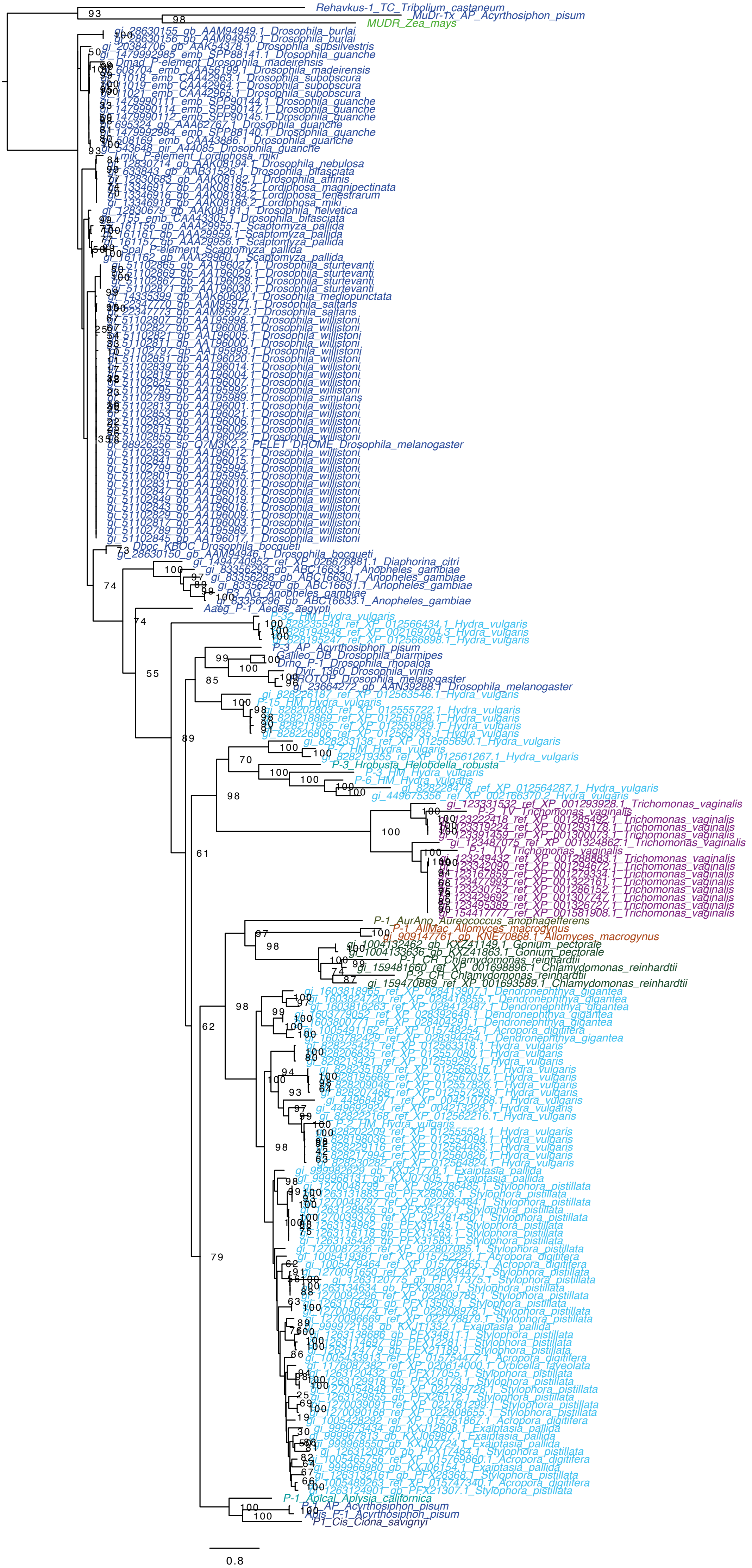

Rehavkus-1_TC_Tribolium_castaneum
MuDr-Tx_AP_Acyrthosiphon_pisum
MUDR_Zea_mays
Dmad_P-element_Drosophila_madeirensis
Lmik_P-element_Lordiphosa_miki
Spal_P-element_Scaptomyza_pallida
Dboc_KBOC_Drosophila_bocqueti
P3_AG_Anopheles_gambiae
Aaeg_P-1_Aedes_aegypti
P-32_HM_Hydra_vulgaris
P-3_AP_Acyrthosiphon_pisum
Galileo_DB_Drosophila_biarmipes
Drho_P-1_Drosophila_rhopaloa
Dvir_1360_Drosophila_virilis
PROTOP_Drosophila_melanogaster
P-15_HM_Hydra_vulgaris
P-7_HM_Hydra_vulgaris
P-3_Hrobusta_Helobdella_robusta
P-3_HM_Hydra_vulgaris
P-6_HM_Hydra_vulgaris
P-2_TV_Trichomonas_vaginalis
P-1_TV_Trichomonas_vaginalis
P-1_AurAno_Aureococcus_anophagefferens
P-1_AllMac_Allomyces_macrogynus
P-1_CR_Chlamydomonas_reinhardtii
P-2_CR_Chlamydomonas_reinhardtii
P-2_HM_Hydra_vulgaris
P-1_Aplcal_Aplysia_californica
P-1_AP_Acyrthosiphon_pisum
Apis_P-1_Acyrthosiphon_pisum
P1_Cis_Ciona_savignyi
0.8

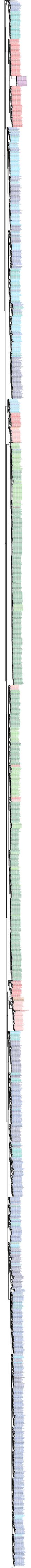

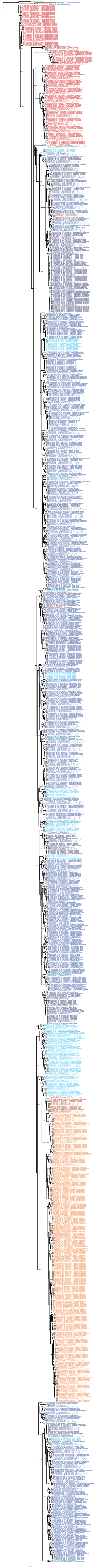

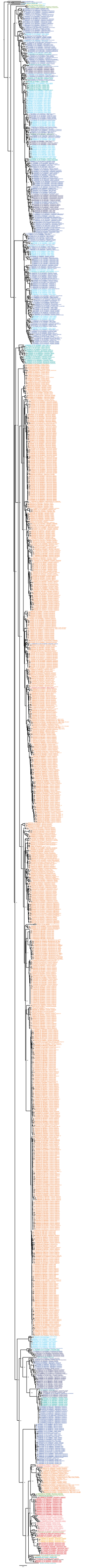

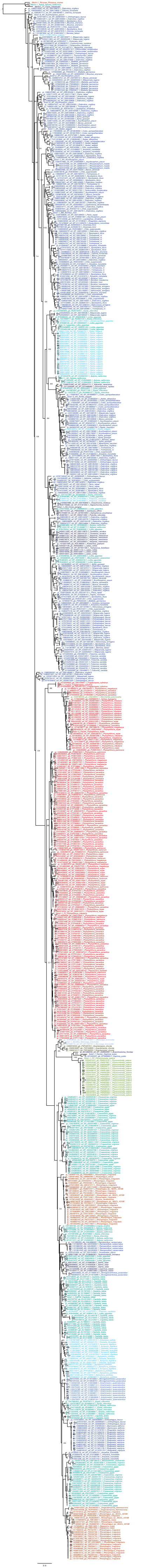

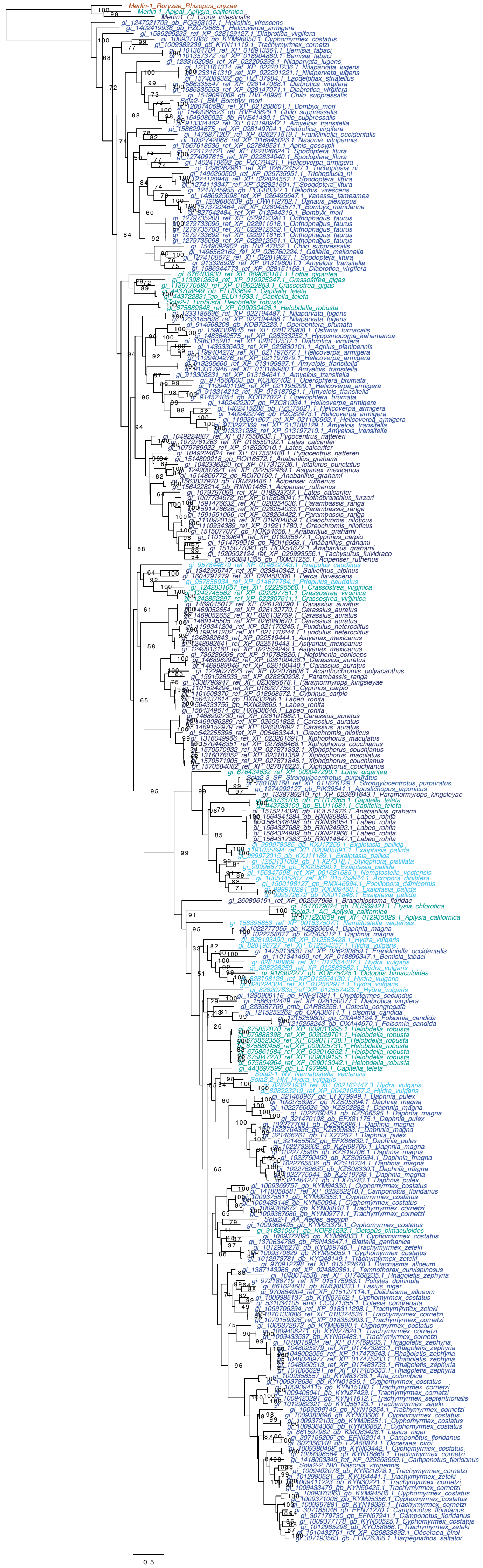
Merlin-1_Roryzae_Rhizopus_oryzae
Merlin-1_Apical_Aplysia_californica
Merlin1_CI_Ciona_intestinalis
0.5

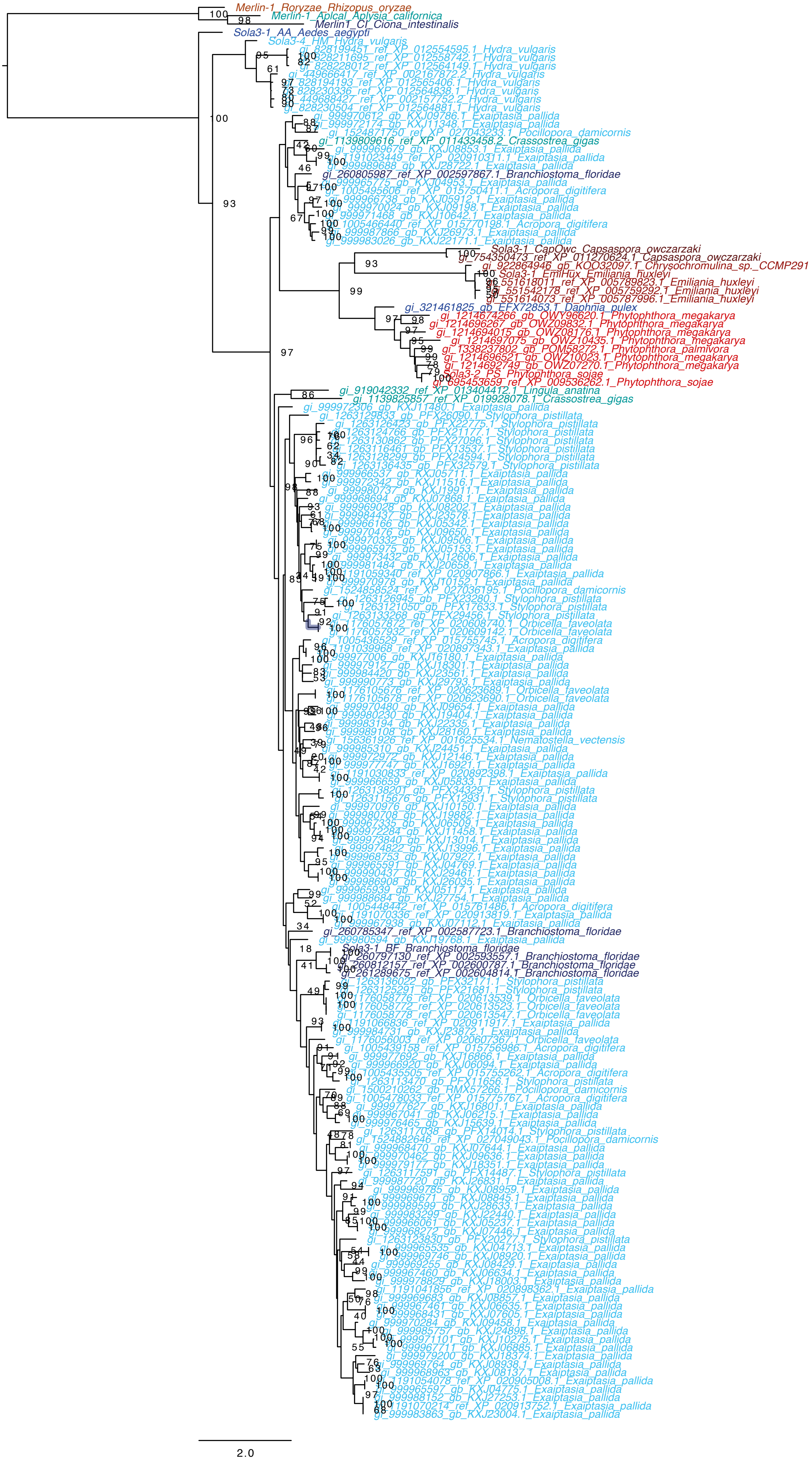
Merlin-1_Roryzae_Rhizopus_oryzae
Merlin-1_Aplcal_Aplysia_californica
Merlin1_CI_Ciona_intestinalis
Sola3-1_AA_Aedes_aegypti
Sola3-4_HM_Hydra_vulgaris
gi_828199451_ref_XP_012554595.1_Hydra_vulgaris
gi_828211695_ref_XP_012558742.1_Hydra_vulgaris
gi_828228012_ref_XP_012564149.1_Hydra_vulgaris
gi_449666417_ref_XP_002167872.2_Hydra_vulgaris
gi_828194193_ref_XP_012565406.1_Hydra_vulgaris
gi_828230336_ref_XP_012564838.1_Hydra_vulgaris
gi_449688427_ref_XP_002157752.2_Hydra_vulgaris
gi_828230504_ref_XP_012564881.1_Hydra_vulgaris
gi_999970612_gb_KXJ09786.1_Exaiptasia_pallida
gi_999972174_gb_KXJ11348.1_Exaiptasia_pallida
gi_1524871750_ref_XP_027043233.1_Pocillopora_damicornis
gi_1139809616_ref_XP_011433458.2_Crassostrea_gigas
gi_999969679_gb_KXJ08853.1_Exaiptasia_pallida
gi_1191023449_ref_XP_020910311.1_Exaiptasia_pallida
gi_999989688_gb_KXJ28722.1_Exaiptasia_pallida
gi_260805987_ref_XP_002597867.1_Branchiostoma_floridae
gi_999965775_gb_KXJ04953.1_Exaiptasia_pallida
gi_1005495606_ref_XP_015750411.1_Acropora_digitifera
gi_999966738_gb_KXJ05912.1_Exaiptasia_pallida
gi_999970024_gb_KXJ09198.1_Exaiptasia_pallida
gi_999971468_gb_KXJ10642.1_Exaiptasia_pallida
gi_1005466440_ref_XP_015770198.1_Acropora_digitifera
gi_999987866_gb_KXJ26973.1_Exaiptasia_pallida
gi_999983026_gb_KXJ22171.1_Exaiptasia_pallida
Sola3-1_CapOwc_Capsaspora_owczarzaki
gi_754350473_ref_XP_011270624.1_Capsaspora_owczarzaki
gi_922864946_gb_KOO32097.1_Chrysochromulina_sp._CCMP291
Sola3-1_EmiHux_Emiliania_huxleyi
gi_551618011_ref_XP_005789823.1_Emiliania_huxleyi
gi_551542178_ref_XP_005759292.1_Emiliania_huxleyi
gi_551614073_ref_XP_005787996.1_Emiliania_huxleyi
gi_321461825_gb_EFX72853.1_Daphnia_pulex
gi_1214674266_gb_OWY96620.1_Phytophthora_megakarya
gi_1214696267_gb_OWZ09832.1_Phytophthora_megakarya
gi_1214694015_gb_OWZ08176.1_Phytophthora_megakarya
gi_1214697075_gb_OWZ10435.1_Phytophthora_megakarya
gi_1338237802_gb_POM58272.1_Phytophthora_palmivora
gi_1214696521_gb_OWZ10023.1_Phytophthora_megakarya
gi_1214692749_gb_OWZ07270.1_Phytophthora_megakarya
Sola3-2_PS_Phytophthora_sojae
gi_695453659_ref_XP_009536262.1_Phytophthora_sojae
gi_919042332_ref_XP_013404412.1_Lingula_anatina
gi_1139825857_ref_XP_019928078.1_Crassostrea_gigas
gi_999972306_gb_KXJ11480.1_Exaiptasia_pallida
gi_1263129833_gb_PFX26090.1_Stylophora_pistillata
gi_1263126423_gb_PFX22775.1_Stylophora_pistillata
gi_1263124766_gb_PFX21177.1_Stylophora_pistillata
gi_1263130862_gb_PFX27096.1_Stylophora_pistillata
gi_1263116461_gb_PFX13537.1_Stylophora_pistillata
gi_1263128299_gb_PFX24594.1_Stylophora_pistillata
gi_1263136435_gb_PFX32579.1_Stylophora_pistillata
gi_999966537_gb_KXJ05711.1_Exaiptasia_pallida
gi_999972342_gb_KXJ11516.1_Exaiptasia_pallida
gi_999980737_gb_KXJ19911.1_Exaiptasia_pallida
gi_999968694_gb_KXJ07868.1_Exaiptasia_pallida
gi_999969028_gb_KXJ08202.1_Exaiptasia_pallida
gi_999984437_gb_KXJ23578.1_Exaiptasia_pallida
gi_999966166_gb_KXJ05342.1_Exaiptasia_pallida
gi_999970476_gb_KXJ09650.1_Exaiptasia_pallida
gi_999970332_gb_KXJ09506.1_Exaiptasia_pallida
gi_999965975_gb_KXJ05153.1_Exaiptasia_pallida
gi_999973432_gb_KXJ12606.1_Exaiptasia_pallida
gi_999981484_gb_KXJ20658.1_Exaiptasia_pallida
gi_1191059340_ref_XP_020907866.1_Exaiptasia_pallida
gi_999970978_gb_KXJ10152.1_Exaiptasia_pallida
gi_1524858524_ref_XP_027036195.1_Pocillopora_damicornis
gi_1263126945_gb_PFX23280.1_Stylophora_pistillata
gi_1263121050_gb_PFX17633.1_Stylophora_pistillata
gi_1263133268_gb_PFX29456.1_Stylophora_pistillata
gi_1176057872_ref_XP_020608740.1_Orbicella_faveolata
gi_1176057932_ref_XP_020609142.1_Orbicella_faveolata
gi_1005436529_ref_XP_015755745.1_Acropora_digitifera
gi_1191039968_ref_XP_020897343.1_Exaiptasia_pallida
gi_999977006_gb_KXJ16180.1_Exaiptasia_pallida
gi_999979127_gb_KXJ18301.1_Exaiptasia_pallida
gi_999984420_gb_KXJ23561.1_Exaiptasia_pallida
gi_999990773_gb_KXJ29793.1_Exaiptasia_pallida
gi_1176105676_ref_XP_020623689.1_Orbicella_faveolata
gi_1176105678_ref_XP_020623690.1_Orbicella_faveolata
gi_999970480_gb_KXJ09654.1_Exaiptasia_pallida
gi_999980230_gb_KXJ19404.1_Exaiptasia_pallida
gi_999983194_gb_KXJ22335.1_Exaiptasia_pallida
gi_999989108_gb_KXJ28160.1_Exaiptasia_pallida
gi_156361926_ref_XP_001625534.1_Nematostella_vectensis
gi_999985310_gb_KXJ24451.1_Exaiptasia_pallida
gi_999972972_gb_KXJ12146.1_Exaiptasia_pallida
gi_999977747_gb_KXJ16921.1_Exaiptasia_pallida
gi_1191030833_ref_XP_020892398.1_Exaiptasia_pallida
gi_999966659_gb_KXJ05833.1_Exaiptasia_pallida
gi_1263138207_gb_PFX34329.1_Stylophora_pistillata
gi_1263115676_gb_PFX12931.1_Stylophora_pistillata
gi_999970976_gb_KXJ10150.1_Exaiptasia_pallida
gi_999980708_gb_KXJ19882.1_Exaiptasia_pallida
gi_999967335_gb_KXJ06509.1_Exaiptasia_pallida
gi_999972284_gb_KXJ11458.1_Exaiptasia_pallida
gi_999973840_gb_KXJ13014.1_Exaiptasia_pallida
gi_999974822_gb_KXJ13996.1_Exaiptasia_pallida
gi_999968753_gb_KXJ07927.1_Exaiptasia_pallida
gi_999965591_gb_KXJ04769.1_Exaiptasia_pallida
gi_999990437_gb_KXJ29461.1_Exaiptasia_pallida
gi_999986908_gb_KXJ26035.1_Exaiptasia_pallida
gi_999965939_gb_KXJ05117.1_Exaiptasia_pallida
gi_999988684_gb_KXJ27754.1_Exaiptasia_pallida
gi_1005448442_ref_XP_015761486.1_Acropora_digitifera
gi_1191070336_ref_XP_020913819.1_Exaiptasia_pallida
gi_999967938_gb_KXJ07112.1_Exaiptasia_pallida
gi_260785347_ref_XP_002587723.1_Branchiostoma_floridae
gi_999980594_gb_KXJ19768.1_Exaiptasia_pallida
Sola3-1_BF_Branchiostoma_floridae
gi_260797130_ref_XP_002593557.1_Branchiostoma_floridae
gi_260812157_ref_XP_002600787.1_Branchiostoma_floridae
gi_261289675_ref_XP_002604814.1_Branchiostoma_floridae
gi_1263136022_gb_PFX32171.1_Stylophora_pistillata
gi_1263125291_gb_PFX21681.1_Stylophora_pistillata
gi_1176058776_ref_XP_020613539.1_Orbicella_faveolata
gi_1176058772_ref_XP_020613523.1_Orbicella_faveolata
gi_1176058778_ref_XP_020613547.1_Orbicella_faveolata
gi_1191066836_ref_XP_020911917.1_Exaiptasia_pallida
gi_999984731_gb_KXJ23872.1_Exaiptasia_pallida
gi_1176056003_ref_XP_020607367.1_Orbicella_faveolata
gi_1005439158_ref_XP_015756986.1_Acropora_digitifera
gi_999977692_gb_KXJ16866.1_Exaiptasia_pallida
gi_999966920_gb_KXJ06094.1_Exaiptasia_pallida
gi_1005435505_ref_XP_015755262.1_Acropora_digitifera
gi_1263113470_gb_PFX11656.1_Stylophora_pistillata
gi_1500210262_gb_RMX57266.1_Pocillopora_damicornis
gi_1005478033_ref_XP_015775767.1_Acropora_digitifera
gi_999977627_gb_KXJ16801.1_Exaiptasia_pallida
gi_999967041_gb_KXJ06215.1_Exaiptasia_pallida
gi_999976465_gb_KXJ15639.1_Exaiptasia_pallida
gi_1263117038_gb_PFX14014.1_Stylophora_pistillata
gi_1524882646_ref_XP_027049043.1_Pocillopora_damicornis
gi_999968470_gb_KXJ07644.1_Exaiptasia_pallida
gi_999970462_gb_KXJ09636.1_Exaiptasia_pallida
gi_999979177_gb_KXJ18351.1_Exaiptasia_pallida
gi_1263117591_gb_PFX14487.1_Stylophora_pistillata
gi_999987720_gb_KXJ26831.1_Exaiptasia_pallida
gi_999969785_gb_KXJ08959.1_Exaiptasia_pallida
gi_999969671_gb_KXJ08845.1_Exaiptasia_pallida
gi_999989599_gb_KXJ28633.1_Exaiptasia_pallida
gi_999983299_gb_KXJ22440.1_Exaiptasia_pallida
gi_999966061_gb_KXJ05237.1_Exaiptasia_pallida
gi_999968272_gb_KXJ07446.1_Exaiptasia_pallida
gi_1263123830_gb_PFX20277.1_Stylophora_pistillata
gi_999965535_gb_KXJ04713.1_Exaiptasia_pallida
gi_999969746_gb_KXJ08920.1_Exaiptasia_pallida
gi_999969255_gb_KXJ08429.1_Exaiptasia_pallida
gi_999967460_gb_KXJ06634.1_Exaiptasia_pallida
gi_999978829_gb_KXJ18003.1_Exaiptasia_pallida
gi_1191041856_ref_XP_020898362.1_Exaiptasia_pallida
gi_999969683_gb_KXJ08857.1_Exaiptasia_pallida
gi_999967461_gb_KXJ06635.1_Exaiptasia_pallida
gi_999968431_gb_KXJ07605.1_Exaiptasia_pallida
gi_999970284_gb_KXJ09458.1_Exaiptasia_pallida
gi_999985757_gb_KXJ24898.1_Exaiptasia_pallida
gi_999971101_gb_KXJ10275.1_Exaiptasia_pallida
gi_999967711_gb_KXJ06885.1_Exaiptasia_pallida
gi_999979200_gb_KXJ18374.1_Exaiptasia_pallida
gi_999969764_gb_KXJ08938.1_Exaiptasia_pallida
gi_999968963_gb_KXJ08137.1_Exaiptasia_pallida
gi_1191054078_ref_XP_020905008.1_Exaiptasia_pallida
gi_999965597_gb_KXJ04775.1_Exaiptasia_pallida
gi_999988152_gb_KXJ27253.1_Exaiptasia_pallida
gi_1191070214_ref_XP_020913752.1_Exaiptasia_pallida
gi_999983863_gb_KXJ23004.1_Exaiptasia_pallida
2.0

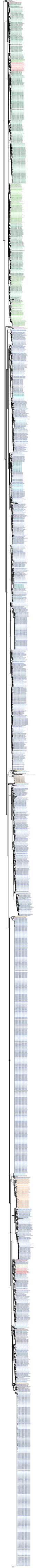

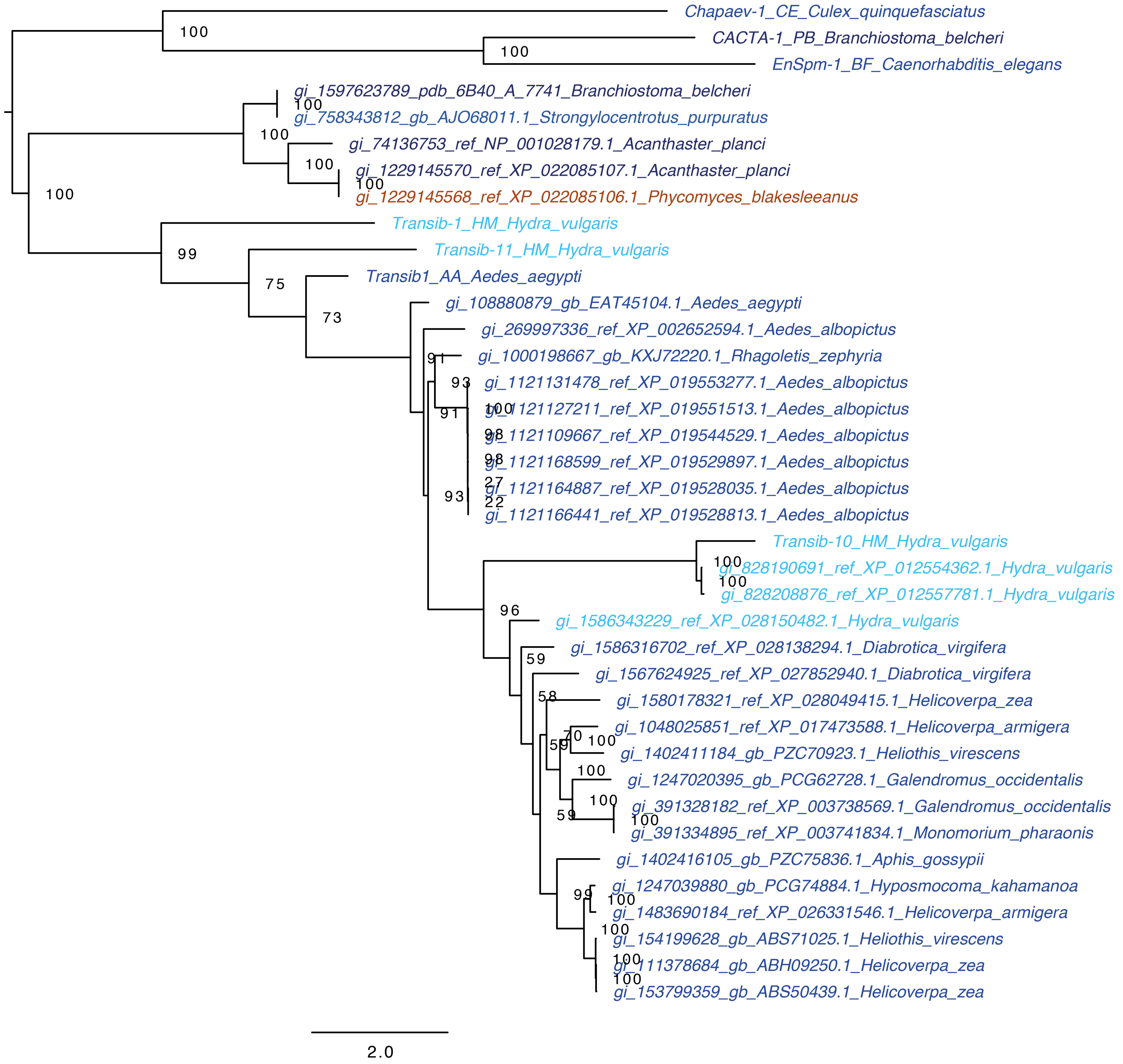
Chapaev-1_CE_Culex_quinquefasciatus
CACTA-1_PB_Branchiostoma_belcheri
EnSpm-1_BF_Caenorhabditis_elegans
gi_1597623789_pdb_6B40_A_7741_Branchiostoma_belcheri
gi_758343812_gb_AJO68011.1_Strongylocentrotus_purpuratus
gi_74136753_ref_NP_001028179.1_Acanthaster_planci
gi_1229145570_ref_XP_022085107.1_Acanthaster_planci
gi_1229145568_ref_XP_022085106.1_Phycomyces_blakesleeanus
Transib-1_HM_Hydra_vulgaris
Transib-11_HM_Hydra_vulgaris
Transib1_AA_Aedes_aegypti
gi_108880879_gb_EAT45104.1_Aedes_aegypti
gi_269997336_ref_XP_002652594.1_Aedes_albopictus
gi_1000198667_gb_KXJ72220.1_Rhagoletis_zephyria
gi_1121131478_ref_XP_019553277.1_Aedes_albopictus
gi_1121127211_ref_XP_019551513.1_Aedes_albopictus
gi_1121109667_ref_XP_019544529.1_Aedes_albopictus
gi_1121168599_ref_XP_019529897.1_Aedes_albopictus
gi_1121164887_ref_XP_019528035.1_Aedes_albopictus
gi_1121166441_ref_XP_019528813.1_Aedes_albopictus
Transib-10_HM_Hydra_vulgaris
gi_828190691_ref_XP_012554362.1_Hydra_vulgaris
gi_828208876_ref_XP_012557781.1_Hydra_vulgaris
gi_1586343229_ref_XP_028150482.1_Hydra_vulgaris
gi_1586316702_ref_XP_028138294.1_Diabrotica_virgifera
gi_1567624925_ref_XP_027852940.1_Diabrotica_virgifera
gi_1580178321_ref_XP_028049415.1_Helicoverpa_zea
gi_1048025851_ref_XP_017473588.1_Helicoverpa_armigera
gi_1402411184_gb_PZC70923.1_Heliothis_virescens
gi_1247020395_gb_PCG62728.1_Galendromus_occidentalis
gi_391328182_ref_XP_003738569.1_Galendromus_occidentalis
gi_391334895_ref_XP_003741834.1_Monomorium_pharaonis
gi_1402416105_gb_PZC75836.1_Aphis_gossypii
gi_1247039880_gb_PCG74884.1_Hyposmocoma_kahamanoa
gi_1483690184_ref_XP_026331546.1_Helicoverpa_armigera
gi_154199628_gb_ABS71025.1_Heliothis_virescens
gi_111378684_gb_ABH09250.1_Helicoverpa_zea
gi_153799359_gb_ABS50439.1_Helicoverpa_zea
100
100
100
100
100
100
100
99
75
73
91
93
91
100
98
98
27
93
22
100
100
96
59
58
70
59
100
100
100
59
100
99
100
100
100
100
2.0

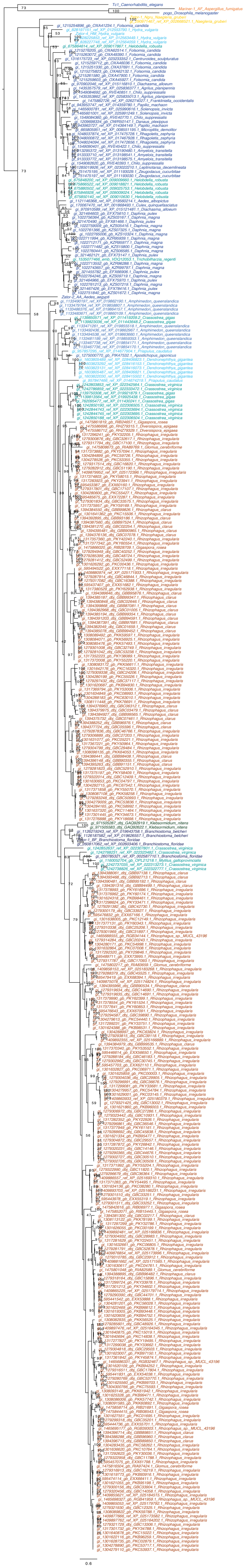

Academem

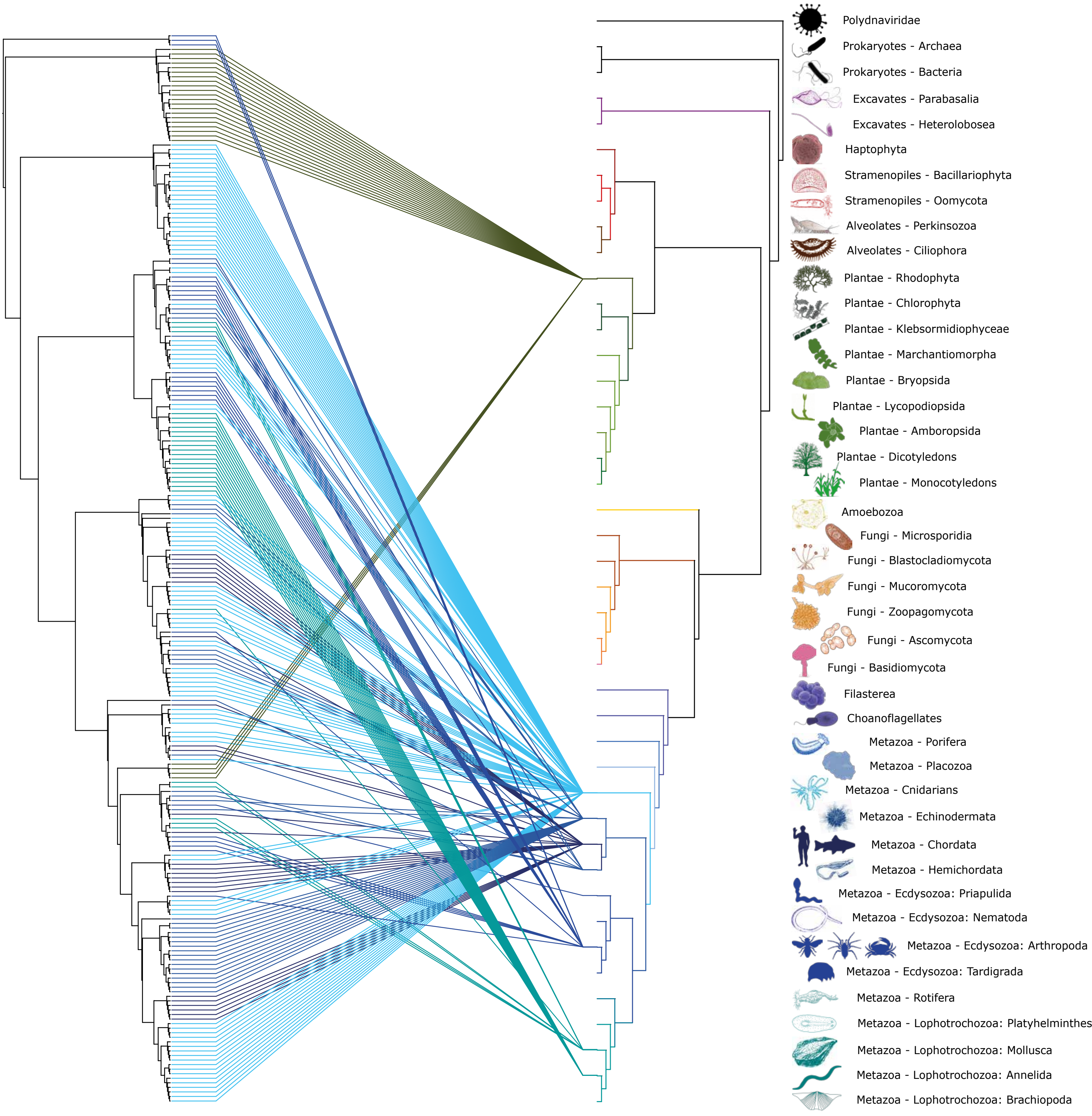

Polydnaviridae
Prokaryotes - Archaea
Prokaryotes - Bacteria
Excavates - Parabasalia
Excavates - Heterolobosea
Haptophyta
Stramenopiles - Bacillariophyta
Stramenopiles - Oomycota
Alveolates - Perkinsozoa
Alveolates - Ciliophora
Plantae - Rhodophyta
Plantae - Chlorophyta
Plantae - Klebsormidiophyceae
Plantae - Marchantiomorpha
Plantae - Bryopsida
Plantae - Lycopodiopsida
Plantae - Amboropsida
Plantae - Dicotyledons
Plantae - Monocotyledons
Amoebozoa
Fungi - Microsporidia
Fungi - Blastocladiomycota
Fungi - Mucoromycota
Fungi - Zoopagomycota
Fungi - Ascomycota
Fungi - Basidiomycota
Filasterea
Choanoflagellates
Metazoa - Porifera
Metazoa - Placozoa
Metazoa - Cnidarians
Metazoa - Echinodermata
Metazoa - Chordata
Metazoa - Hemichordata
Metazoa - Ecdysozoa: Priapulida
Metazoa - Ecdysozoa: Nematoda
Metazoa - Ecdysozoa: Arthropoda
Metazoa - Ecdysozoa: Tardigrada
Metazoa - Rotifera
Metazoa - Lophotrochozoa: Platyhelminthes
Metazoa - Lophotrochozoa: Mollusca
Metazoa - Lophotrochozoa: Annelida
Metazoa - Lophotrochozoa: Brachiopoda

#### Academ

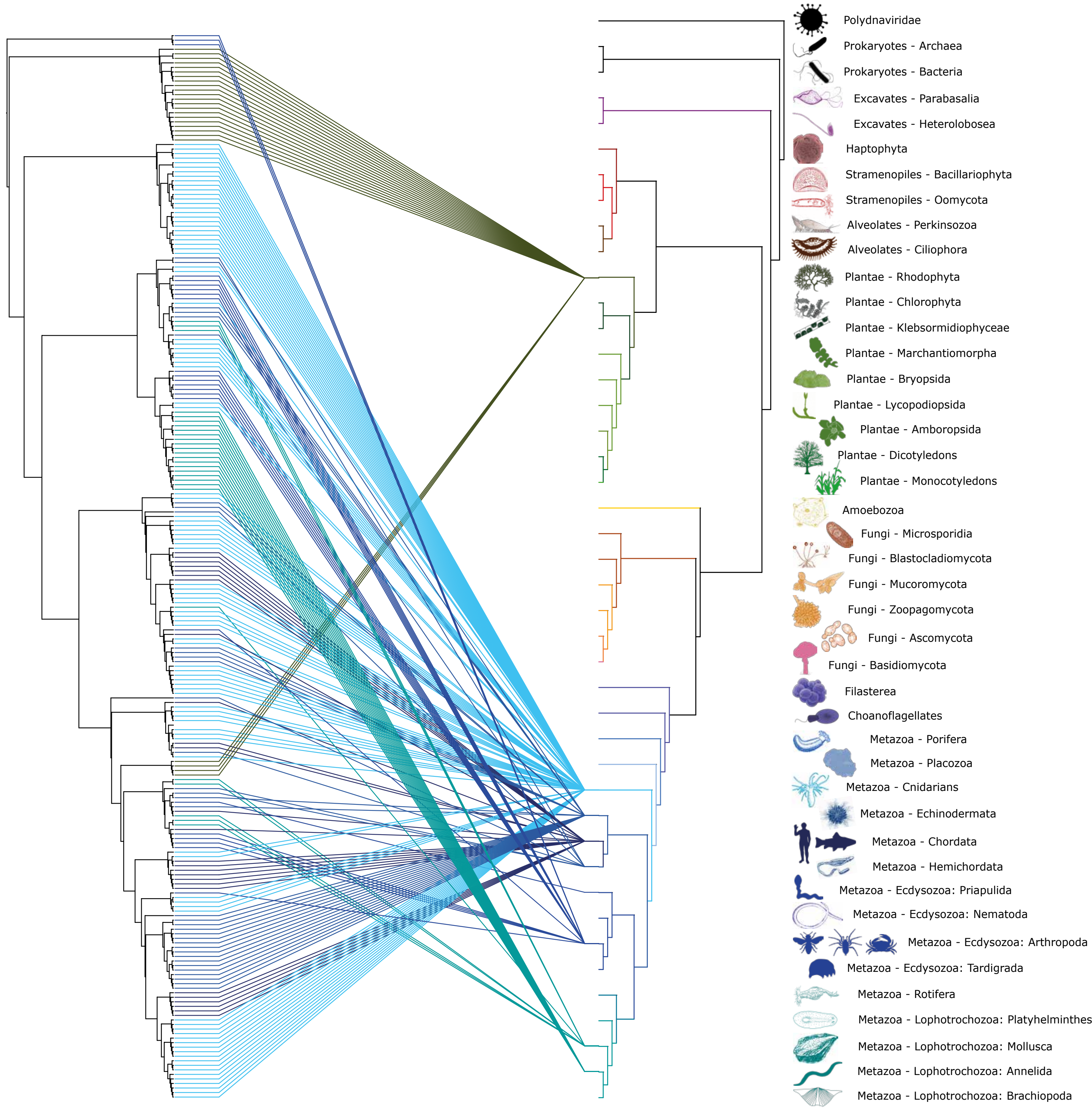

Polydnaviridae
Prokaryotes - Archaea
Prokaryotes - Bacteria
Excavates - Parabasalia
Excavates - Heterolobosea
Haptophyta
Stramenopiles - Bacillariophyta
Stramenopiles - Oomycota
Alveolates - Perkinsozoa
Alveolates - Ciliophora
Plantae - Rhodophyta
Plantae - Chlorophyta
Plantae - Klebsormidiophyceae
Plantae - Marchantiomorpha
Plantae - Bryopsida
Plantae - Lycopodiopsida
Plantae - Amboropsida
Plantae - Dicotyledons
Plantae - Monocotyledons
Amoebozoa
Fungi - Microsporidia
Fungi - Blastocladiomycota
Fungi - Mucoromycota
Fungi - Zoopagomycota
Fungi - Ascomycota
Fungi - Basidiomycota
Filasterea
Choanoflagellates
Metazoa - Porifera
Metazoa - Placozoa
Metazoa - Cnidarians
Metazoa - Echinodermata
Metazoa - Chordata
Metazoa - Hemichordata
Metazoa - Ecdysozoa: Priapulida
Metazoa - Ecdysozoa: Nematoda
Metazoa - Ecdysozoa: Arthropoda
Metazoa - Ecdysozoa: Tardigrada
Metazoa - Rotifera
Metazoa - Lophotrochozoa: Platyhelminthes
Metazoa - Lophotrochozoa: Mollusca
Metazoa - Lophotrochozoa: Annelida
Metazoa - Lophotrochozoa: Brachiopoda

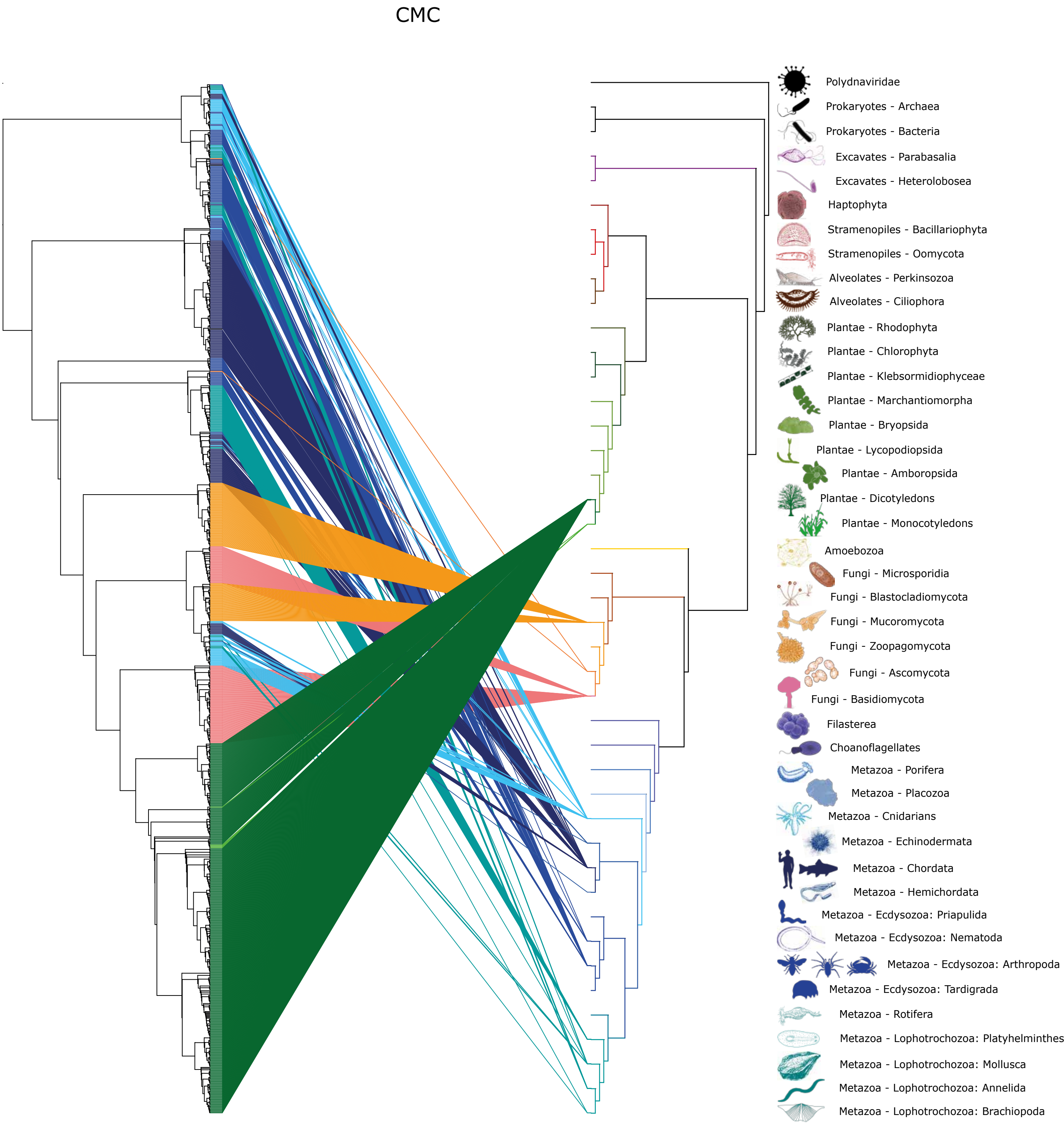
CMC
Polydnaviridae
Prokaryotes - Archaea
Prokaryotes - Bacteria
Excavates - Parabasalia
Excavates - Heterolobosea
Haptophyta
Stramenopiles - Bacillariophyta
Stramenopiles - Oomycota
Alveolates - Perkinsozoa
Alveolates - Ciliophora
Plantae - Rhodophyta
Plantae - Chlorophyta
Plantae - Klebsormidiophyceae
Plantae - Marchantiomorpha
Plantae - Bryopsida
Plantae - Lycopodiopsida
Plantae - Amboropsida
Plantae - Dicotyledons
Plantae - Monocotyledons
Amoebozoa
Fungi - Microsporidia
Fungi - Blastocladiomycota
Fungi - Mucoromycota
Fungi - Zoopagomycota
Fungi - Ascomycota
Fungi - Basidiomycota
Filasterea
Choanoflagellates
Metazoa - Porifera
Metazoa - Placozoa
Metazoa - Cnidarians
Metazoa - Echinodermata
Metazoa - Chordata
Metazoa - Hemichordata
Metazoa - Ecdysozoa: Priapulida
Metazoa - Ecdysozoa: Nematoda
Metazoa - Ecdysozoa: Arthropoda
Metazoa - Ecdysozoa: Tardigrada
Metazoa - Rotifera
Metazoa - Lophotrochozoa: Platyhelminthes
Metazoa - Lophotrochozoa: Mollusca
Metazoa - Lophotrochozoa: Annelida
Metazoa - Lophotrochozoa: Brachiopoda

### Ginger

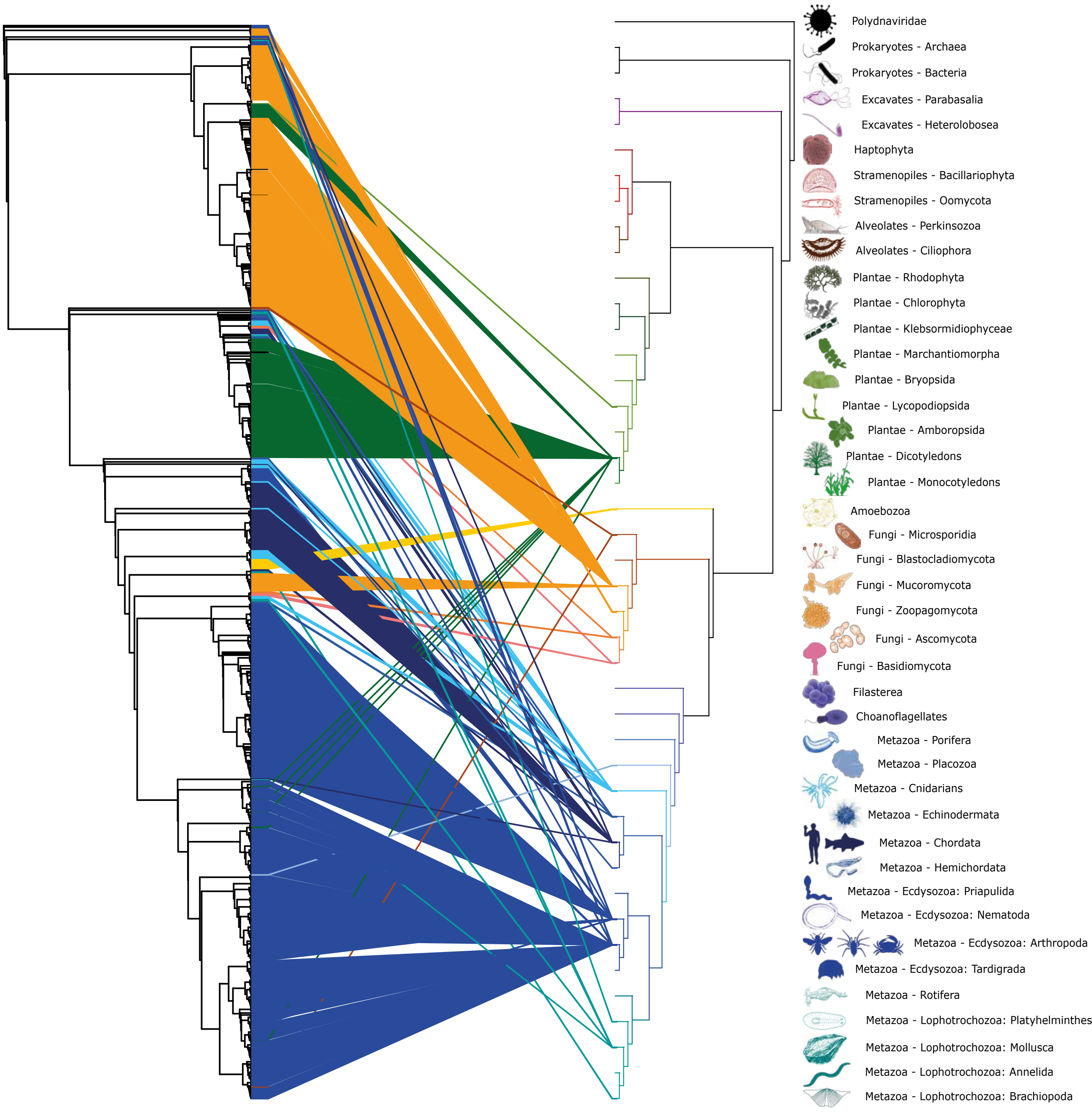

Polydnaviridae
Prokaryotes - Archaea
Prokaryotes - Bacteria
Excavates - Parabasalia
Excavates - Heterolobosea
Haptophyta
Stramenopiles - Bacillariophyta
Stramenopiles - Oomycota
Alveolates - Perkinsozoa
Alveolates - Ciliophora
Plantae - Rhodophyta
Plantae - Chlorophyta
Plantae - Klebsormidiophyceae
Plantae - Marchantiomorpha
Plantae - Bryopsida
Plantae - Lycopodiopsida
Plantae - Amboropsida
Plantae - Dicotyledons
Plantae - Monocotyledons
Amoebozoa
Fungi - Microsporidia
Fungi - Blastocladiomycota
Fungi - Mucoromycota
Fungi - Zoopagomycota
Fungi - Ascomycota
Fungi - Basidiomycota
Filasterea
Choanoflagellates
Metazoa - Porifera
Metazoa - Placozoa
Metazoa - Cnidarians
Metazoa - Echinodermata
Metazoa - Chordata
Metazoa - Hemichordata
Metazoa - Ecdysozoa: Priapulida
Metazoa - Ecdysozoa: Nematoda
Metazoa - Ecdysozoa: Arthropoda
Metazoa - Ecdysozoa: Tardigrada
Metazoa - Rotifera
Metazoa - Lophotrochozoa: Platyhelminthes
Metazoa - Lophotrochozoa: Mollusca
Metazoa - Lophotrochozoa: Annelida
Metazoa - Lophotrochozoa: Brachiopoda

GingerRoot

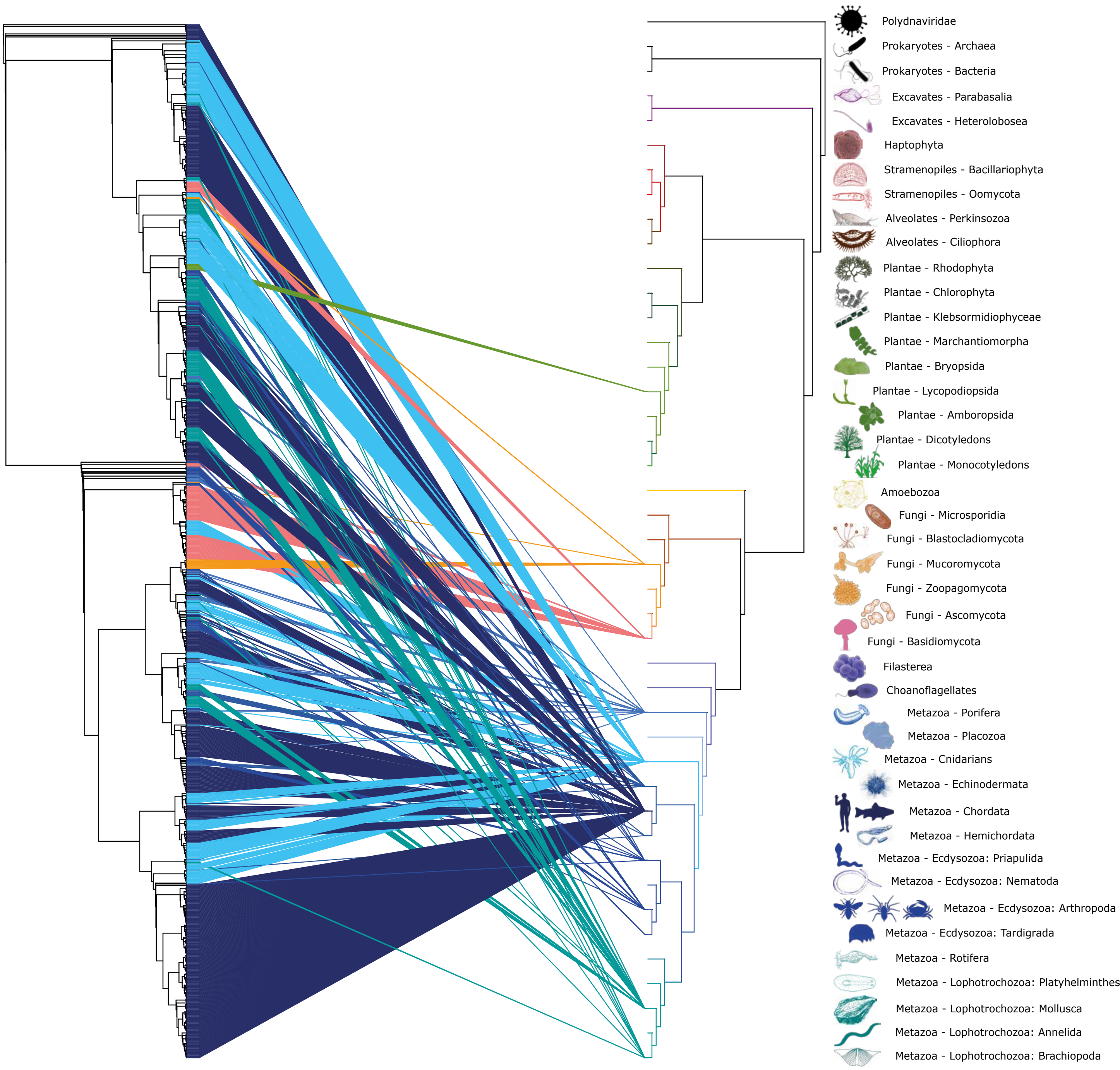

Polydnaviridae
Prokaryotes - Archaea
Prokaryotes - Bacteria
Excavates - Parabasalia
Excavates - Heterolobosea
Haptophyta
Stramenopiles - Bacillariophyta
Stramenopiles - Oomycota
Alveolates - Perkinsozoa
Alveolates - Ciliophora
Plantae - Rhodophyta
Plantae - Chlorophyta
Plantae - Klebsormidiophyceae
Plantae - Marchantiomorpha
Plantae - Bryopsida
Plantae - Lycopodiopsida
Plantae - Amboropsida
Plantae - Dicotyledons
Plantae - Monocotyledons
Amoebozoa
Fungi - Microsporidia
Fungi - Blastocladiomycota
Fungi - Mucoromycota
Fungi - Zoopagomycota
Fungi - Ascomycota
Fungi - Basidiomycota
Filasterea
Choanoflagellates
Metazoa - Porifera
Metazoa - Placozoa
Metazoa - Cnidarians
Metazoa - Echinodermata
Metazoa - Chordata
Metazoa - Hemichordata
Metazoa - Ecdysozoa: Priapulida
Metazoa - Ecdysozoa: Nematoda
Metazoa - Ecdysozoa: Arthropoda
Metazoa - Ecdysozoa: Tardigrada
Metazoa - Rotifera
Metazoa - Lophotrochozoa: Platyhelminthes
Metazoa - Lophotrochozoa: Mollusca
Metazoa - Lophotrochozoa: Annelida
Metazoa - Lophotrochozoa: Brachiopoda

hATs

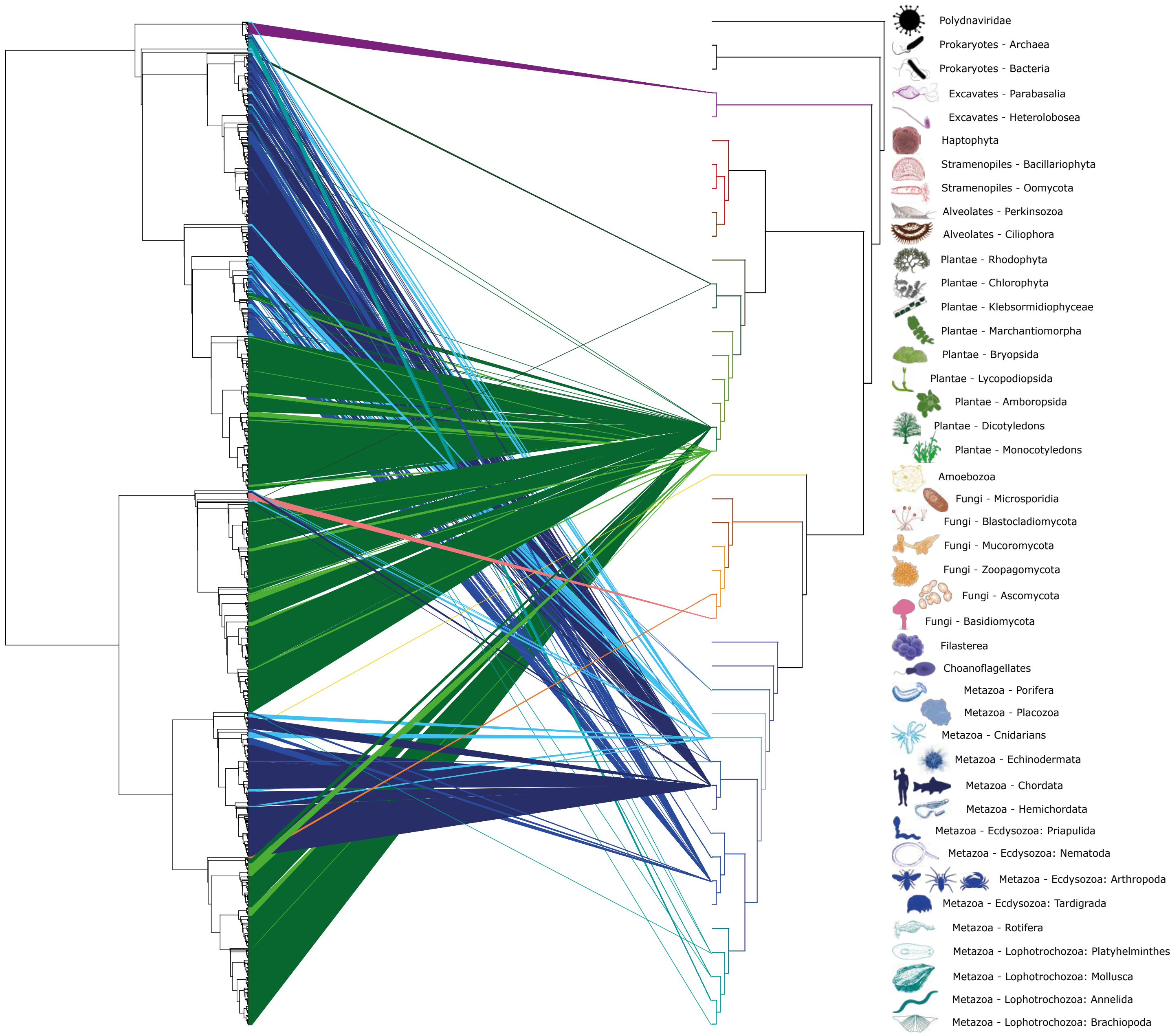
Polydnaviridae
Prokaryotes - Archaea
Prokaryotes - Bacteria
Excavates - Parabasalia
Excavates - Heterolobosea
Haptophyta
Stramenopiles - Bacillariophyta
Stramenopiles - Oomycota
Alveolates - Perkinsozoa
Alveolates - Ciliophora
Plantae - Rhodophyta
Plantae - Chlorophyta
Plantae - Klebsormidiophyceae
Plantae - Marchantiomorpha
Plantae - Bryopsida
Plantae - Lycopodiopsida
Plantae - Amboropsida
Plantae - Dicotyledons
Plantae - Monocotyledons
Amoebozoa
Fungi - Microsporidia
Fungi - Blastocladiomycota
Fungi - Mucoromycota
Fungi - Zoopagomycota
Fungi - Ascomycota
Fungi - Basidiomycota
Filasterea
Choanoflagellates
Metazoa - Porifera
Metazoa - Placozoa
Metazoa - Cnidarians
Metazoa - Echinodermata
Metazoa - Chordata
Metazoa - Hemichordata
Metazoa - Ecdysozoa: Priapulida
Metazoa - Ecdysozoa: Nematoda
Metazoa - Ecdysozoa: Arthropoda
Metazoa - Ecdysozoa: Tardigrada
Metazoa - Rotifera
Metazoa - Lophotrochozoa: Platyhelminthes
Metazoa - Lophotrochozoa: Mollusca
Metazoa - Lophotrochozoa: Annelida
Metazoa - Lophotrochozoa: Brachiopoda

Kolobok

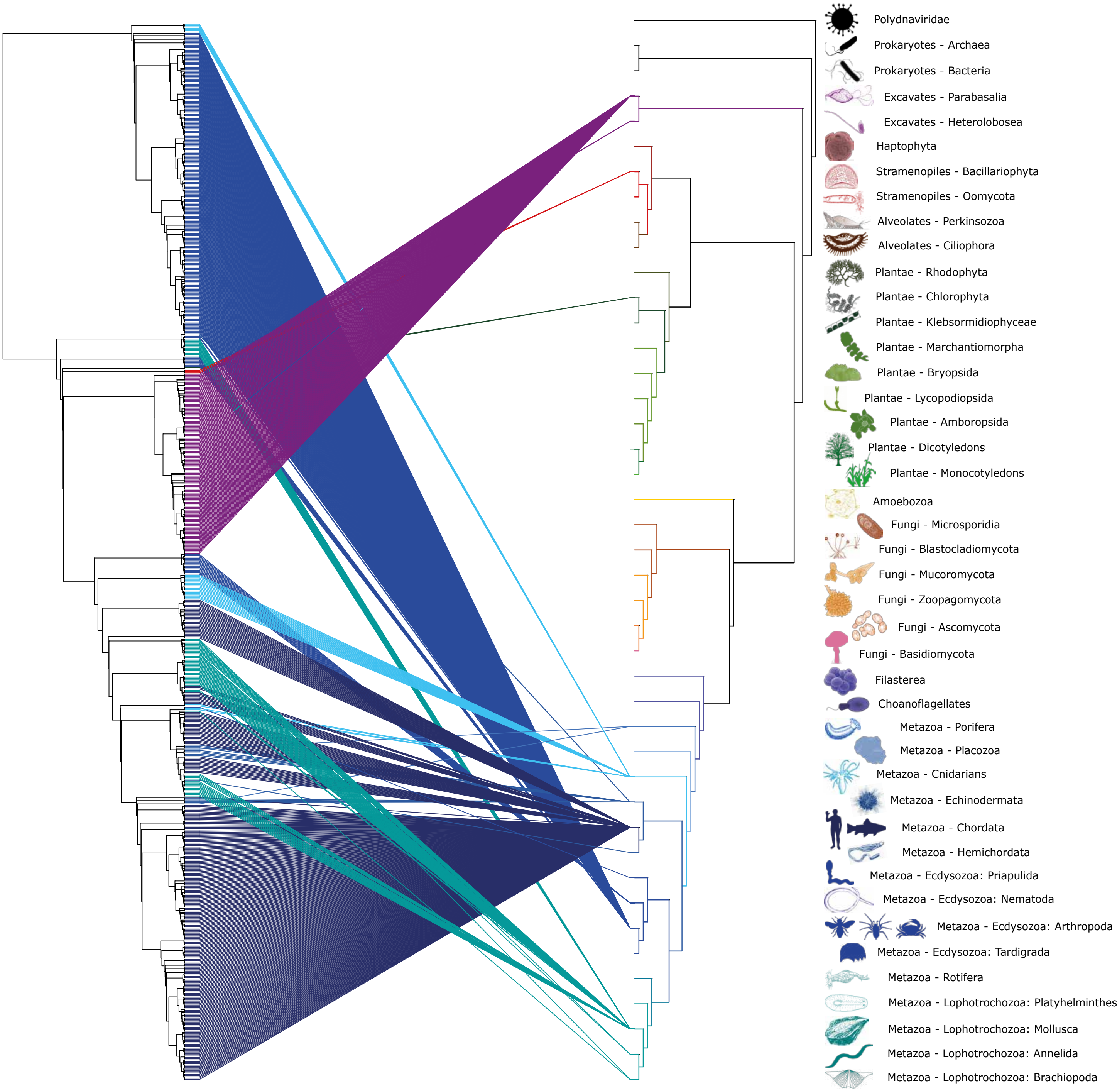

Polydnaviridae
Prokaryotes - Archaea
Prokaryotes - Bacteria
Excavates - Parabasalia
Excavates - Heterolobosea
Haptophyta
Stramenopiles - Bacillariophyta
Stramenopiles - Oomycota
Alveolates - Perkinsozoa
Alveolates - Ciliophora
Plantae - Rhodophyta
Plantae - Chlorophyta
Plantae - Klebsormidiophyceae
Plantae - Marchantiomorpha
Plantae - Bryopsida
Plantae - Lycopodiopsida
Plantae - Amboropsida
Plantae - Dicotyledons
Plantae - Monocotyledons
Amoebozoa
Fungi - Microsporidia
Fungi - Blastocladiomycota
Fungi - Mucoromycota
Fungi - Zoopagomycota
Fungi - Ascomycota
Fungi - Basidiomycota
Filasterea
Choanoflagellates
Metazoa - Porifera
Metazoa - Placozoa
Metazoa - Cnidarians
Metazoa - Echinodermata
Metazoa - Chordata
Metazoa - Hemichordata
Metazoa - Ecdysozoa: Priapulida
Metazoa - Ecdysozoa: Nematoda
Metazoa - Ecdysozoa: Arthropoda
Metazoa - Ecdysozoa: Tardigrada
Metazoa - Rotifera
Metazoa - Lophotrochozoa: Platyhelminthes
Metazoa - Lophotrochozoa: Mollusca
Metazoa - Lophotrochozoa: Annelida
Metazoa - Lophotrochozoa: Brachiopoda

### Merlin

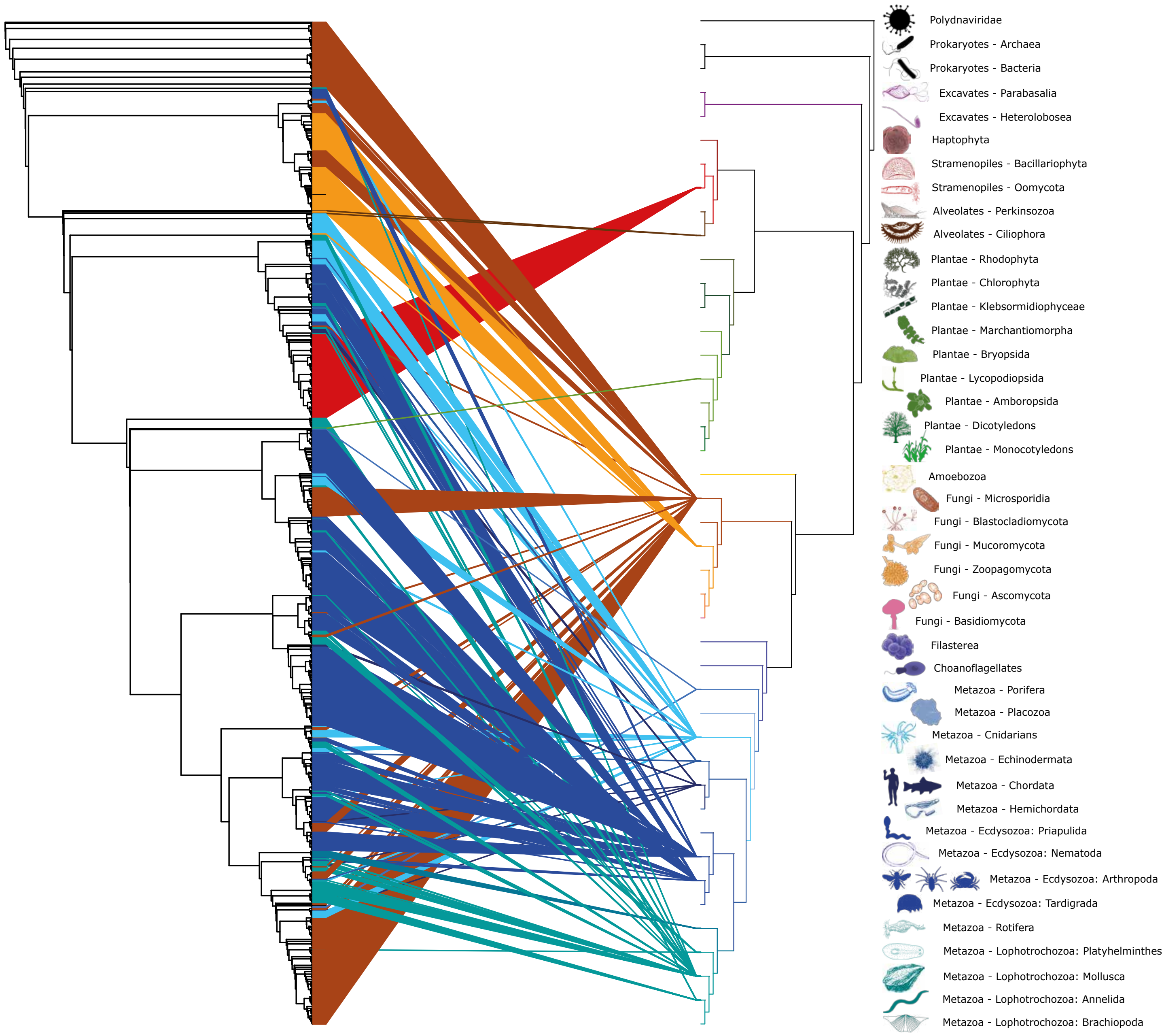

Polydnaviridae
Prokaryotes - Archaea
Prokaryotes - Bacteria
Excavates - Parabasalia
Excavates - Heterolobosea
Haptophyta
Stramenopiles - Bacillariophyta
Stramenopiles - Oomycota
Alveolates - Perkinsozoa
Alveolates - Ciliophora
Plantae - Rhodophyta
Plantae - Chlorophyta
Plantae - Klebsormidiophyceae
Plantae - Marchantiomorpha
Plantae - Bryopsida
Plantae - Lycopodiopsida
Plantae - Amboropsida
Plantae - Dicotyledons
Plantae - Monocotyledons
Amoebozoa
Fungi - Microsporidia
Fungi - Blastocladiomycota
Fungi - Mucoromycota
Fungi - Zoopagomycota
Fungi - Ascomycota
Fungi - Basidiomycota
Filasterea
Choanoflagellates
Metazoa - Porifera
Metazoa - Placozoa
Metazoa - Cnidarians
Metazoa - Echinodermata
Metazoa - Chordata
Metazoa - Hemichordata
Metazoa - Ecdysozoa: Priapulida
Metazoa - Ecdysozoa: Nematoda
Metazoa - Ecdysozoa: Arthropoda
Metazoa - Ecdysozoa: Tardigrada
Metazoa - Rotifera
Metazoa - Lophotrochozoa: Platyhelminthes
Metazoa - Lophotrochozoa: Mollusca
Metazoa - Lophotrochozoa: Annelida
Metazoa - Lophotrochozoa: Brachiopoda

### Mutators

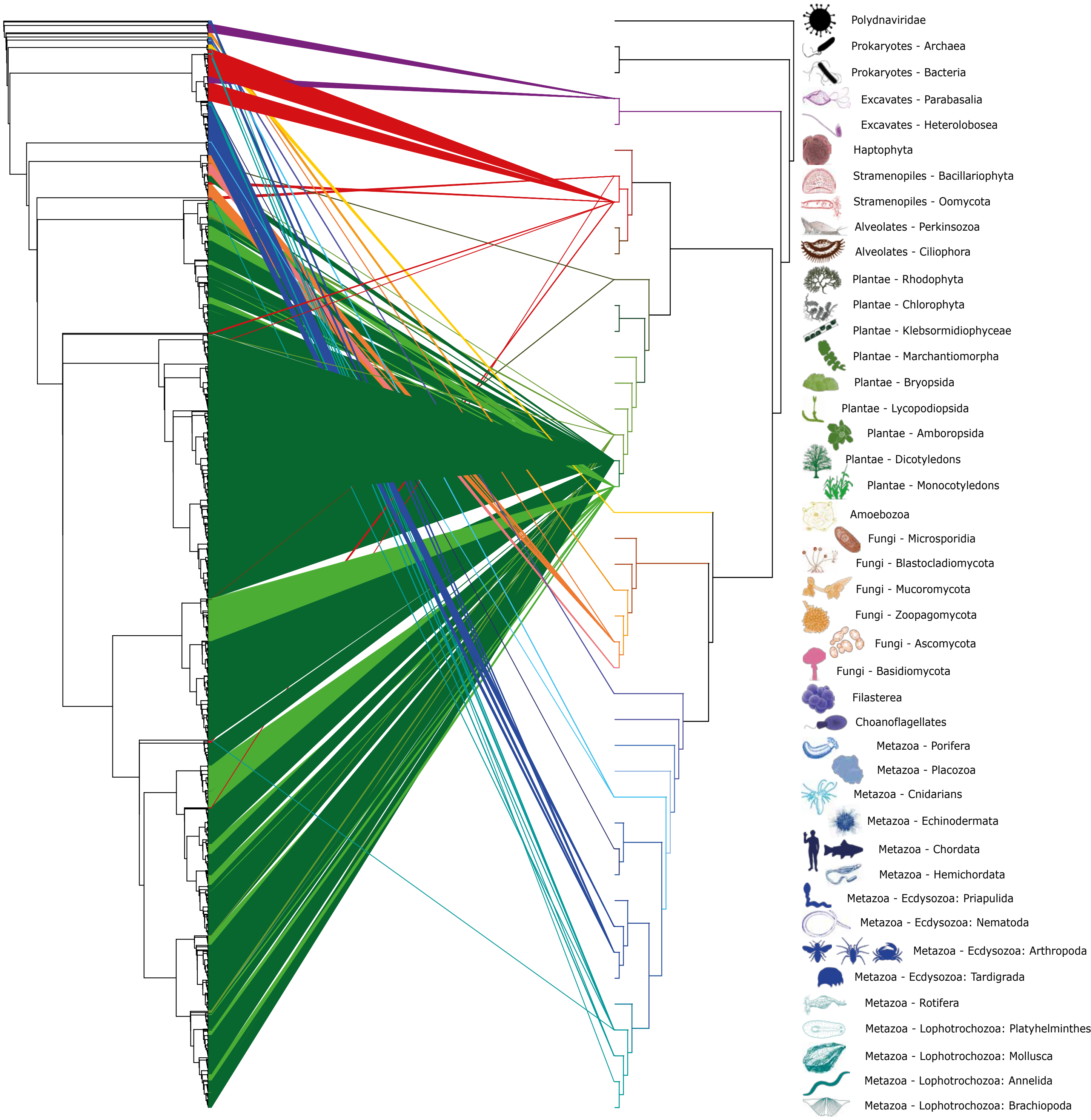
Polydnaviridae
Prokaryotes - Archaea
Prokaryotes - Bacteria
Excavates - Parabasalia
Excavates - Heterolobosea
Haptophyta
Stramenopiles - Bacillariophyta
Stramenopiles - Oomycota
Alveolates - Perkinsozoa
Alveolates - Ciliophora
Plantae - Rhodophyta
Plantae - Chlorophyta
Plantae - Klebsormidiophyceae
Plantae - Marchantiomorpha
Plantae - Bryopsida
Plantae - Lycopodiopsida
Plantae - Amboropsida
Plantae - Dicotyledons
Plantae - Monocotyledons
Amoebozoa
Fungi - Microsporidia
Fungi - Blastocladiomycota
Fungi - Mucoromycota
Fungi - Zoopagomycota
Fungi - Ascomycota
Fungi - Basidiomycota
Filasterea
Choanoflagellates
Metazoa - Porifera
Metazoa - Placozoa
Metazoa - Cnidarians
Metazoa - Echinodermata
Metazoa - Chordata
Metazoa - Hemichordata
Metazoa - Ecdysozoa: Priapulida
Metazoa - Ecdysozoa: Nematoda
Metazoa - Ecdysozoa: Arthropoda
Metazoa - Ecdysozoa: Tardigrada
Metazoa - Rotifera
Metazoa - Lophotrochozoa: Platyhelminthes
Metazoa - Lophotrochozoa: Mollusca
Metazoa - Lophotrochozoa: Annelida
Metazoa - Lophotrochozoa: Brachiopoda

Novosib

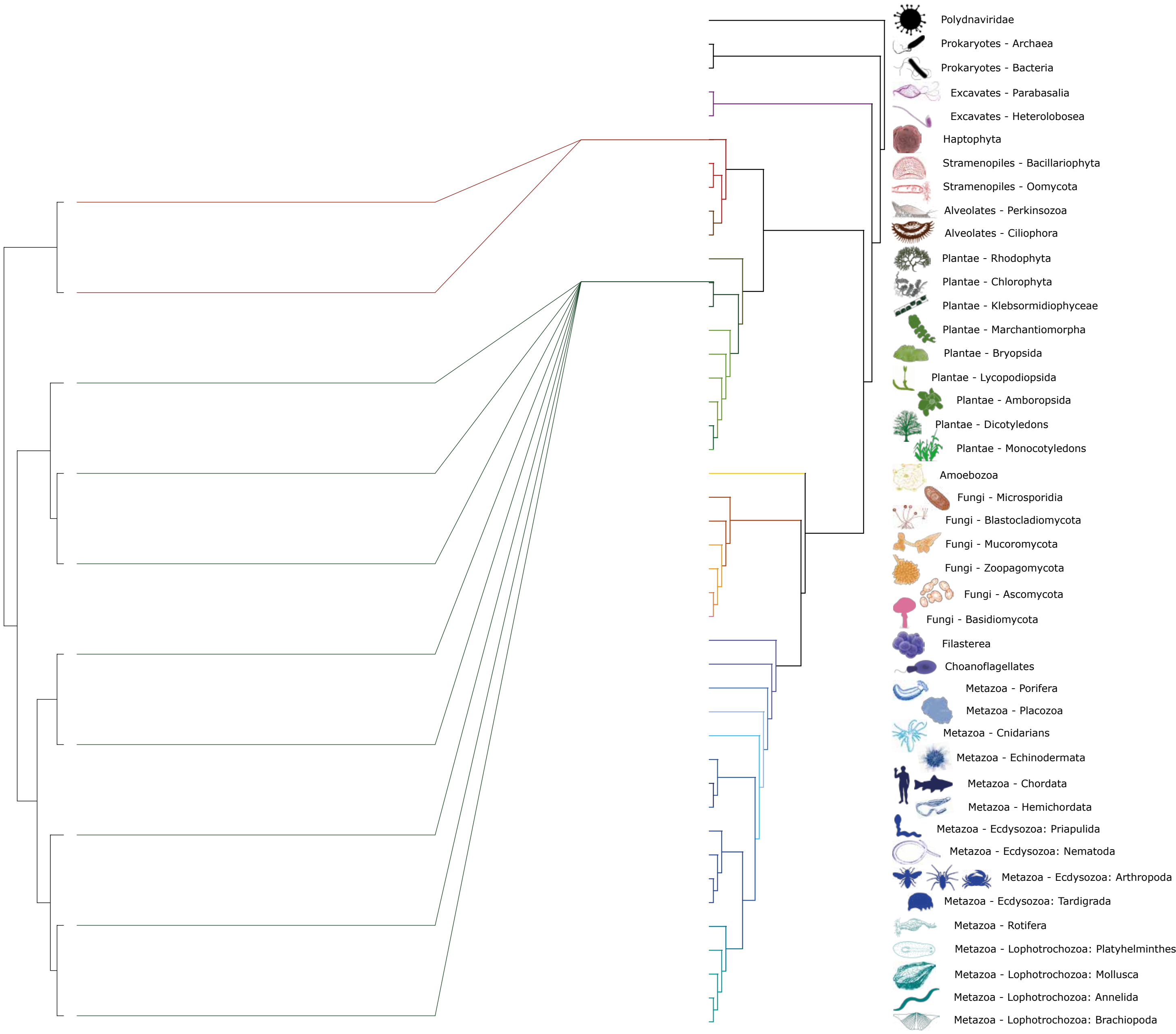
Polydnaviridae
Prokaryotes - Archaea
Prokaryotes - Bacteria
Excavates - Parabasalia
Excavates - Heterolobosea
Haptophyta
Stramenopiles - Bacillariophyta
Stramenopiles - Oomycota
Alveolates - Perkinsozoa
Alveolates - Ciliophora
Plantae - Rhodophyta
Plantae - Chlorophyta
Plantae - Klebsormidiophyceae
Plantae - Marchantiomorpha
Plantae - Bryopsida
Plantae - Lycopodiopsida
Plantae - Amboropsida
Plantae - Dicotyledons
Plantae - Monocotyledons
Amoebozoa
Fungi - Microsporidia
Fungi - Blastocladiomycota
Fungi - Mucoromycota
Fungi - Zoopagomycota
Fungi - Ascomycota
Fungi - Basidiomycota
Filasterea
Choanoflagellates
Metazoa - Porifera
Metazoa - Placozoa
Metazoa - Cnidarians
Metazoa - Echinodermata
Metazoa - Chordata
Metazoa - Hemichordata
Metazoa - Ecdysozoa: Priapulida
Metazoa - Ecdysozoa: Nematoda
Metazoa - Ecdysozoa: Arthropoda
Metazoa - Ecdysozoa: Tardigrada
Metazoa - Rotifera
Metazoa - Lophotrochozoa: Platyhelminthes
Metazoa - Lophotrochozoa: Mollusca
Metazoa - Lophotrochozoa: Annelida
Metazoa - Lophotrochozoa: Brachiopoda

P

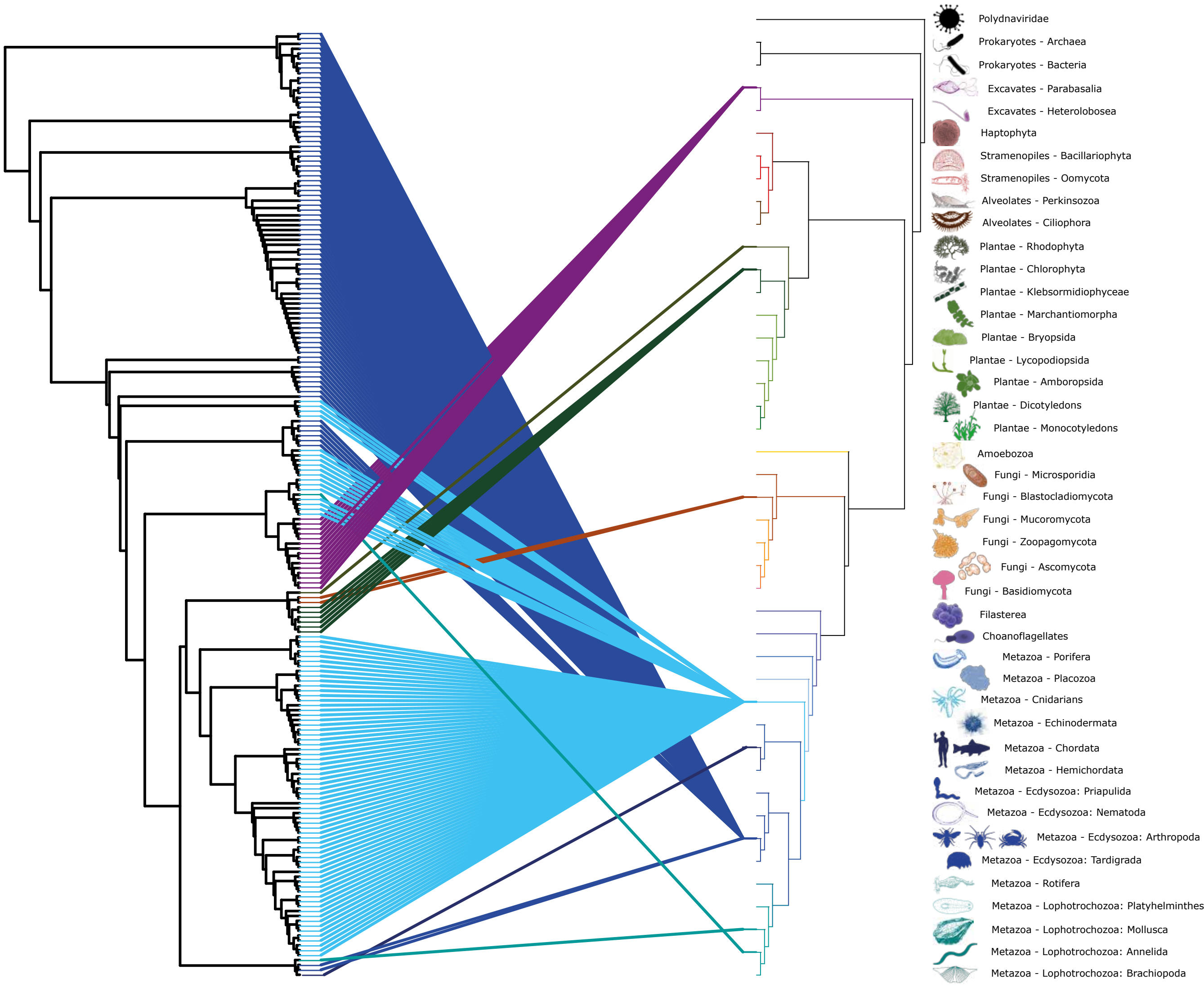
Polydnaviridae
Prokaryotes - Archaea
Prokaryotes - Bacteria
Excavates - Parabasalia
Excavates - Heterolobosea
Haptophyta
Stramenopiles - Bacillariophyta
Stramenopiles - Oomycota
Alveolates - Perkinsozoa
Alveolates - Ciliophora
Plantae - Rhodophyta
Plantae - Chlorophyta
Plantae - Klebsormidiophyceae
Plantae - Marchantiomorpha
Plantae - Bryopsida
Plantae - Lycopodiopsida
Plantae - Amboropsida
Plantae - Dicotyledons
Plantae - Monocotyledons
Amoebozoa
Fungi - Microsporidia
Fungi - Blastocladiomycota
Fungi - Mucoromycota
Fungi - Zoopagomycota
Fungi - Ascomycota
Fungi - Basidiomycota
Filasterea
Choanoflagellates
Metazoa - Porifera
Metazoa - Placozoa
Metazoa - Cnidarians
Metazoa - Echinodermata
Metazoa - Chordata
Metazoa - Hemichordata
Metazoa - Ecdysozoa: Priapulida
Metazoa - Ecdysozoa: Nematoda
Metazoa - Ecdysozoa: Arthropoda
Metazoa - Ecdysozoa: Tardigrada
Metazoa - Rotifera
Metazoa - Lophotrochozoa: Platyhelminthes
Metazoa - Lophotrochozoa: Mollusca
Metazoa - Lophotrochozoa: Annelida
Metazoa - Lophotrochozoa: Brachiopoda

### PHIS

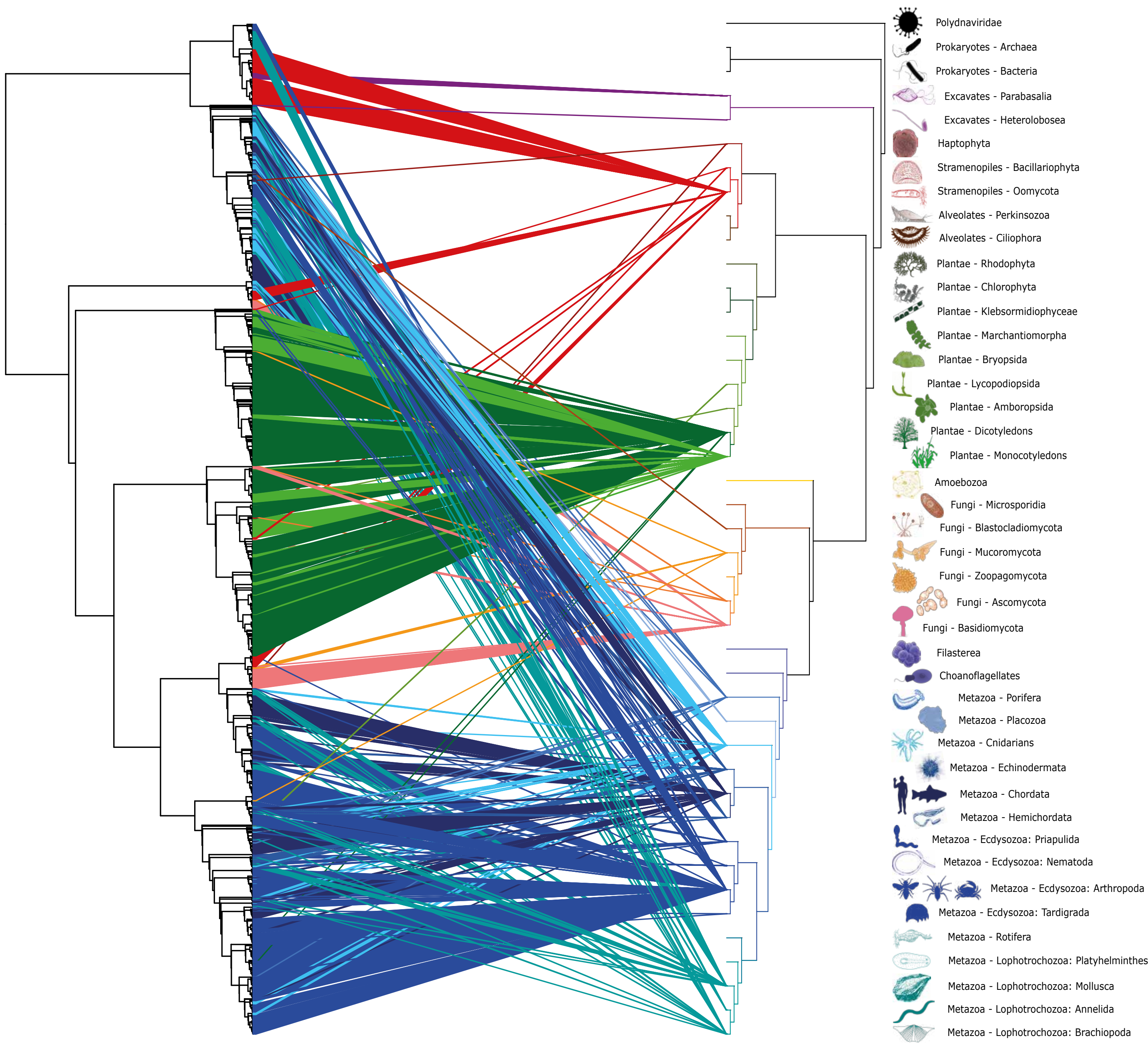

Polydnaviridae
Prokaryotes - Archaea
Prokaryotes - Bacteria
Excavates - Parabasalia
Excavates - Heterolobosea
Haptophyta
Stramenopiles - Bacillariophyta
Stramenopiles - Oomycota
Alveolates - Perkinsozoa
Alveolates - Ciliophora
Plantae - Rhodophyta
Plantae - Chlorophyta
Plantae - Klebsormidiophyceae
Plantae - Marchantiomorpha
Plantae - Bryopsida
Plantae - Lycopodiopsida
Plantae - Amboropsida
Plantae - Dicotyledons
Plantae - Monocotyledons
Amoebozoa
Fungi - Microsporidia
Fungi - Blastocladiomycota
Fungi - Mucoromycota
Fungi - Zoopagomycota
Fungi - Ascomycota
Fungi - Basidiomycota
Filasterea
Choanoflagellates
Metazoa - Porifera
Metazoa - Placozoa
Metazoa - Cnidarians
Metazoa - Echinodermata
Metazoa - Chordata
Metazoa - Hemichordata
Metazoa - Ecdysozoa: Priapulida
Metazoa - Ecdysozoa: Nematoda
Metazoa - Ecdysozoa: Arthropoda
Metazoa - Ecdysozoa: Tardigrada
Metazoa - Rotifera
Metazoa - Lophotrochozoa: Platyhelminthes
Metazoa - Lophotrochozoa: Mollusca
Metazoa - Lophotrochozoa: Annelida
Metazoa - Lophotrochozoa: Brachiopoda

### PiggyBac

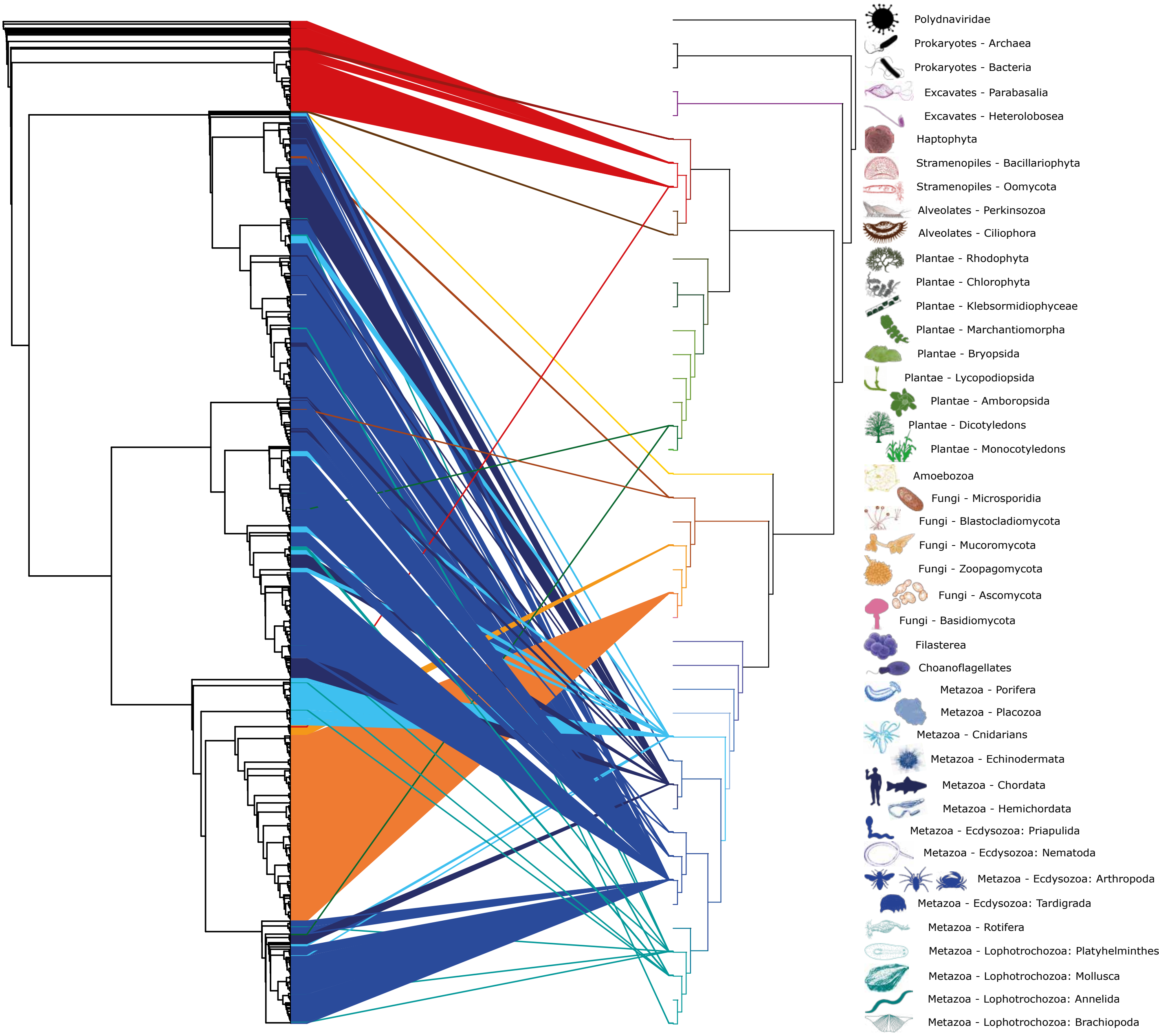
Polydnaviridae
Prokaryotes - Archaea
Prokaryotes - Bacteria
Excavates - Parabasalia
Excavates - Heterolobosea
Haptophyta
Stramenopiles - Bacillariophyta
Stramenopiles - Oomycota
Alveolates - Perkinsozoa
Alveolates - Ciliophora
Plantae - Rhodophyta
Plantae - Chlorophyta
Plantae - Klebsormidiophyceae
Plantae - Marchantiomorpha
Plantae - Bryopsida
Plantae - Lycopodiopsida
Plantae - Amboropsida
Plantae - Dicotyledons
Plantae - Monocotyledons
Amoebozoa
Fungi - Microsporidia
Fungi - Blastocladiomycota
Fungi - Mucoromycota
Fungi - Zoopagomycota
Fungi - Ascomycota
Fungi - Basidiomycota
Filasterea
Choanoflagellates
Metazoa - Porifera
Metazoa - Placozoa
Metazoa - Cnidarians
Metazoa - Echinodermata
Metazoa - Chordata
Metazoa - Hemichordata
Metazoa - Ecdysozoa: Priapulida
Metazoa - Ecdysozoa: Nematoda
Metazoa - Ecdysozoa: Arthropoda
Metazoa - Ecdysozoa: Tardigrada
Metazoa - Rotifera
Metazoa - Lophotrochozoa: Platyhelminthes
Metazoa - Lophotrochozoa: Mollusca
Metazoa - Lophotrochozoa: Annelida
Metazoa - Lophotrochozoa: Brachiopoda

### Pogo

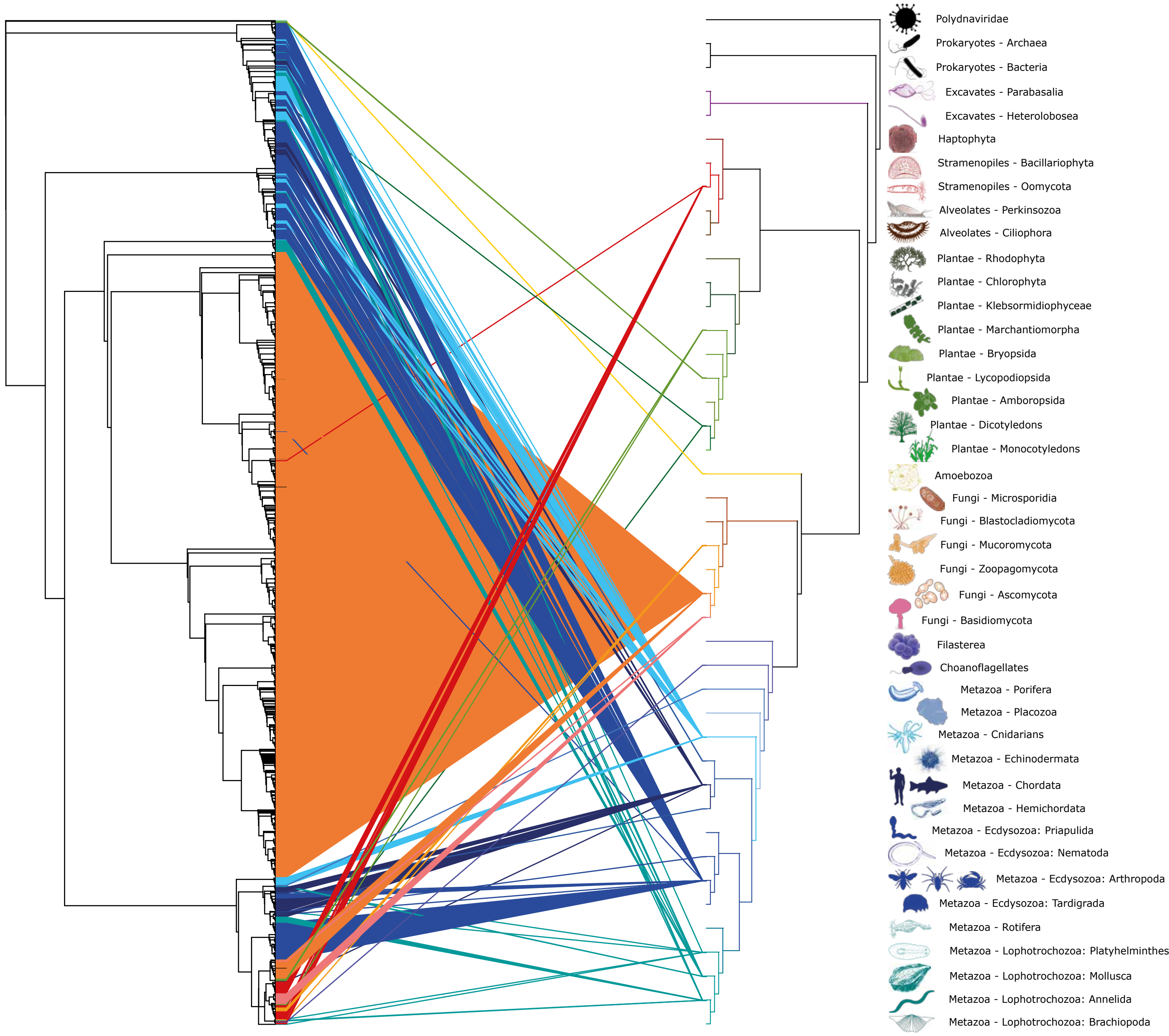

Polydnaviridae
Prokaryotes - Archaea
Prokaryotes - Bacteria
Excavates - Parabasalia
Excavates - Heterolobosea
Haptophyta
Stramenopiles - Bacillariophyta
Stramenopiles - Oomycota
Alveolates - Perkinsozoa
Alveolates - Ciliophora
Plantae - Rhodophyta
Plantae - Chlorophyta
Plantae - Klebsormidiophyceae
Plantae - Marchantiomorpha
Plantae - Bryopsida
Plantae - Lycopodiopsida
Plantae - Amboropsida
Plantae - Dicotyledons
Plantae - Monocotyledons
Amoebozoa
Fungi - Microsporidia
Fungi - Blastocladiomycota
Fungi - Mucoromycota
Fungi - Zoopagomycota
Fungi - Ascomycota
Fungi - Basidiomycota
Filasterea
Choanoflagellates
Metazoa - Porifera
Metazoa - Placozoa
Metazoa - Cnidarians
Metazoa - Echinodermata
Metazoa - Chordata
Metazoa - Hemichordata
Metazoa - Ecdysozoa: Priapulida
Metazoa - Ecdysozoa: Nematoda
Metazoa - Ecdysozoa: Arthropoda
Metazoa - Ecdysozoa: Tardigrada
Metazoa - Rotifera
Metazoa - Lophotrochozoa: Platyhelminthes
Metazoa - Lophotrochozoa: Mollusca
Metazoa - Lophotrochozoa: Annelida
Metazoa - Lophotrochozoa: Brachiopoda

### Sola1

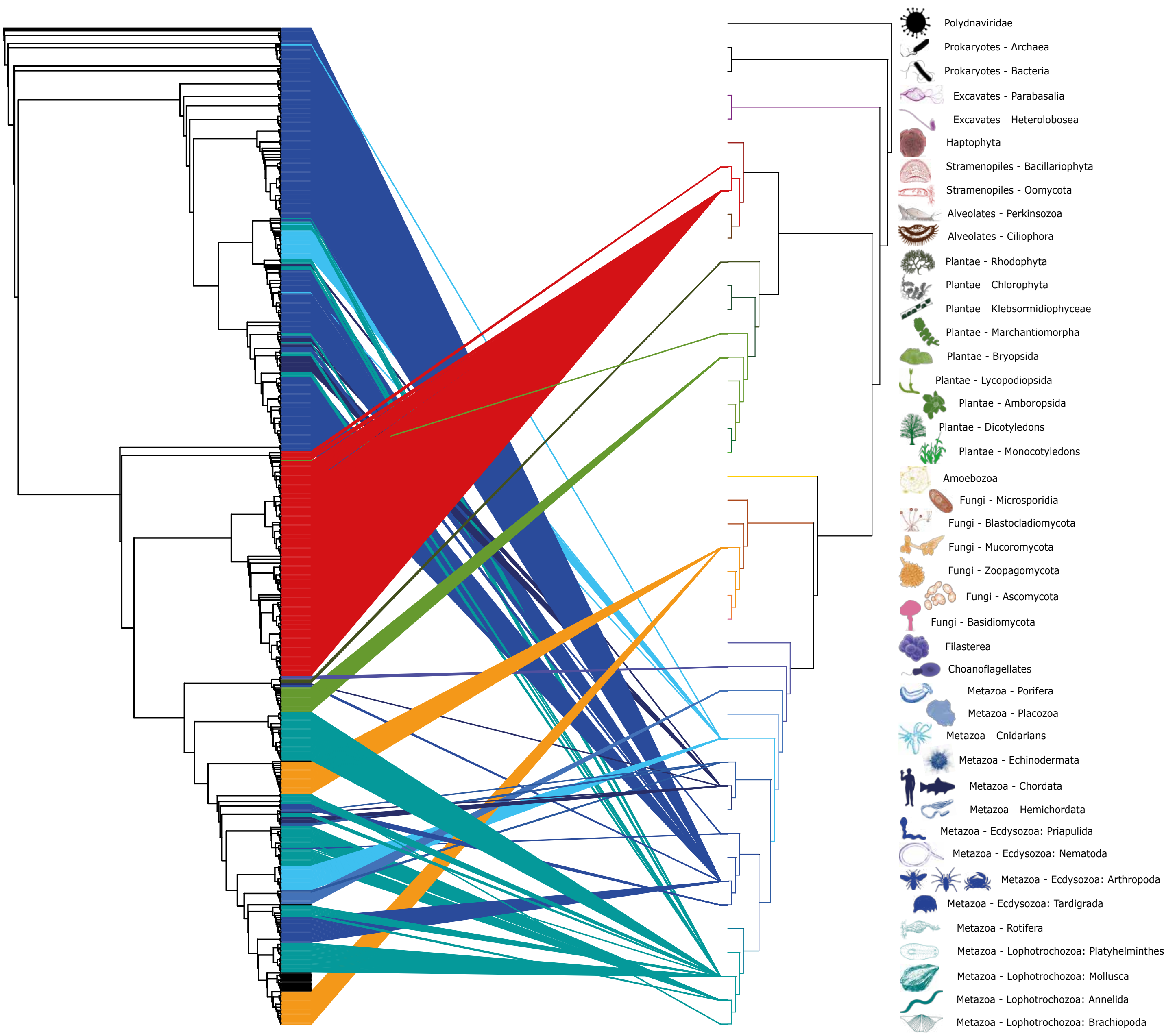

Polydnaviridae
Prokaryotes - Archaea
Prokaryotes - Bacteria
Excavates - Parabasalia
Excavates - Heterolobosea
Haptophyta
Stramenopiles - Bacillariophyta
Stramenopiles - Oomycota
Alveolates - Perkinsozoa
Alveolates - Ciliophora
Plantae - Rhodophyta
Plantae - Chlorophyta
Plantae - Klebsormidiophyceae
Plantae - Marchantiomorpha
Plantae - Bryopsida
Plantae - Lycopodiopsida
Plantae - Amboropsida
Plantae - Dicotyledons
Plantae - Monocotyledons
Amoebozoa
Fungi - Microsporidia
Fungi - Blastocladiomycota
Fungi - Mucoromycota
Fungi - Zoopagomycota
Fungi - Ascomycota
Fungi - Basidiomycota
Filasterea
Choanoflagellates
Metazoa - Porifera
Metazoa - Placozoa
Metazoa - Cnidarians
Metazoa - Echinodermata
Metazoa - Chordata
Metazoa - Hemichordata
Metazoa - Ecdysozoa: Priapulida
Metazoa - Ecdysozoa: Nematoda
Metazoa - Ecdysozoa: Arthropoda
Metazoa - Ecdysozoa: Tardigrada
Metazoa - Rotifera
Metazoa - Lophotrochozoa: Platyhelminthes
Metazoa - Lophotrochozoa: Mollusca
Metazoa - Lophotrochozoa: Annelida
Metazoa - Lophotrochozoa: Brachiopoda

Sola2

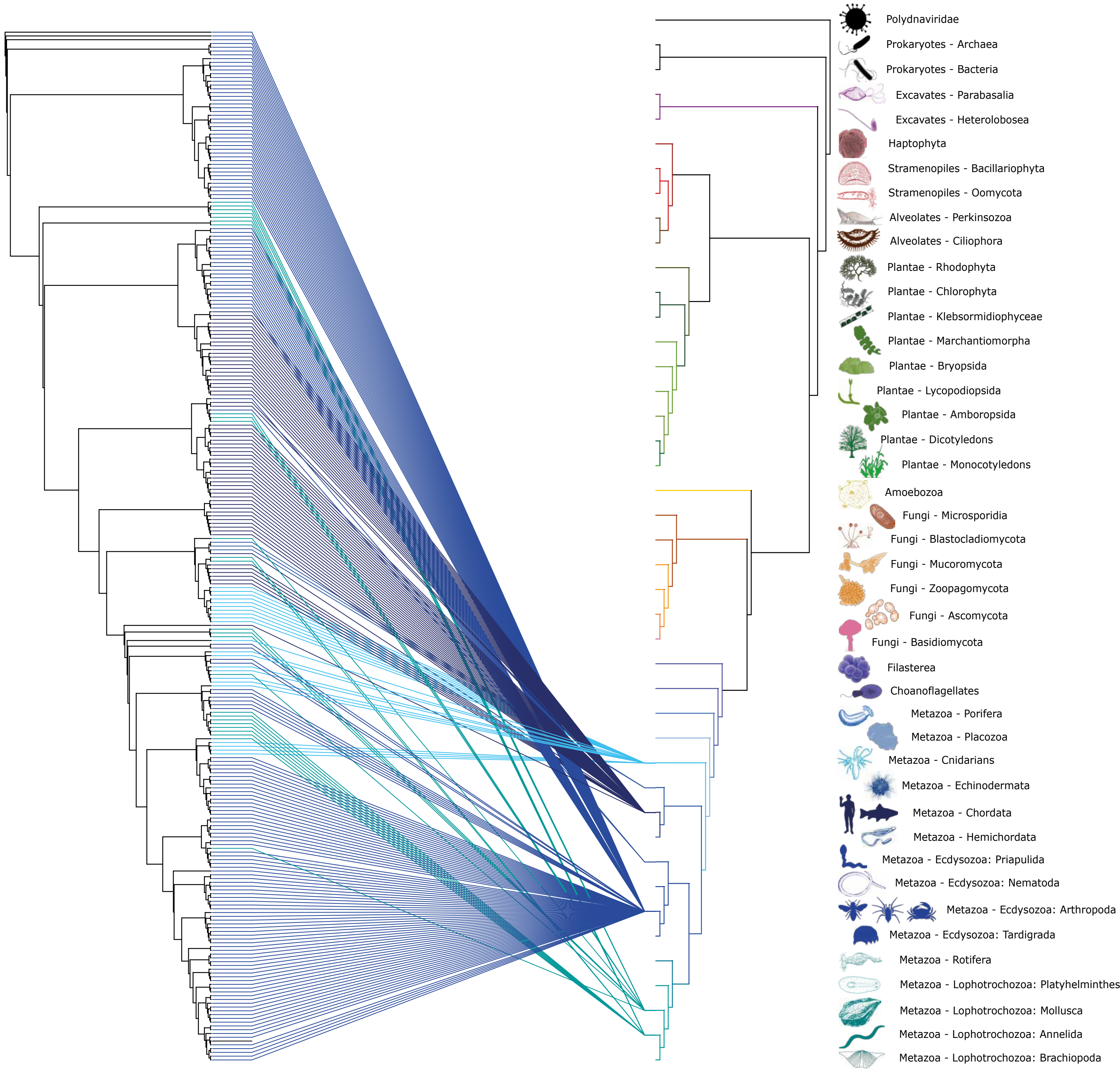
Polydnaviridae
Prokaryotes - Archaea
Prokaryotes - Bacteria
Excavates - Parabasalia
Excavates - Heterolobosea
Haptophyta
Stramenopiles - Bacillariophyta
Stramenopiles - Oomycota
Alveolates - Perkinsozoa
Alveolates - Ciliophora
Plantae - Rhodophyta
Plantae - Chlorophyta
Plantae - Klebsormidiophyceae
Plantae - Marchantiomorpha
Plantae - Bryopsida
Plantae - Lycopodiopsida
Plantae - Amboropsida
Plantae - Dicotyledons
Plantae - Monocotyledons
Amoebozoa
Fungi - Microsporidia
Fungi - Blastocladiomycota
Fungi - Mucoromycota
Fungi - Zoopagomycota
Fungi - Ascomycota
Fungi - Basidiomycota
Filasterea
Choanoflagellates
Metazoa - Porifera
Metazoa - Placozoa
Metazoa - Cnidarians
Metazoa - Echinodermata
Metazoa - Chordata
Metazoa - Hemichordata
Metazoa - Ecdysozoa: Priapulida
Metazoa - Ecdysozoa: Nematoda
Metazoa - Ecdysozoa: Arthropoda
Metazoa - Ecdysozoa: Tardigrada
Metazoa - Rotifera
Metazoa - Lophotrochozoa: Platyhelminthes
Metazoa - Lophotrochozoa: Mollusca
Metazoa - Lophotrochozoa: Annelida
Metazoa - Lophotrochozoa: Brachiopoda

### Sola3

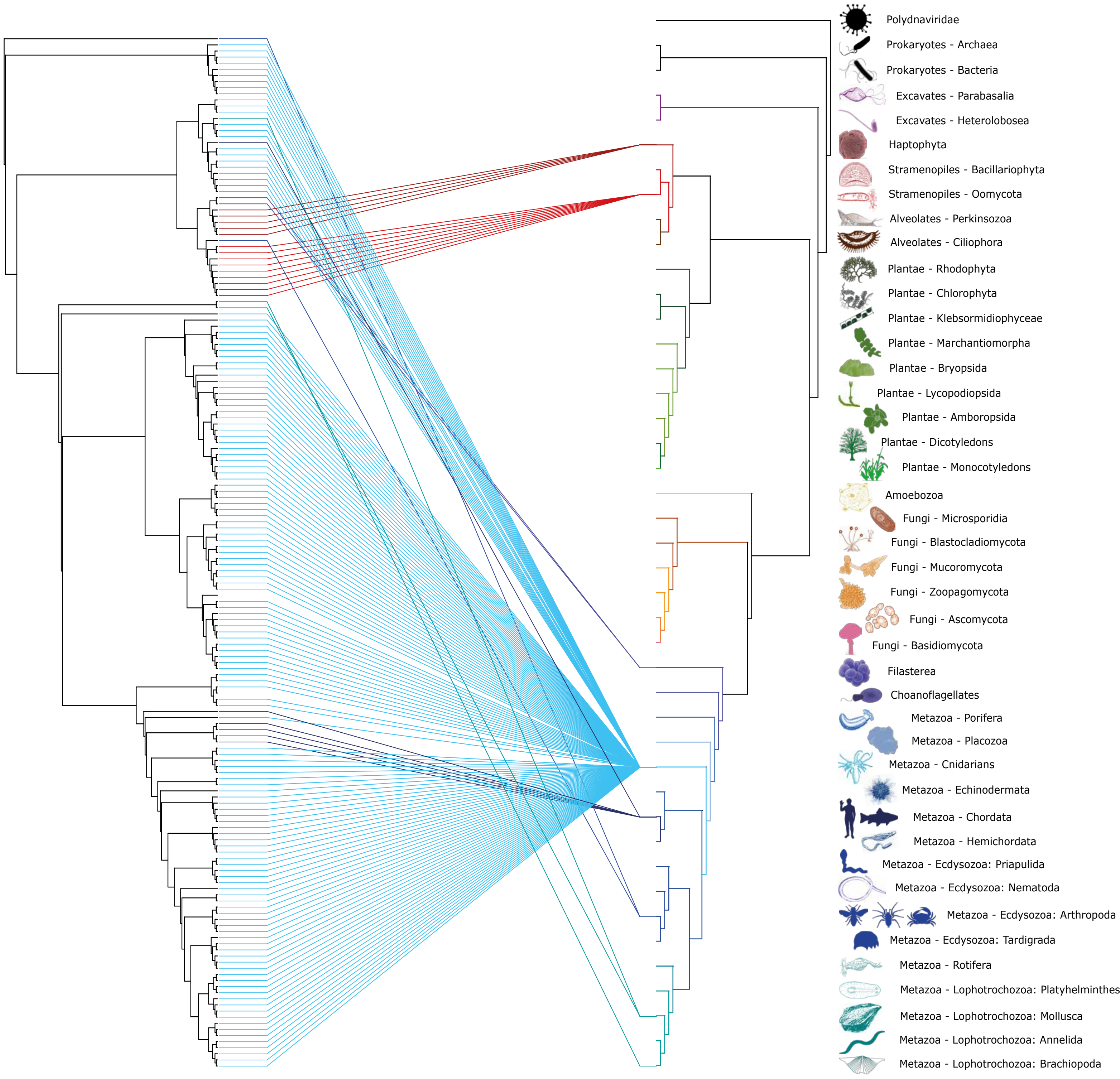

Polydnaviridae
Prokaryotes - Archaea
Prokaryotes - Bacteria
Excavates - Parabasalia
Excavates - Heterolobosea
Haptophyta
Stramenopiles - Bacillariophyta
Stramenopiles - Oomycota
Alveolates - Perkinsozoa
Alveolates - Ciliophora
Plantae - Rhodophyta
Plantae - Chlorophyta
Plantae - Klebsormidiophyceae
Plantae - Marchantiomorpha
Plantae - Bryopsida
Plantae - Lycopodiopsida
Plantae - Amboropsida
Plantae - Dicotyledons
Plantae - Monocotyledons
Amoebozoa
Fungi - Microsporidia
Fungi - Blastocladiomycota
Fungi - Mucoromycota
Fungi - Zoopagomycota
Fungi - Ascomycota
Fungi - Basidiomycota
Filasterea
Choanoflagellates
Metazoa - Porifera
Metazoa - Placozoa
Metazoa - Cnidarians
Metazoa - Echinodermata
Metazoa - Chordata
Metazoa - Hemichordata
Metazoa - Ecdysozoa: Priapulida
Metazoa - Ecdysozoa: Nematoda
Metazoa - Ecdysozoa: Arthropoda
Metazoa - Ecdysozoa: Tardigrada
Metazoa - Rotifera
Metazoa - Lophotrochozoa: Platyhelminthes
Metazoa - Lophotrochozoa: Mollusca
Metazoa - Lophotrochozoa: Annelida
Metazoa - Lophotrochozoa: Brachiopoda

### Tc1marPlm

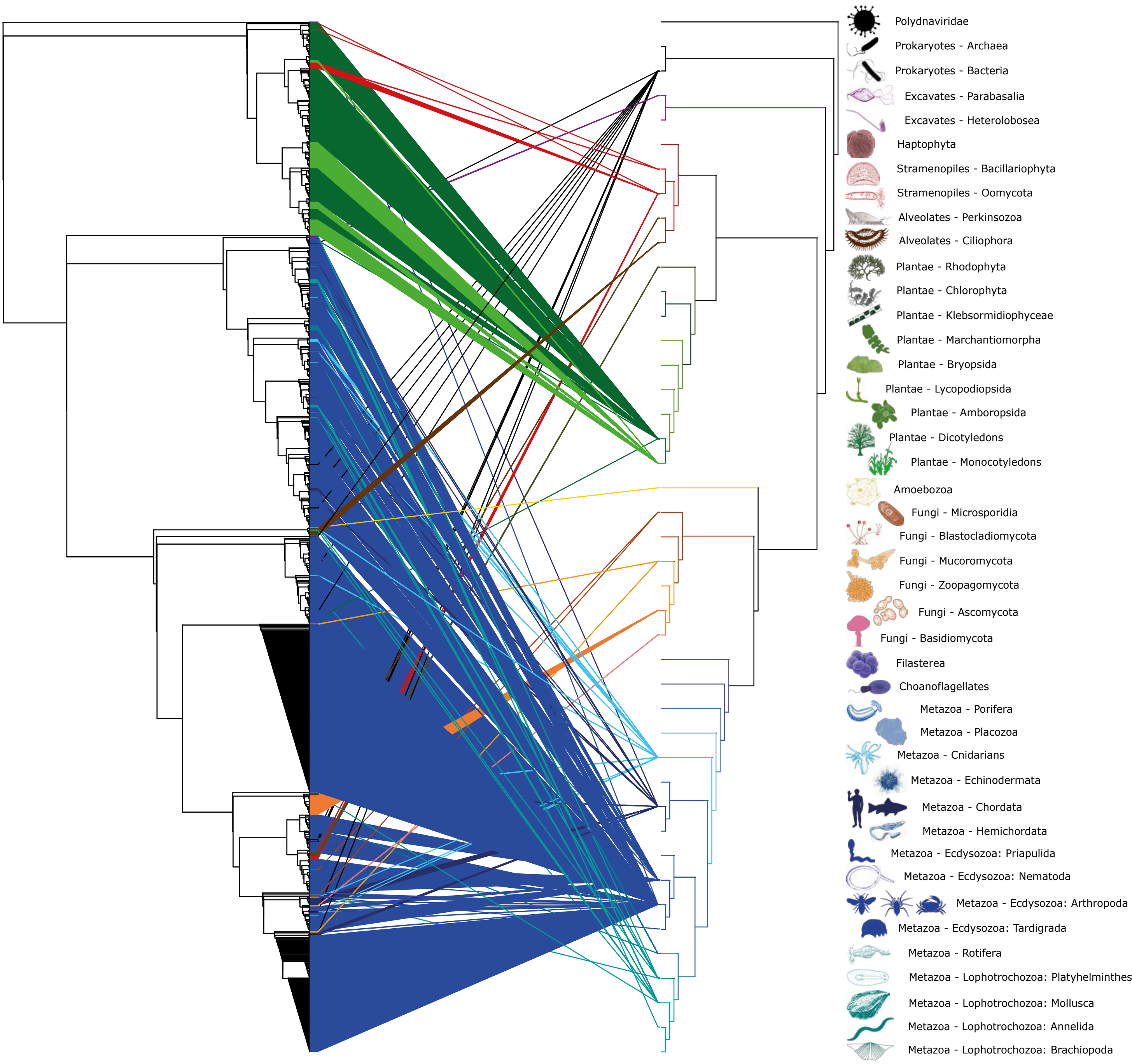
Polydnaviridae
Prokaryotes - Archaea
Prokaryotes - Bacteria
Excavates - Parabasalia
Excavates - Heterolobosea
Haptophyta
Stramenopiles - Bacillariophyta
Stramenopiles - Oomycota
Alveolates - Perkinsozoa
Alveolates - Ciliophora
Plantae - Rhodophyta
Plantae - Chlorophyta
Plantae - Klebsormidiophyceae
Plantae - Marchantiomorpha
Plantae - Bryopsida
Plantae - Lycopodiopsida
Plantae - Amboropsida
Plantae - Dicotyledons
Plantae - Monocotyledons
Amoebozoa
Fungi - Microsporidia
Fungi - Blastocladiomycota
Fungi - Mucoromycota
Fungi - Zoopagomycota
Fungi - Ascomycota
Fungi - Basidiomycota
Filasterea
Choanoflagellates
Metazoa - Porifera
Metazoa - Placozoa
Metazoa - Cnidarians
Metazoa - Echinodermata
Metazoa - Chordata
Metazoa - Hemichordata
Metazoa - Ecdysozoa: Priapulida
Metazoa - Ecdysozoa: Nematoda
Metazoa - Ecdysozoa: Arthropoda
Metazoa - Ecdysozoa: Tardigrada
Metazoa - Rotifera
Metazoa - Lophotrochozoa: Platyhelminthes
Metazoa - Lophotrochozoa: Mollusca
Metazoa - Lophotrochozoa: Annelida
Metazoa - Lophotrochozoa: Brachiopoda

### Transib

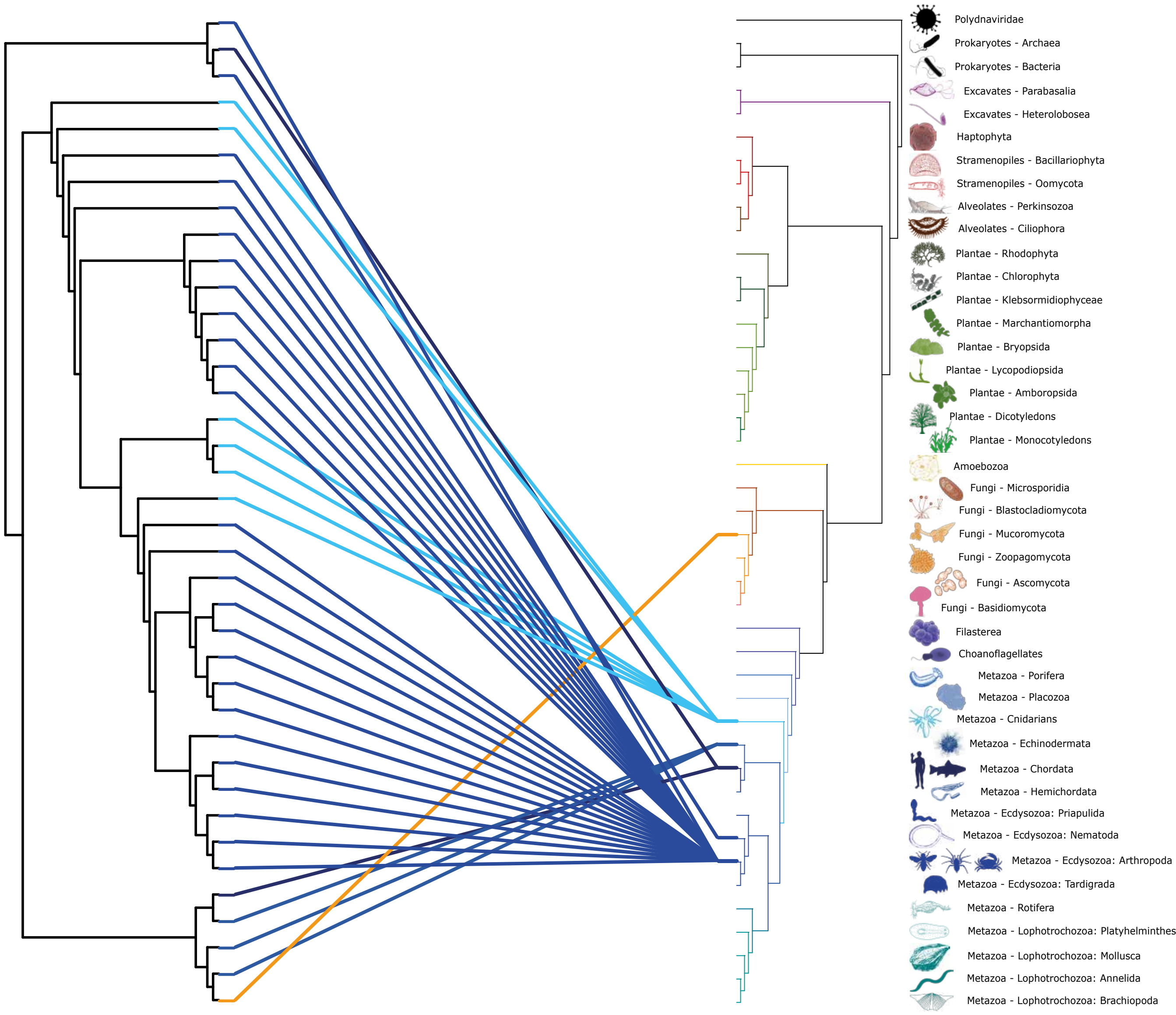

Polydnaviridae
Prokaryotes - Archaea
Prokaryotes - Bacteria
Excavates - Parabasalia
Excavates - Heterolobosea
Haptophyta
Stramenopiles - Bacillariophyta
Stramenopiles - Oomycota
Alveolates - Perkinsozoa
Alveolates - Ciliophora
Plantae - Rhodophyta
Plantae - Chlorophyta
Plantae - Klebsormidiophyceae
Plantae - Marchantiomorpha
Plantae - Bryopsida
Plantae - Lycopodiopsida
Plantae - Amboropsida
Plantae - Dicotyledons
Plantae - Monocotyledons
Amoebozoa
Fungi - Microsporidia
Fungi - Blastocladiomycota
Fungi - Mucoromycota
Fungi - Zoopagomycota
Fungi - Ascomycota
Fungi - Basidiomycota
Filasterea
Choanoflagellates
Metazoa - Porifera
Metazoa - Placozoa
Metazoa - Cnidarians
Metazoa - Echinodermata
Metazoa - Chordata
Metazoa - Hemichordata
Metazoa - Ecdysozoa: Priapulida
Metazoa - Ecdysozoa: Nematoda
Metazoa - Ecdysozoa: Arthropoda
Metazoa - Ecdysozoa: Tardigrada
Metazoa - Rotifera
Metazoa - Lophotrochozoa: Platyhelminthes
Metazoa - Lophotrochozoa: Mollusca
Metazoa - Lophotrochozoa: Annelida
Metazoa - Lophotrochozoa: Brachiopoda

Zator

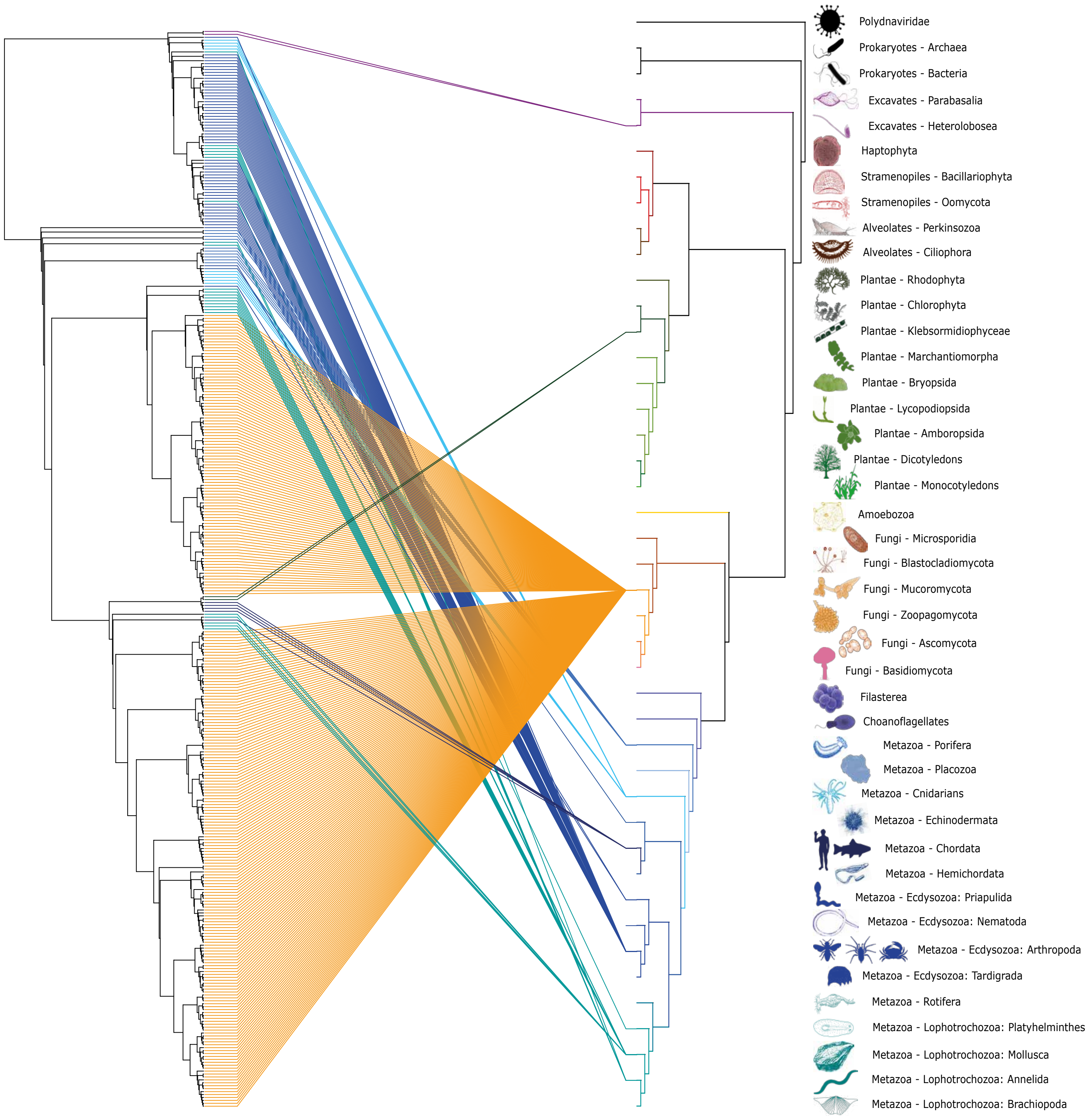
Polydnaviridae
Prokaryotes - Archaea
Prokaryotes - Bacteria
Excavates - Parabasalia
Excavates - Heterolobosea
Haptophyta
Stramenopiles - Bacillariophyta
Stramenopiles - Oomycota
Alveolates - Perkinsozoa
Alveolates - Ciliophora
Plantae - Rhodophyta
Plantae - Chlorophyta
Plantae - Klebsormidiophyceae
Plantae - Marchantiomorpha
Plantae - Bryopsida
Plantae - Lycopodiopsida
Plantae - Amboropsida
Plantae - Dicotyledons
Plantae - Monocotyledons
Amoebozoa
Fungi - Microsporidia
Fungi - Blastocladiomycota
Fungi - Mucoromycota
Fungi - Zoopagomycota
Fungi - Ascomycota
Fungi - Basidiomycota
Filasterea
Choanoflagellates
Metazoa - Porifera
Metazoa - Placozoa
Metazoa - Cnidarians
Metazoa - Echinodermata
Metazoa - Chordata
Metazoa - Hemichordata
Metazoa - Ecdysozoa: Priapulida
Metazoa - Ecdysozoa: Nematoda
Metazoa - Ecdysozoa: Arthropoda
Metazoa - Ecdysozoa: Tardigrada
Metazoa - Rotifera
Metazoa - Lophotrochozoa: Platyhelminthes
Metazoa - Lophotrochozoa: Mollusca
Metazoa - Lophotrochozoa: Annelida
Metazoa - Lophotrochozoa: Brachiopoda

Key to colours used

- Polydnaviridae, Prokaryotes - Archeae, Bacteria
- Excavates - Heterolobosea, Parabasalia
- Haptophyta
- Stramenopiles - Bacillariophyta, Oomycota
- Alveolates - Ciliophora, Perkinsozoa
- Plantae - Rodophyta
- Plantae - Chlorophyta, Klebsormidiophyceae
- Plantae - Amboropsida, Bryophyta, Lycopodiopsida, Marchantiomorpha
- Viridiplantae - Dicotyledons
- Viridiplantae - Monocotyledons
- Amoebozoa
- Fungi - Blastocladiomycota, Microsporidia
- Fungi - Mucoromycota, Zoopagomycota
- Fungi - Ascomycota
- Fungi - Basidiomycota
- Choanoflagellates, Filasterea
- Metazoa - Porifera
- Metazoa - Placozoa
- Metazoa - Cnidaria
- Metazoa - Echinodermata, Hemichordata
- Metazoa - Chordata
- Metazoa - Ecdysozoa: Nematoda, Priapulida, Panarthropoda, Tardigrada
- Metazoa - Rotifera
- Metazoa - Lophotrochozoa: Annelida, Brachiopoda, Mollusca, Platyhelminthes

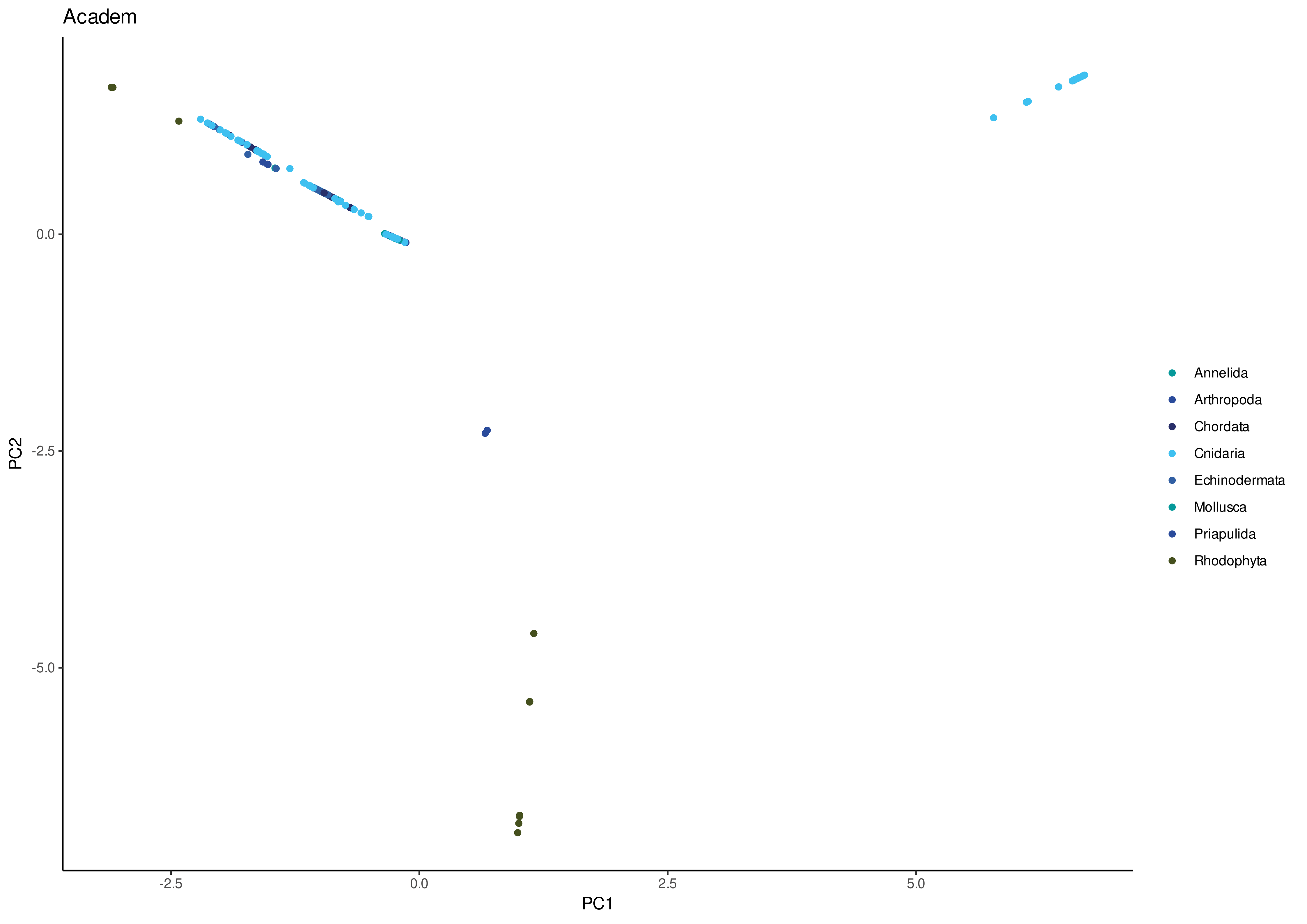
Academ
PC2
0.0
-2.5
-5.0
PC1
-2.5
0.0
2.5
5.0
Annelida
Arthropoda
Chordata
Cnidaria
Echinodermata
Mollusca
Priapulida
Rhodophyta

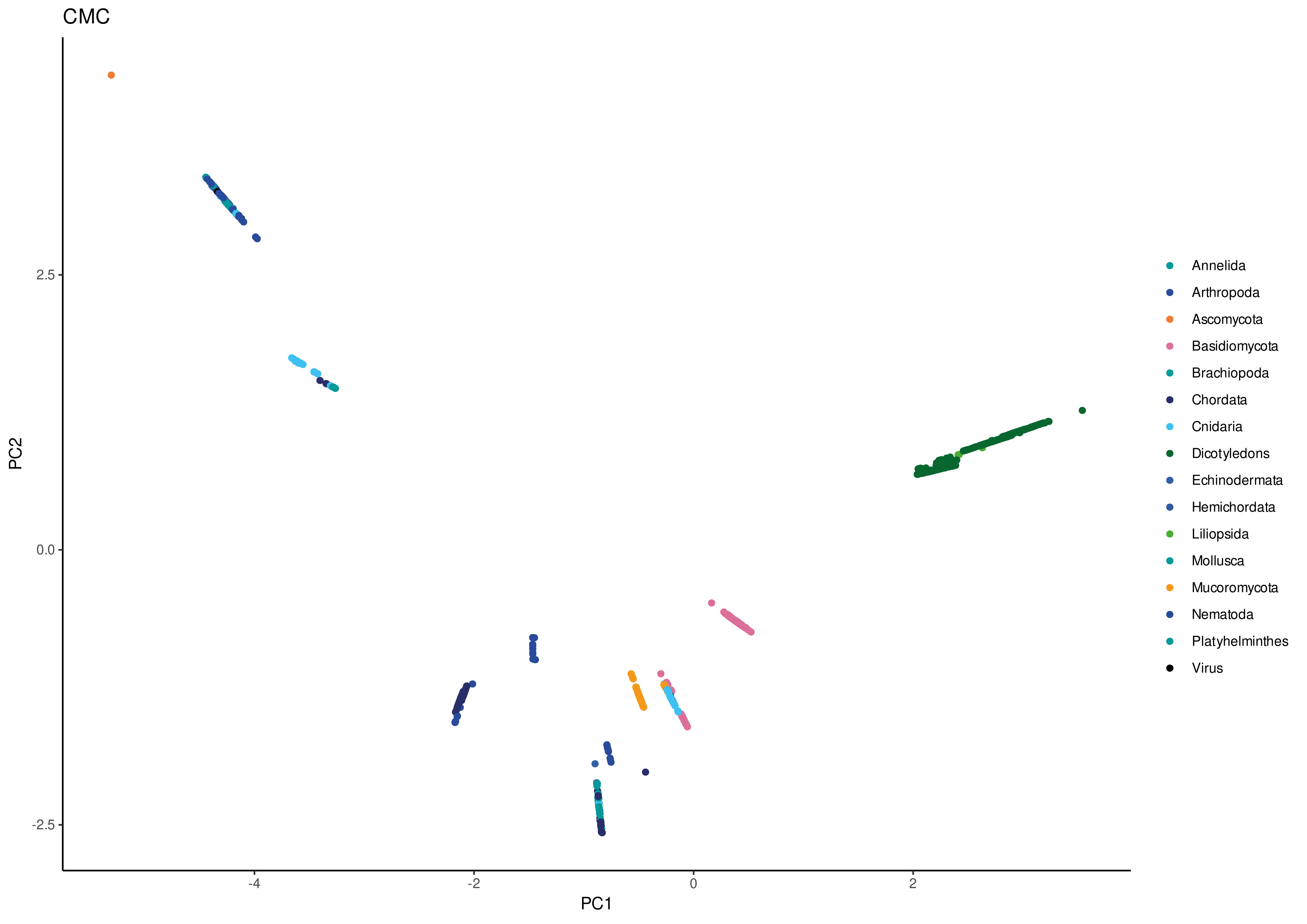
CMC
PC2
2.5
0.0
-2.5
PC1
-4
-2
0
2
Annelida
Arthropoda
Ascomycota
Basidiomycota
Brachiopoda
Chordata
Cnidaria
Dicotyledons
Echinodermata
Hemichordata
Liliopsida
Mollusca
Mucoromycota
Nematoda
Platyhelminthes
Virus

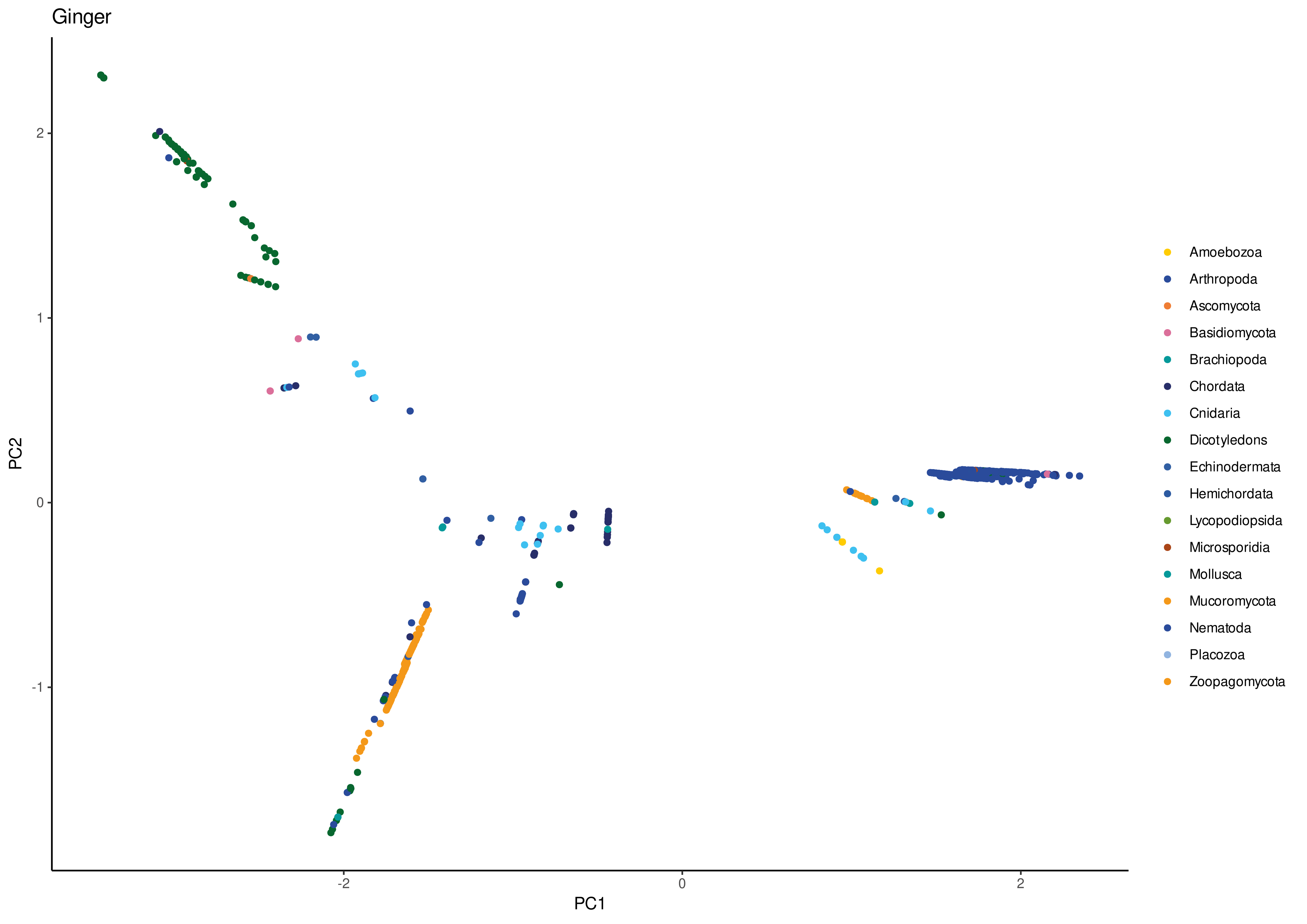
Ginger
PC2
PC1
2
1
0
-1
-2
0
2
Amoebozoa
Arthropoda
Ascomycota
Basidiomycota
Brachiopoda
Chordata
Cnidaria
Dicotyledons
Echinodermata
Hemichordata
Lycopodiopsida
Microsporidia
Mollusca
Mucoromycota
Nematoda
Placozoa
Zoopagomycota

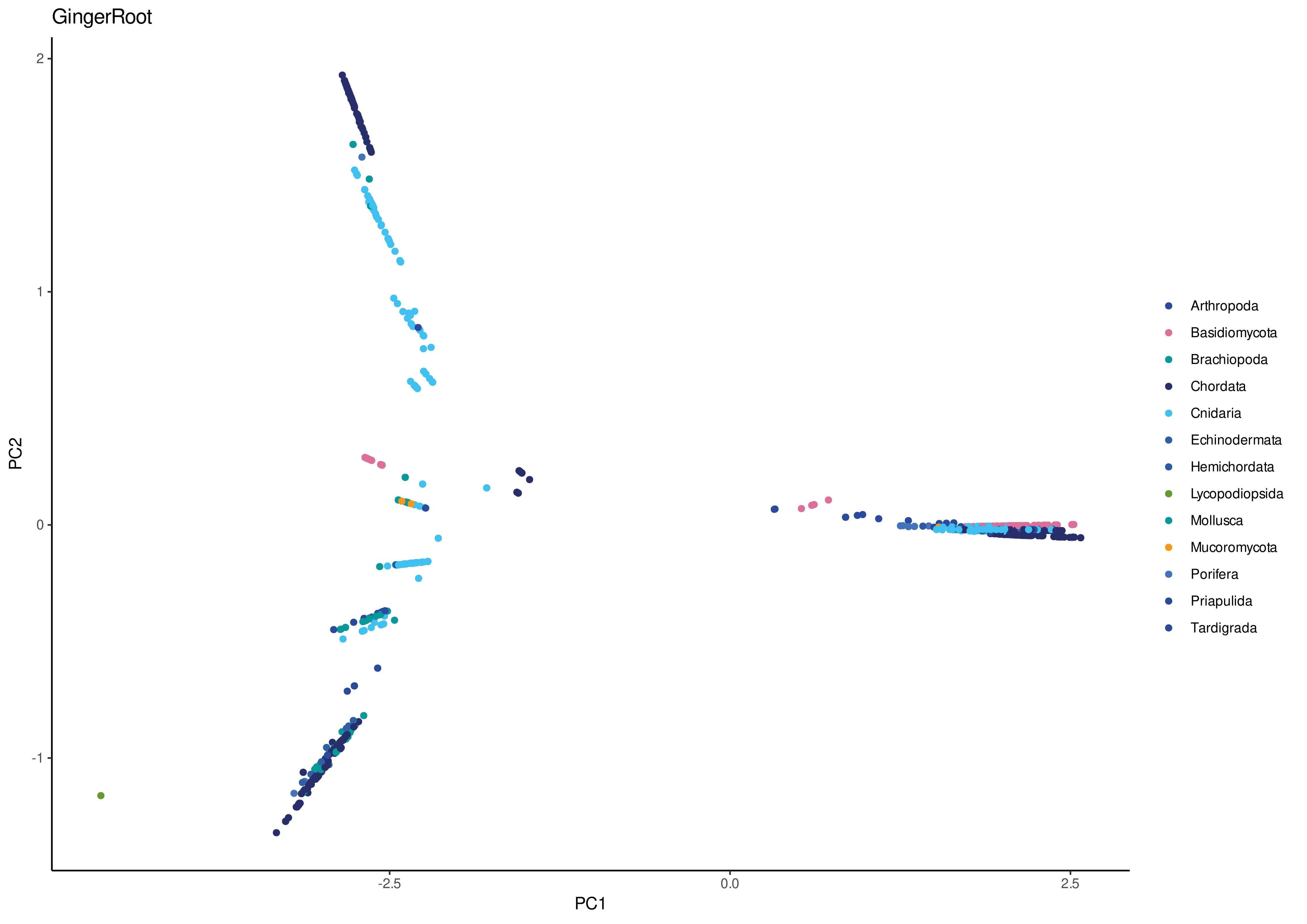
GingerRoot
PC1
PC2
-2.5
0.0
2.5
-1
0
1
2
Arthropoda
Basidiomycota
Brachiopoda
Chordata
Cnidaria
Echinodermata
Hemichordata
Lycopodiopsida
Mollusca
Mucoromycota
Porifera
Priapulida
Tardigrada

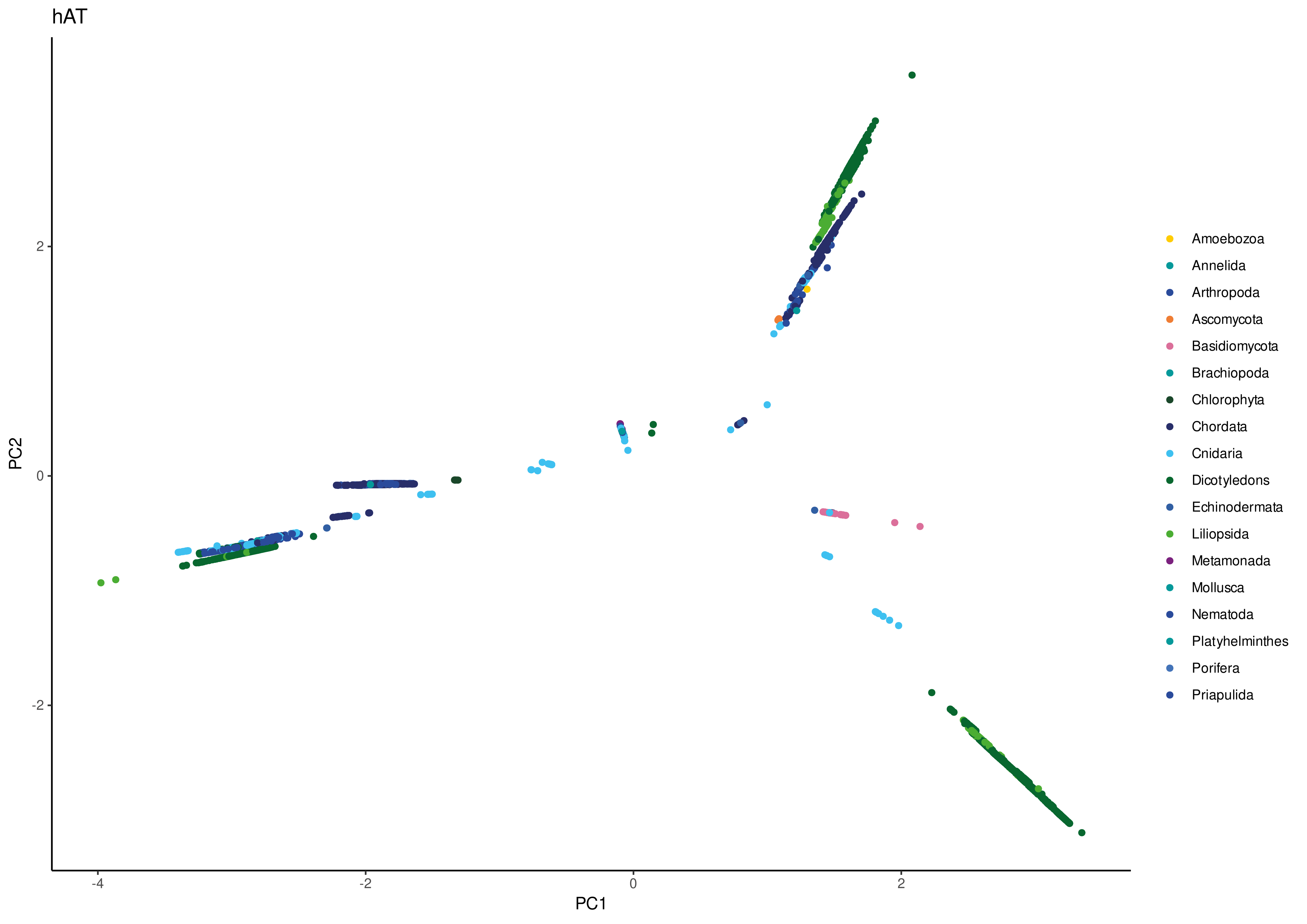
hAT
PC1
PC2
-4
-2
0
2
Amoebozoa
Annelida
Arthropoda
Ascomycota
Basidiomycota
Brachiopoda
Chlorophyta
Chordata
Cnidaria
Dicotyledons
Echinodermata
Liliopsida
Metamonada
Mollusca
Nematoda
Platyhelminthes
Porifera
Priapulida

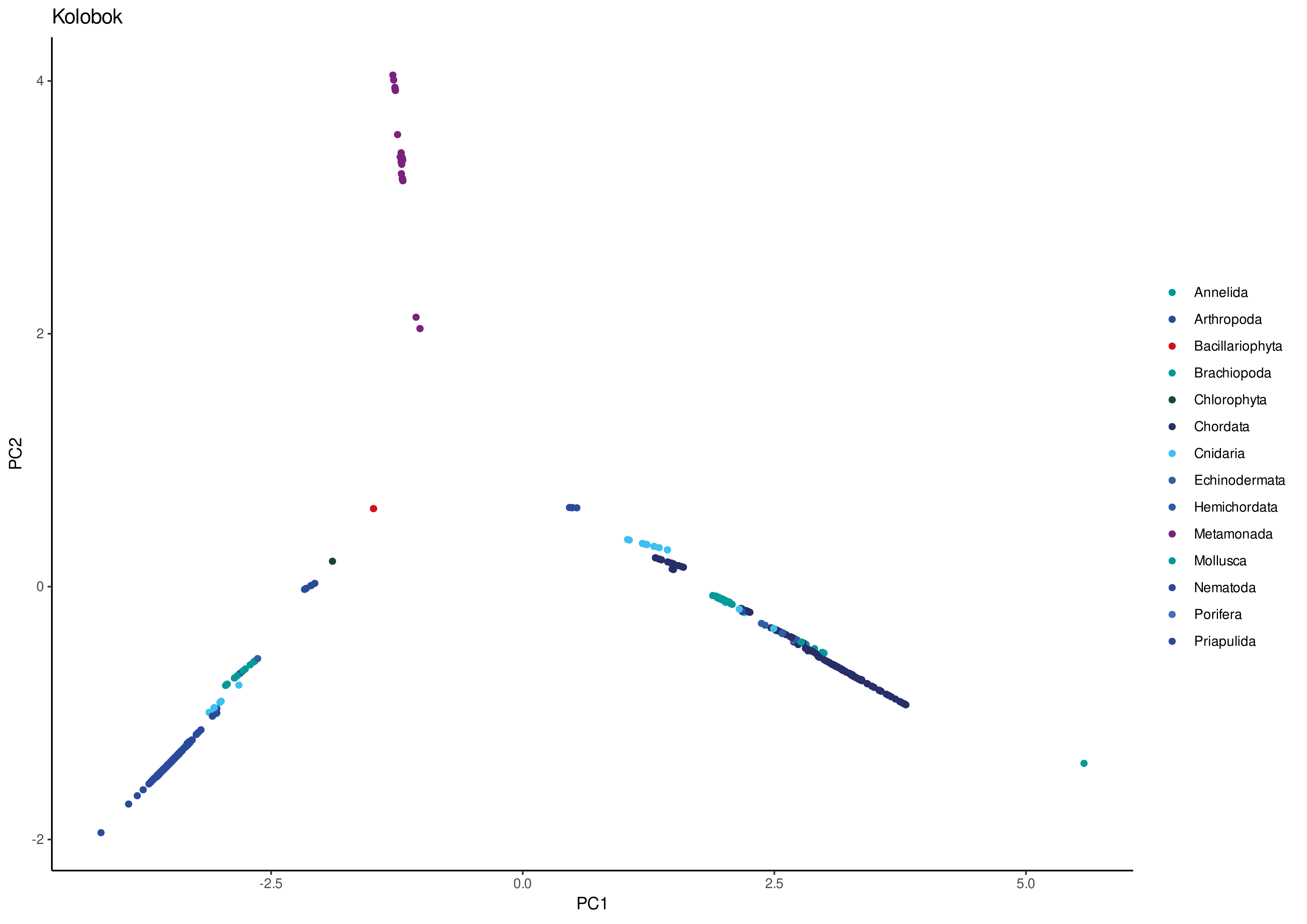
Kolobok
PC2
4
2
0
-2
PC1
-2.5
0.0
2.5
5.0
Annelida
Arthropoda
Bacillariophyta
Brachiopoda
Chlorophyta
Chordata
Cnidaria
Echinodermata
Hemichordata
Metamonada
Mollusca
Nematoda
Porifera
Priapulida

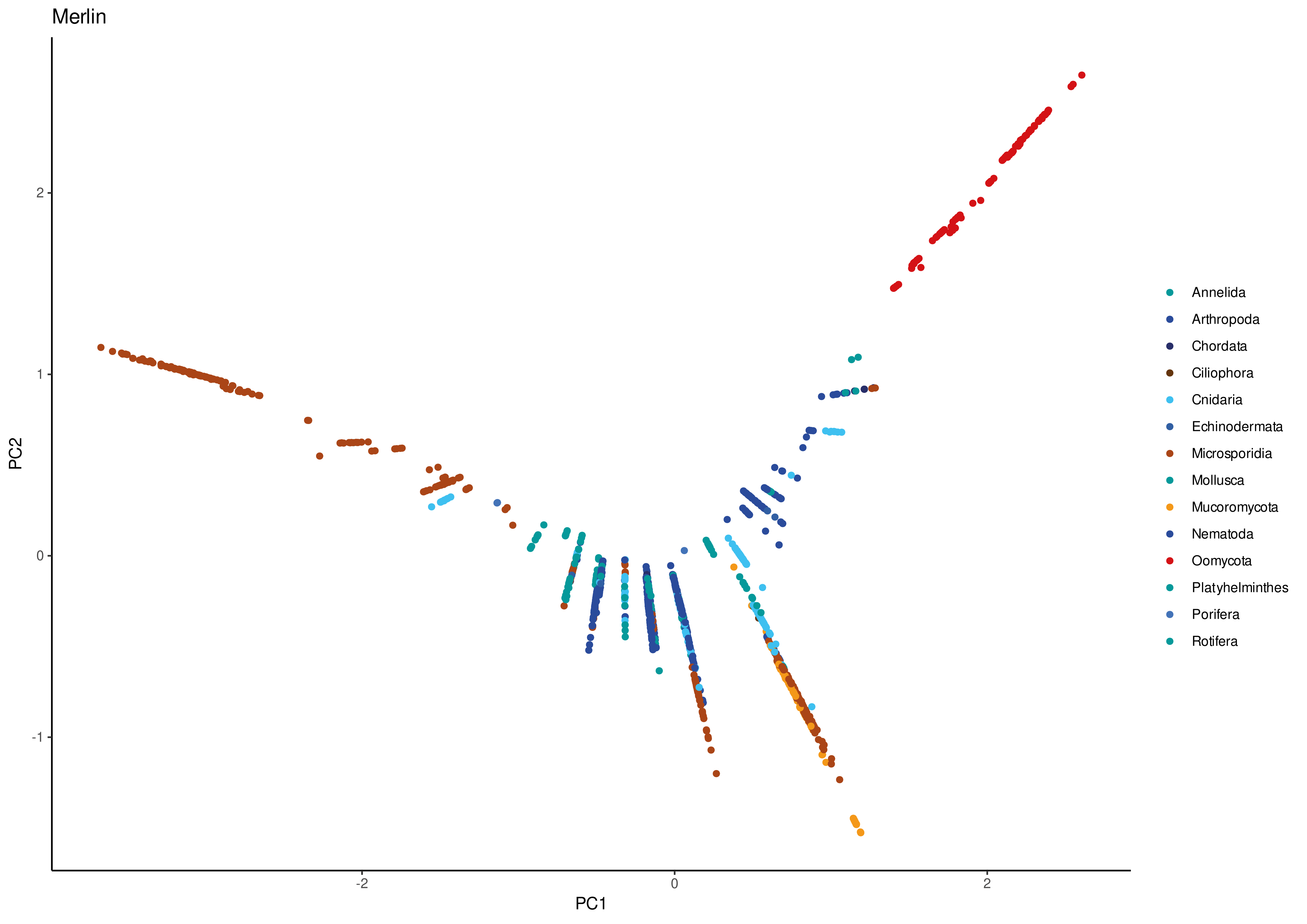
Merlin
PC2
PC1
2
1
0
-1
-2
0
2
Annelida
Arthropoda
Chordata
Ciliophora
Cnidaria
Echinodermata
Microsporidia
Mollusca
Mucoromycota
Nematoda
Oomycota
Platyhelminthes
Porifera
Rotifera

Mutator

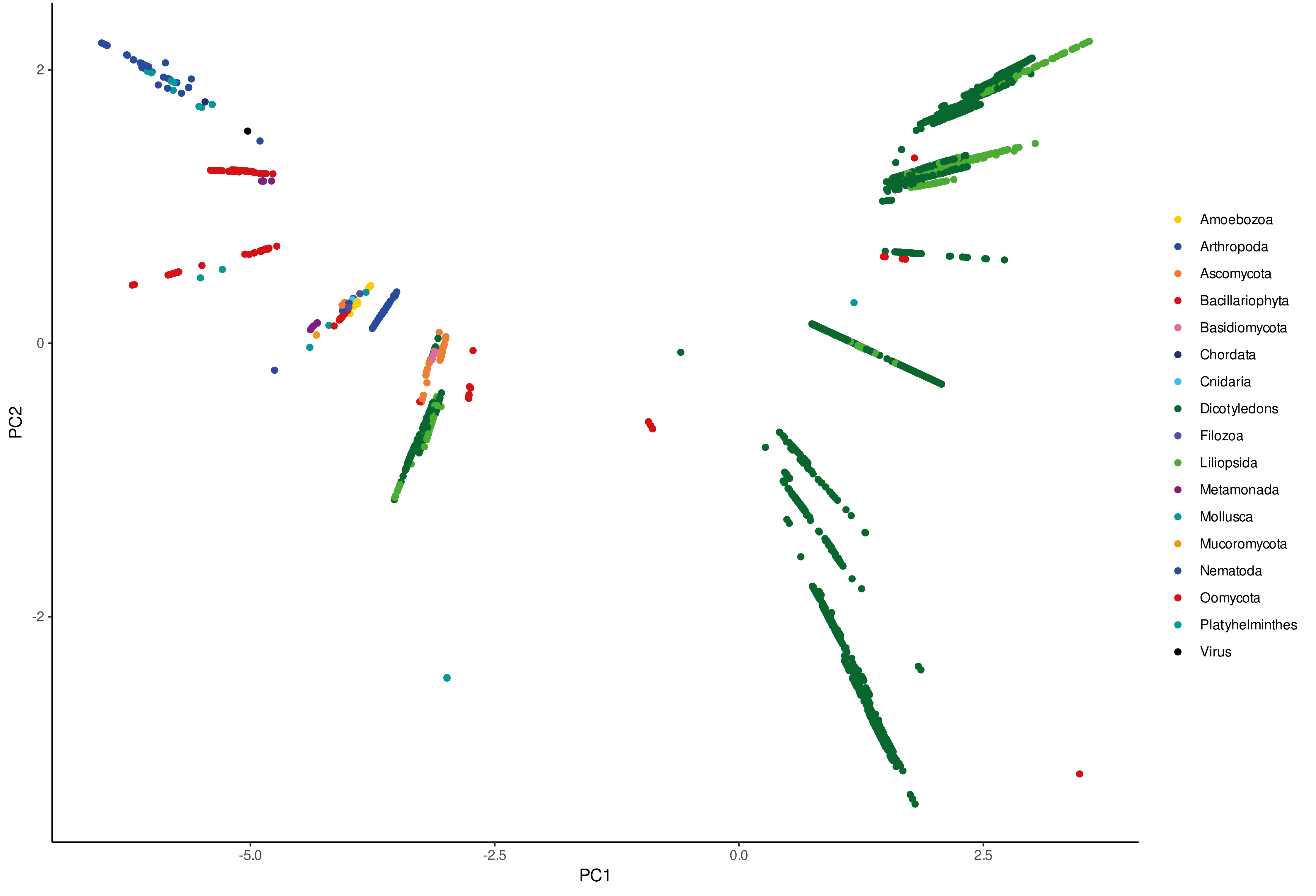

PC2
PC1
2
0
-2
-5.0
-2.5
0.0
2.5
Amoebozoa
Arthropoda
Ascomycota
Bacillariophyta
Basidiomycota
Chordata
Cnidaria
Dicotyledons
Filozoa
Liliopsida
Metamonada
Mollusca
Mucoromycota
Nematoda
Oomycota
Platyhelminthes
Virus

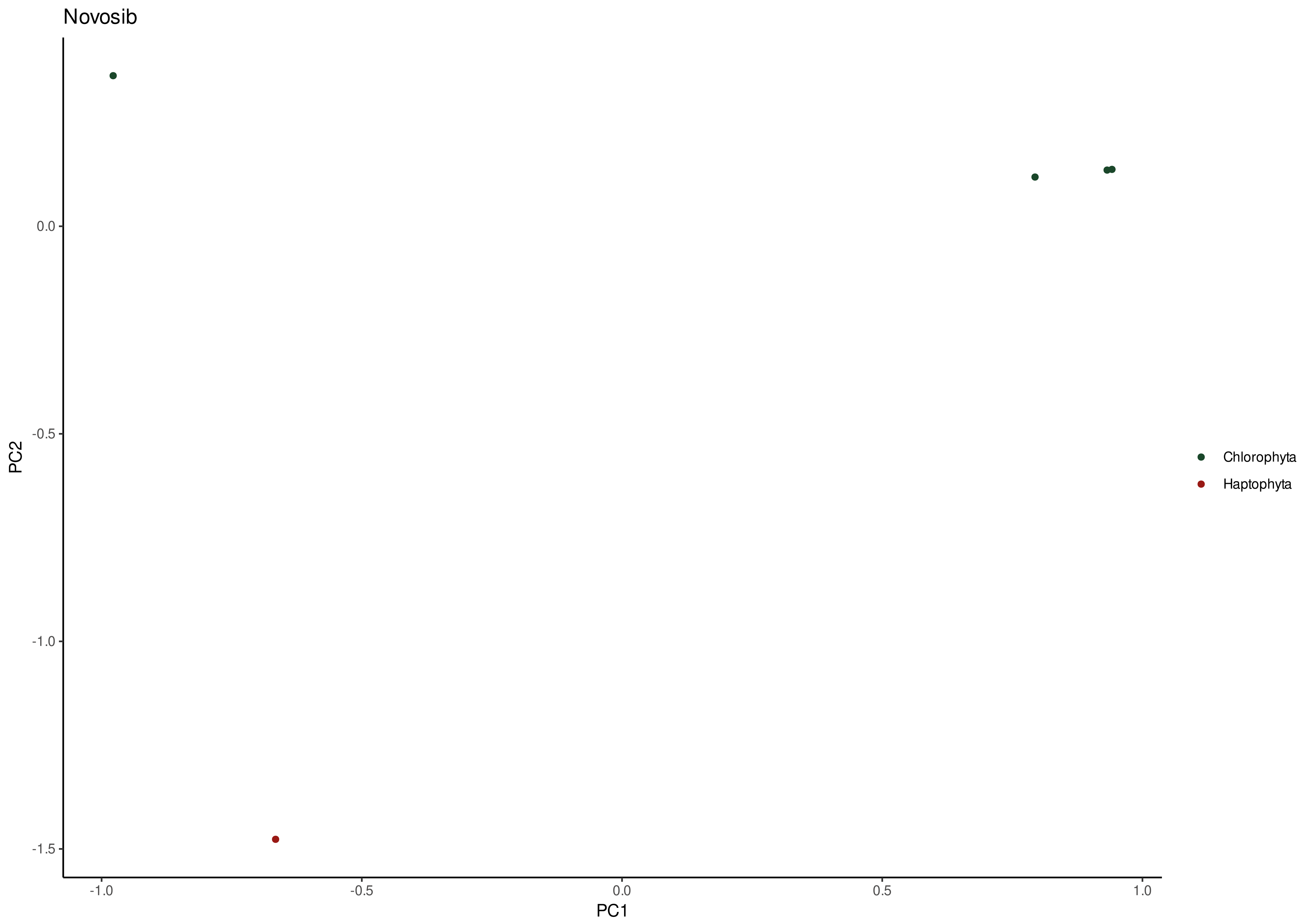
Novosib
PC2
0.0
-0.5
-1.0
-1.5
-1.0
-0.5
0.0
0.5
1.0
PC1
Chlorophyta
Haptophyta

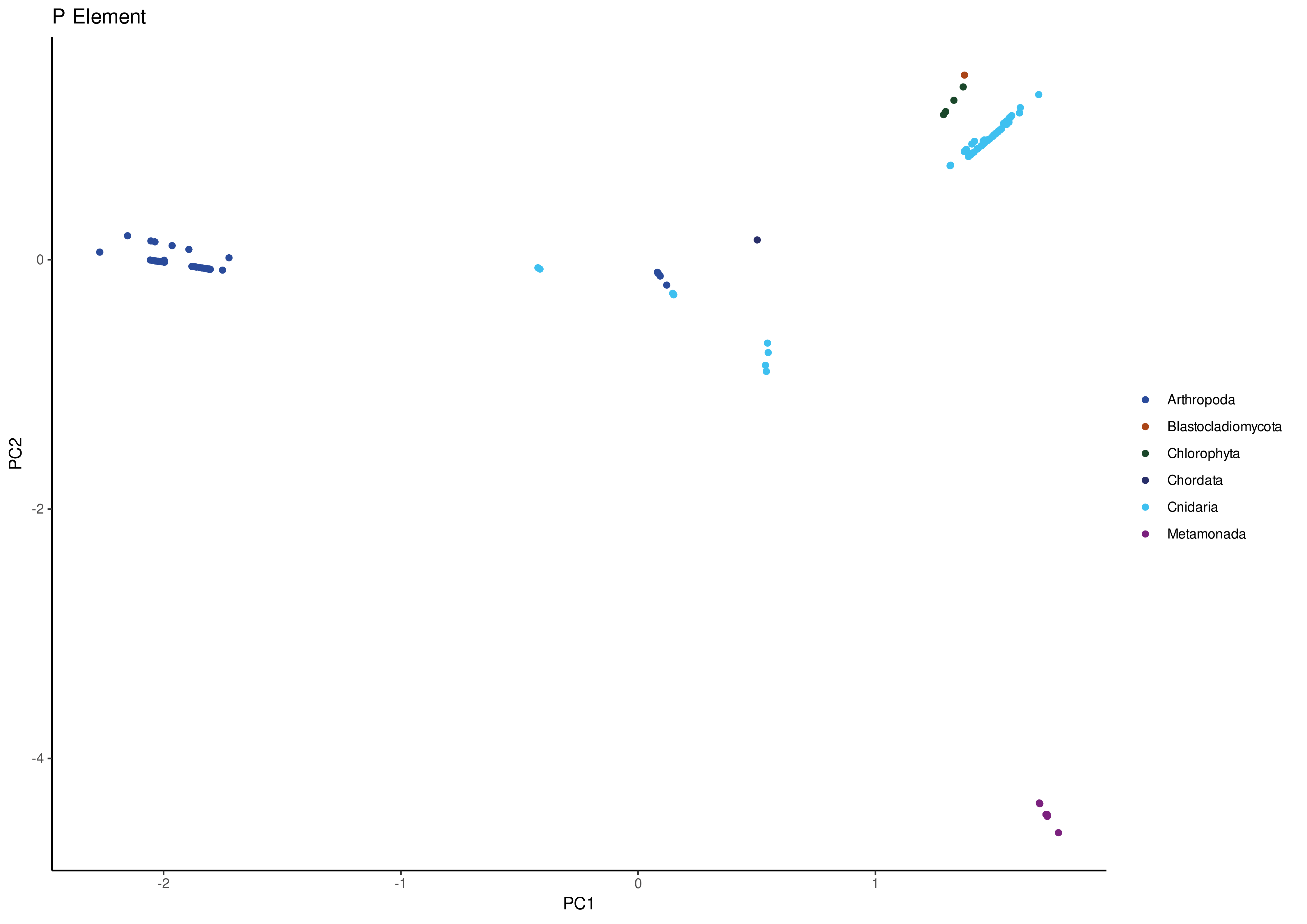
P Element
PC2
0
-2
-4
-2
-1
0
1
PC1
Arthropoda
Blastocladiomycota
Chlorophyta
Chordata
Cnidaria
Metamonada

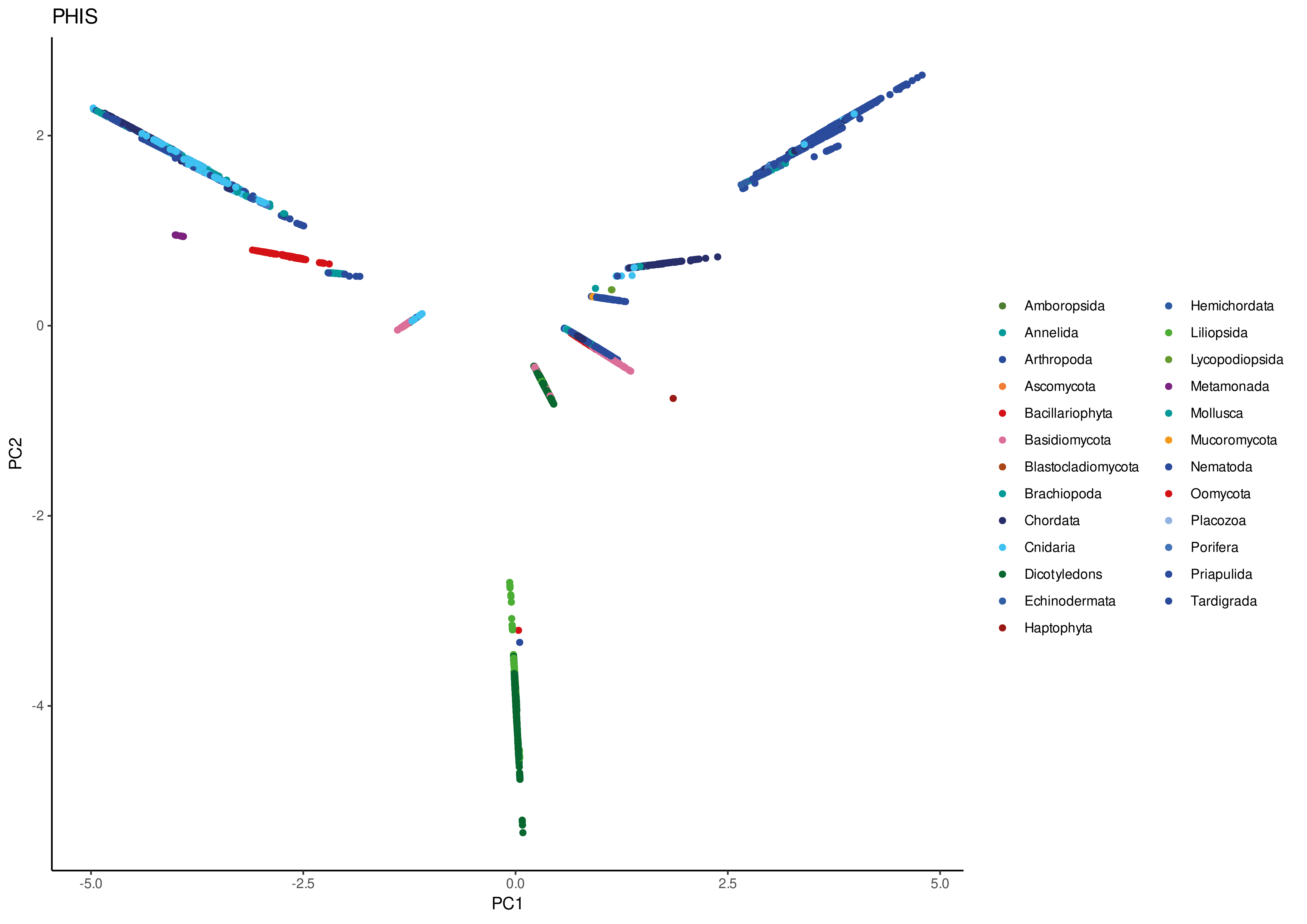
PHIS
PC1
PC2
-5.0
-2.5
0.0
2.5
5.0
2
0
-2
-4
Amboropsida
Annelida
Arthropoda
Ascomycota
Bacillariophyta
Basidiomycota
Blastocladiomycota
Brachiopoda
Chordata
Cnidaria
Dicotyledons
Echinodermata
Haptophyta
Hemichordata
Liliopsida
Lycopodiopsida
Metamonada
Mollusca
Mucoromycota
Nematoda
Oomycota
Placozoa
Porifera
Priapulida
Tardigrada

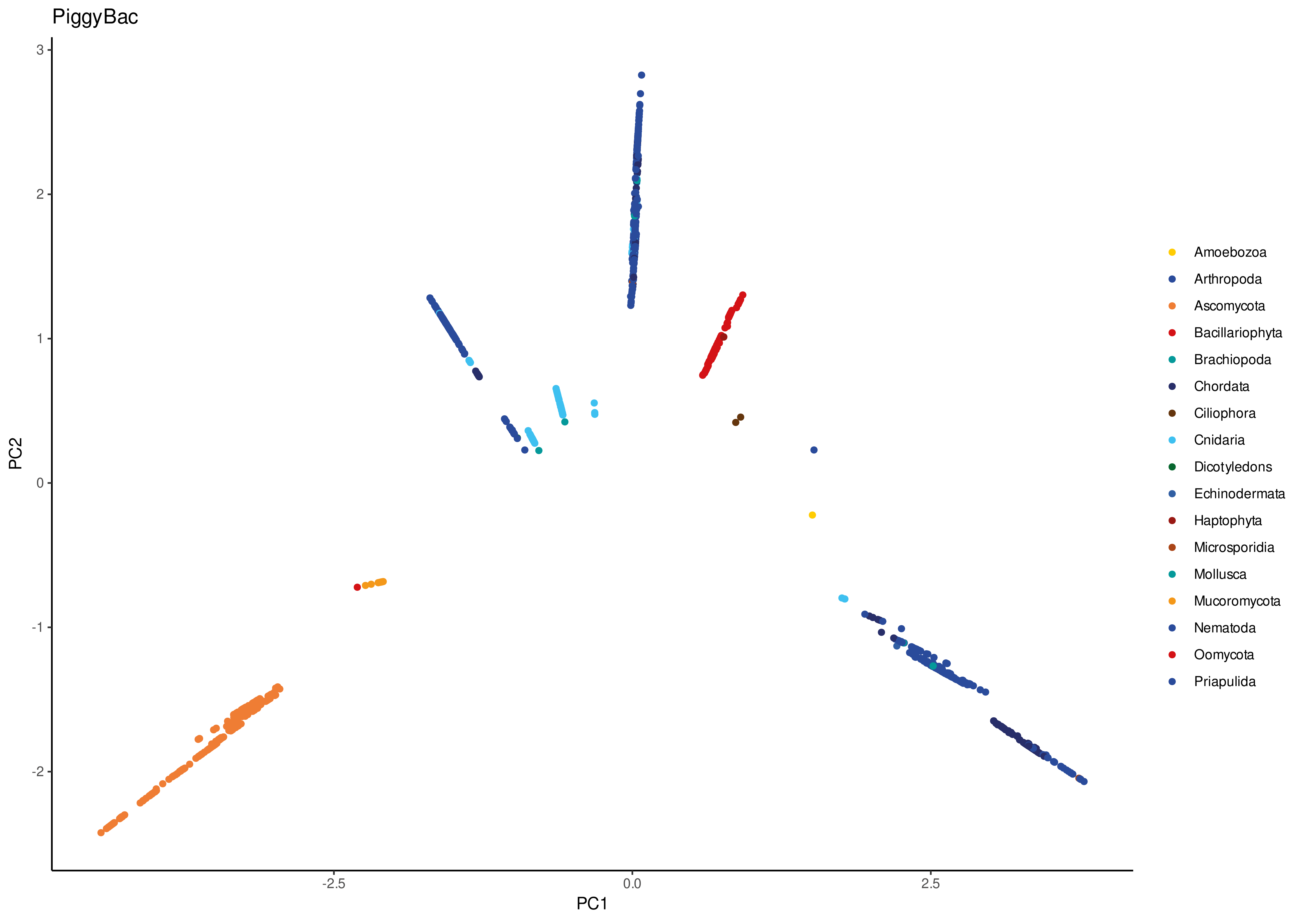
PiggyBac
PC1
PC2
-2.5
0.0
2.5
3
2
1
0
-1
-2
Amoebozoa
Arthropoda
Ascomycota
Bacillariophyta
Brachiopoda
Chordata
Ciliophora
Cnidaria
Dicotyledons
Echinodermata
Haptophyta
Microsporidia
Mollusca
Mucoromycota
Nematoda
Oomycota
Priapulida

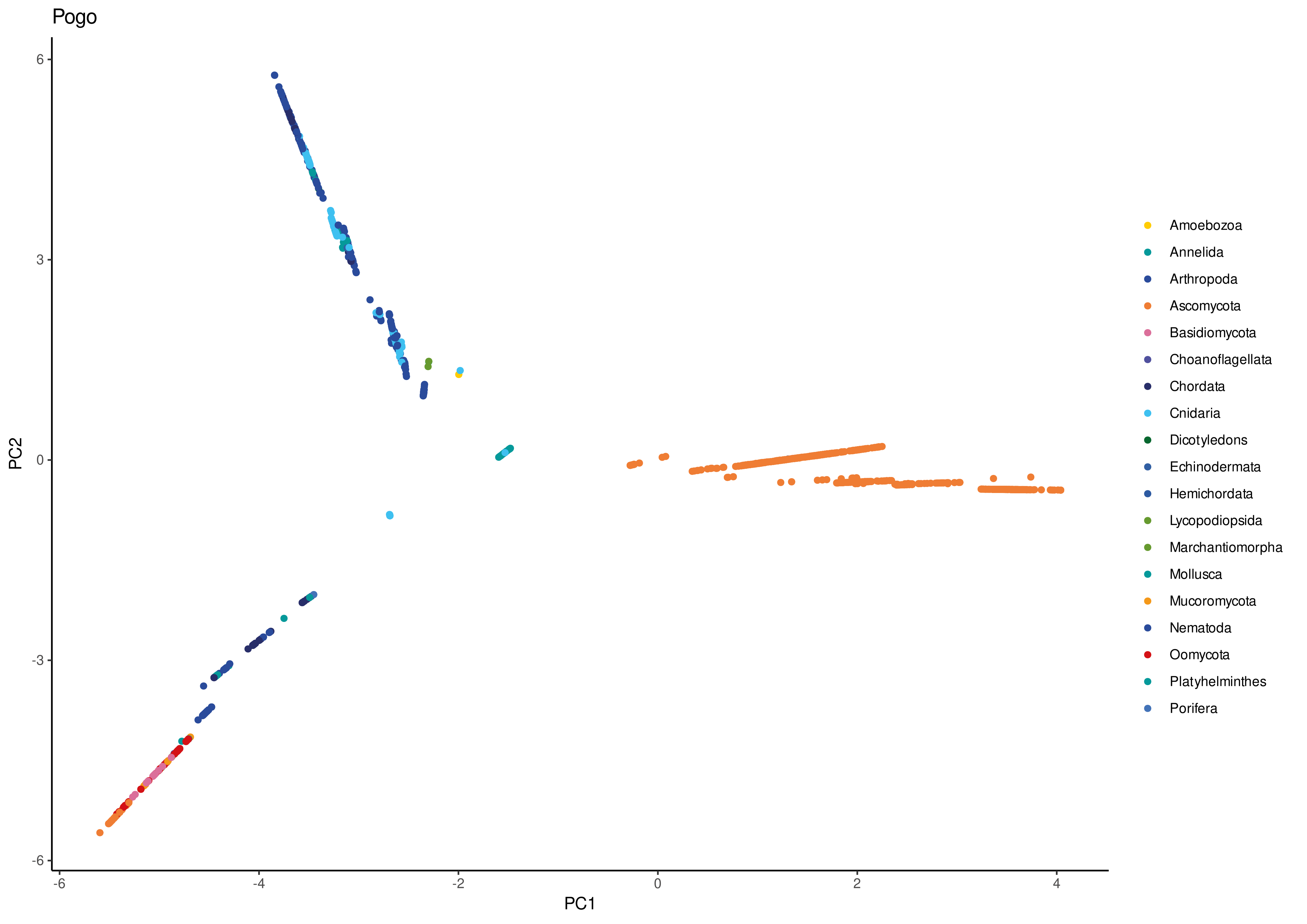
Pogo
PC1
PC2
-6
-4
-2
0
2
4
6
3
0
-3
-6
Amoebozoa
Annelida
Arthropoda
Ascomycota
Basidiomycota
Choanoflagellata
Chordata
Cnidaria
Dicotyledons
Echinodermata
Hemichordata
Lycopodiopsida
Marchantiomorpha
Mollusca
Mucoromycota
Nematoda
Oomycota
Platyhelminthes
Porifera

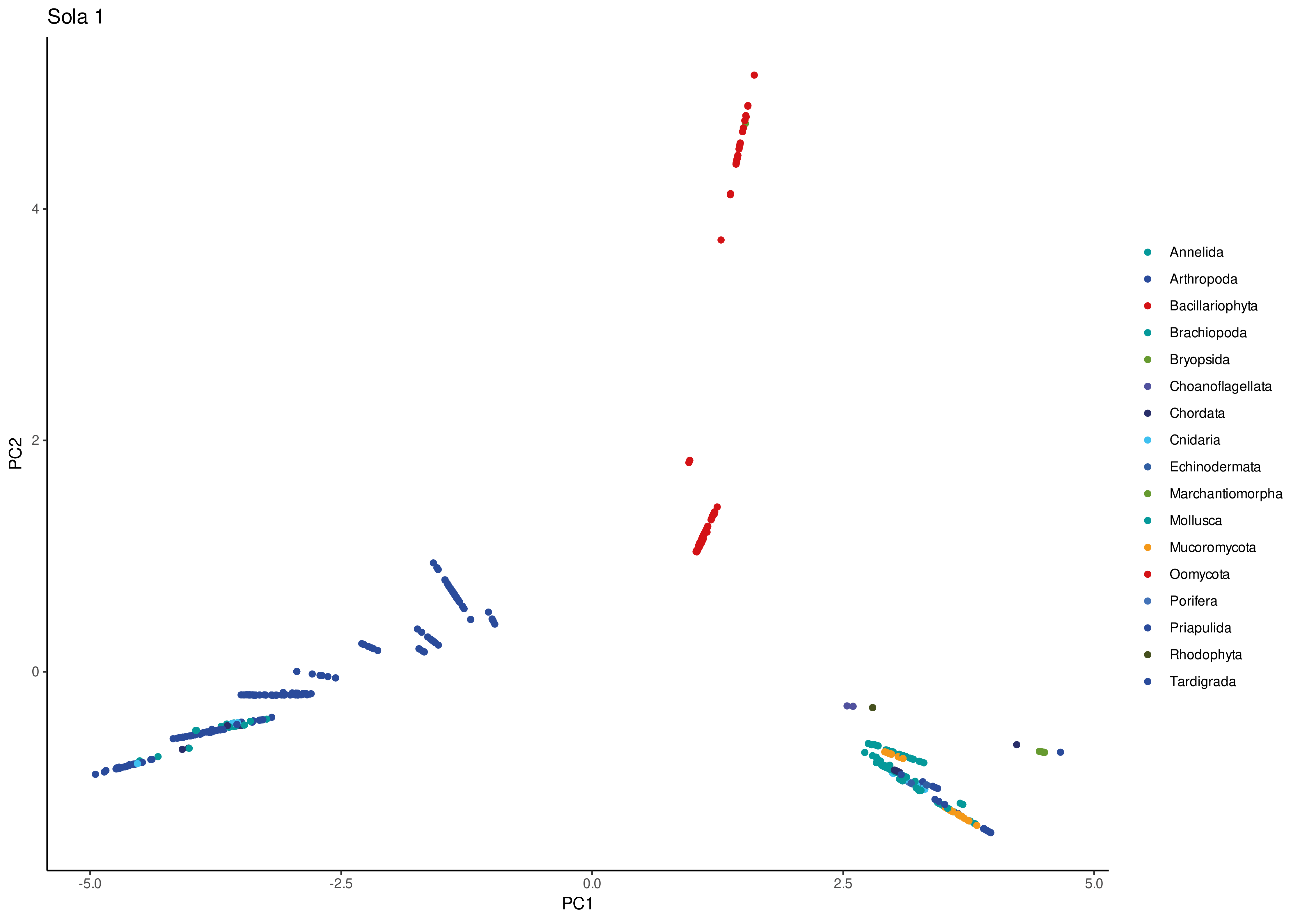
Sola 1
PC2
PC1
4
2
0
-5.0
-2.5
0.0
2.5
5.0
Annelida
Arthropoda
Bacillariophyta
Brachiopoda
Bryopsida
Choanoflagellata
Chordata
Cnidaria
Echinodermata
Marchantiomorpha
Mollusca
Mucoromycota
Oomycota
Porifera
Priapulida
Rhodophyta
Tardigrada

Sola 2

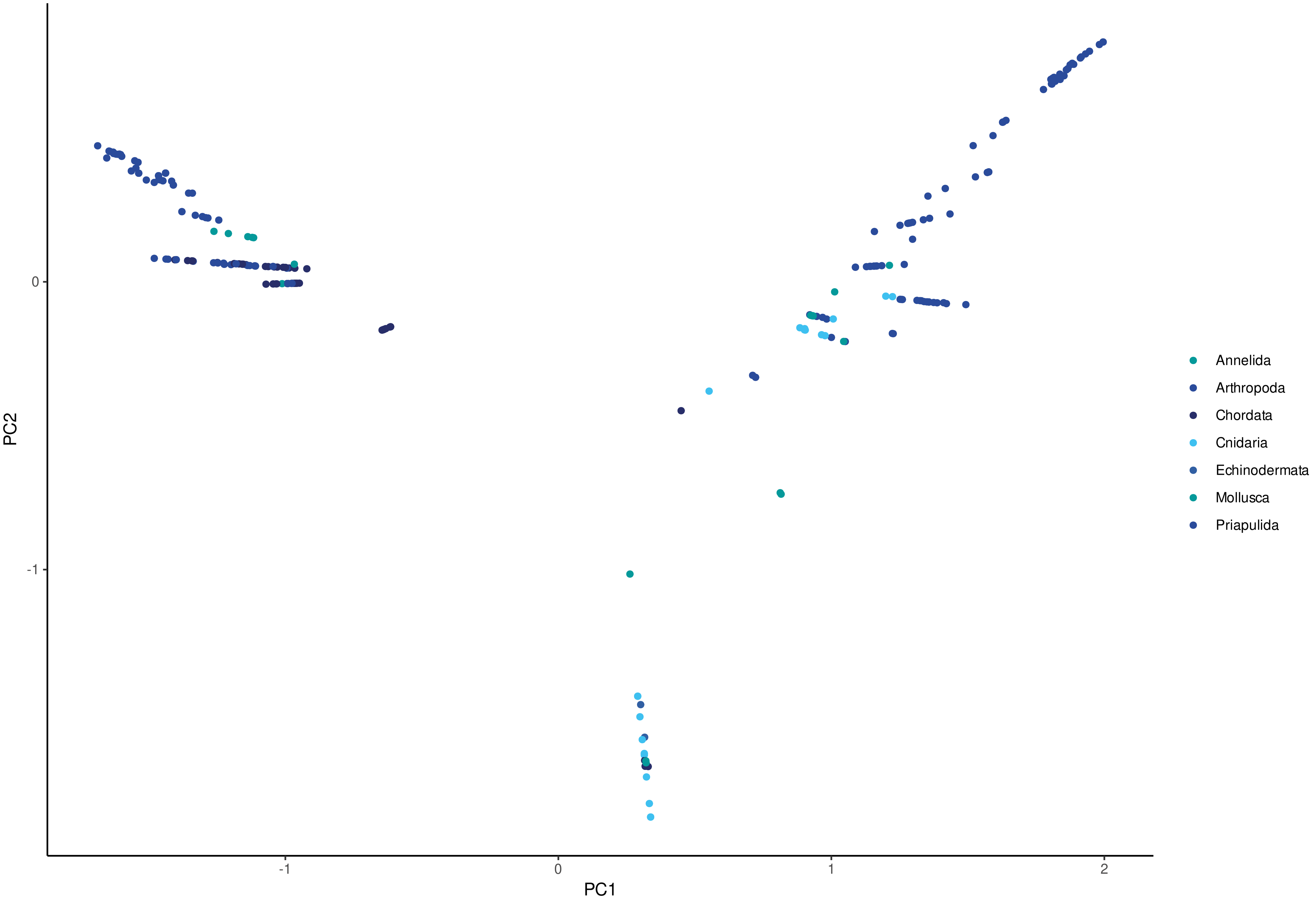

PC2
0
-1
PC1
-1
0
1
2
Annelida
Arthropoda
Chordata
Cnidaria
Echinodermata
Mollusca
Priapulida

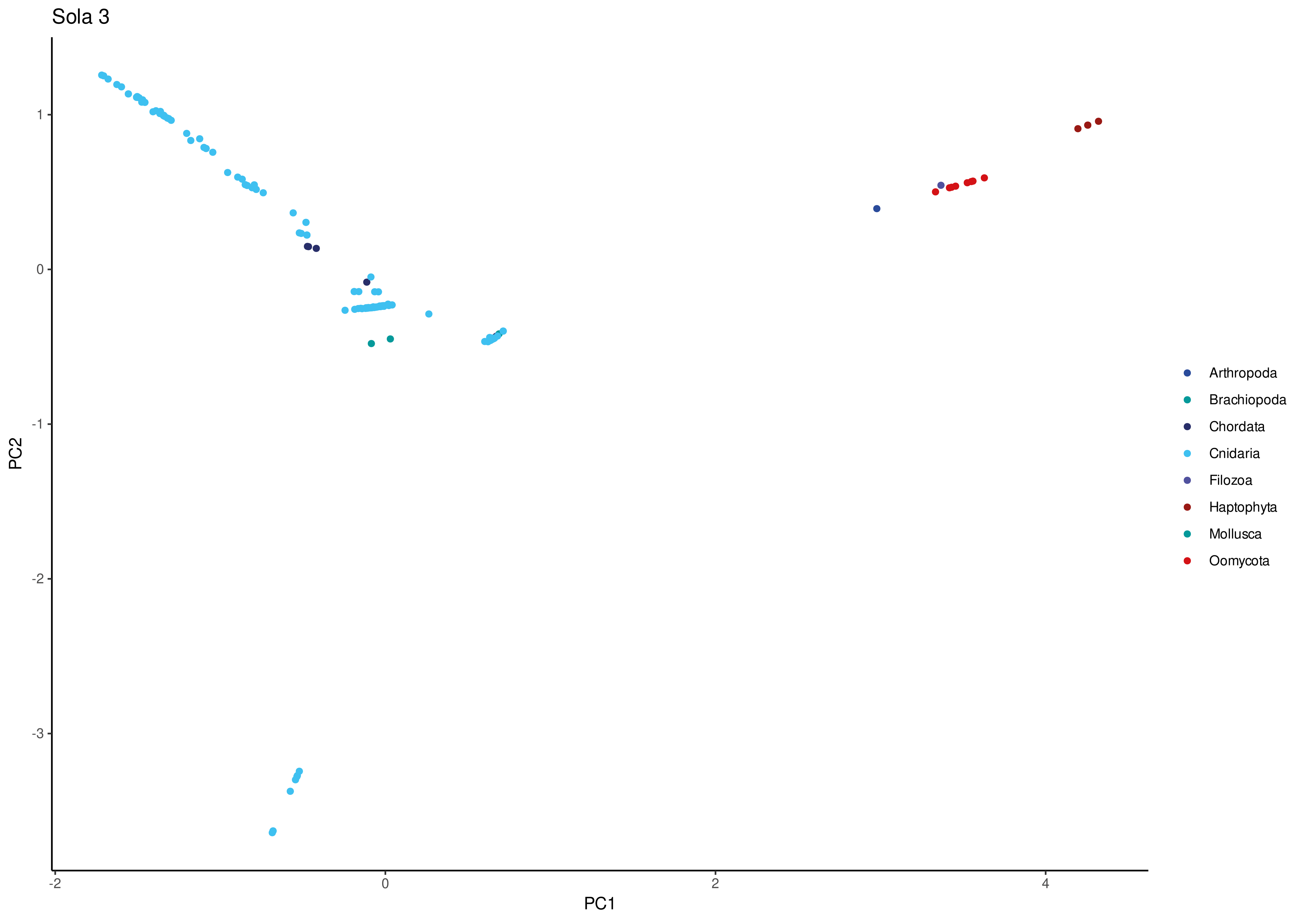

Sola 3
PC2
1
0
-1
-2
-3
PC1
-2
0
2
4
Arthropoda
Brachiopoda
Chordata
Cnidaria
Filozoa
Haptophyta
Mollusca
Oomycota

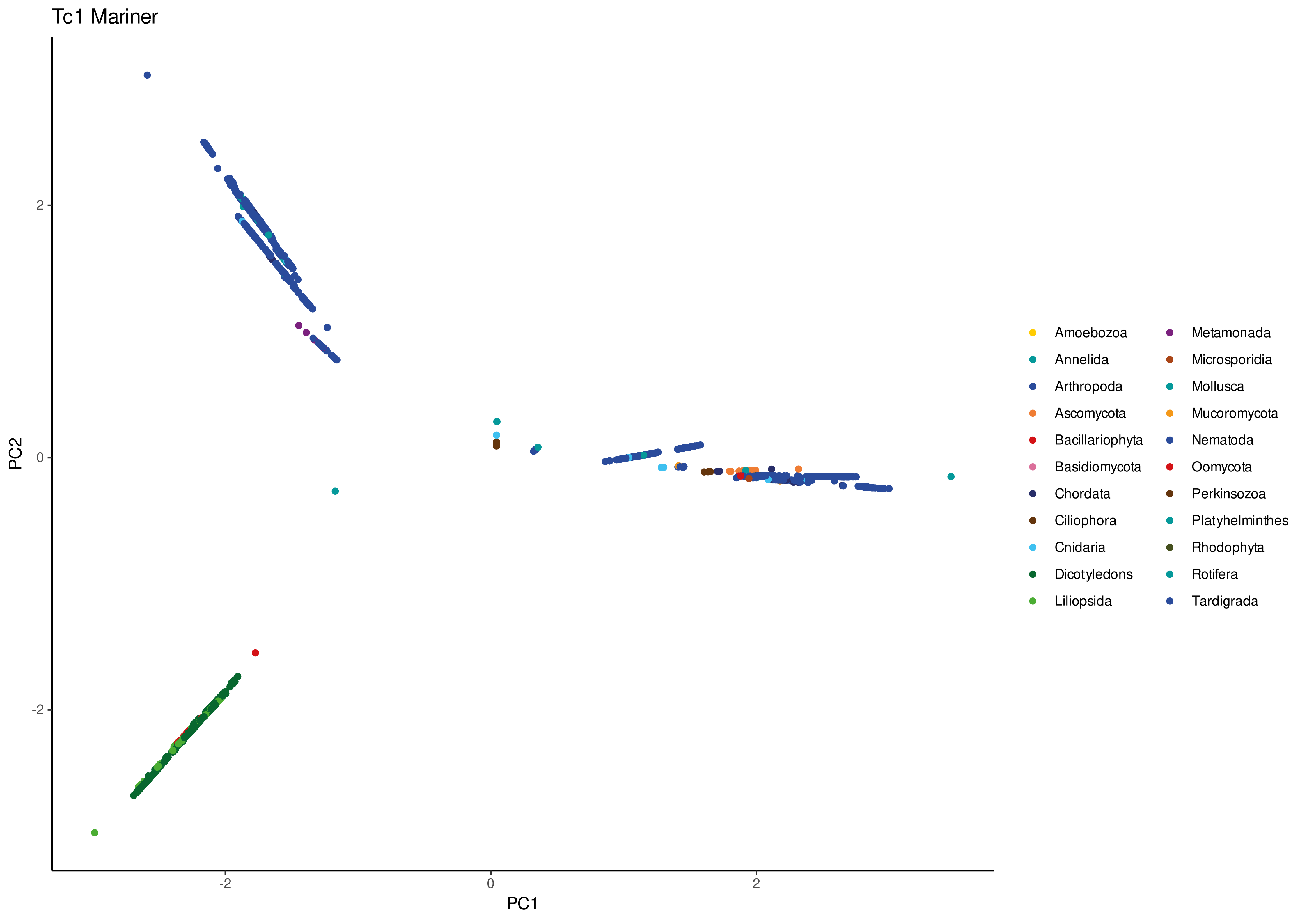

Tc1 Mariner
PC2
2
0
-2
PC1
-2
0
2
Amoebozoa
Annelida
Arthropoda
Ascomycota
Bacillariophyta
Basidiomycota
Chordata
Ciliophora
Cnidaria
Dicotyledons
Liliopsida
Metamonada
Microsporidia
Mollusca
Mucoromycota
Nematoda
Oomycota
Perkinsozoa
Platyhelminthes
Rhodophyta
Rotifera
Tardigrada

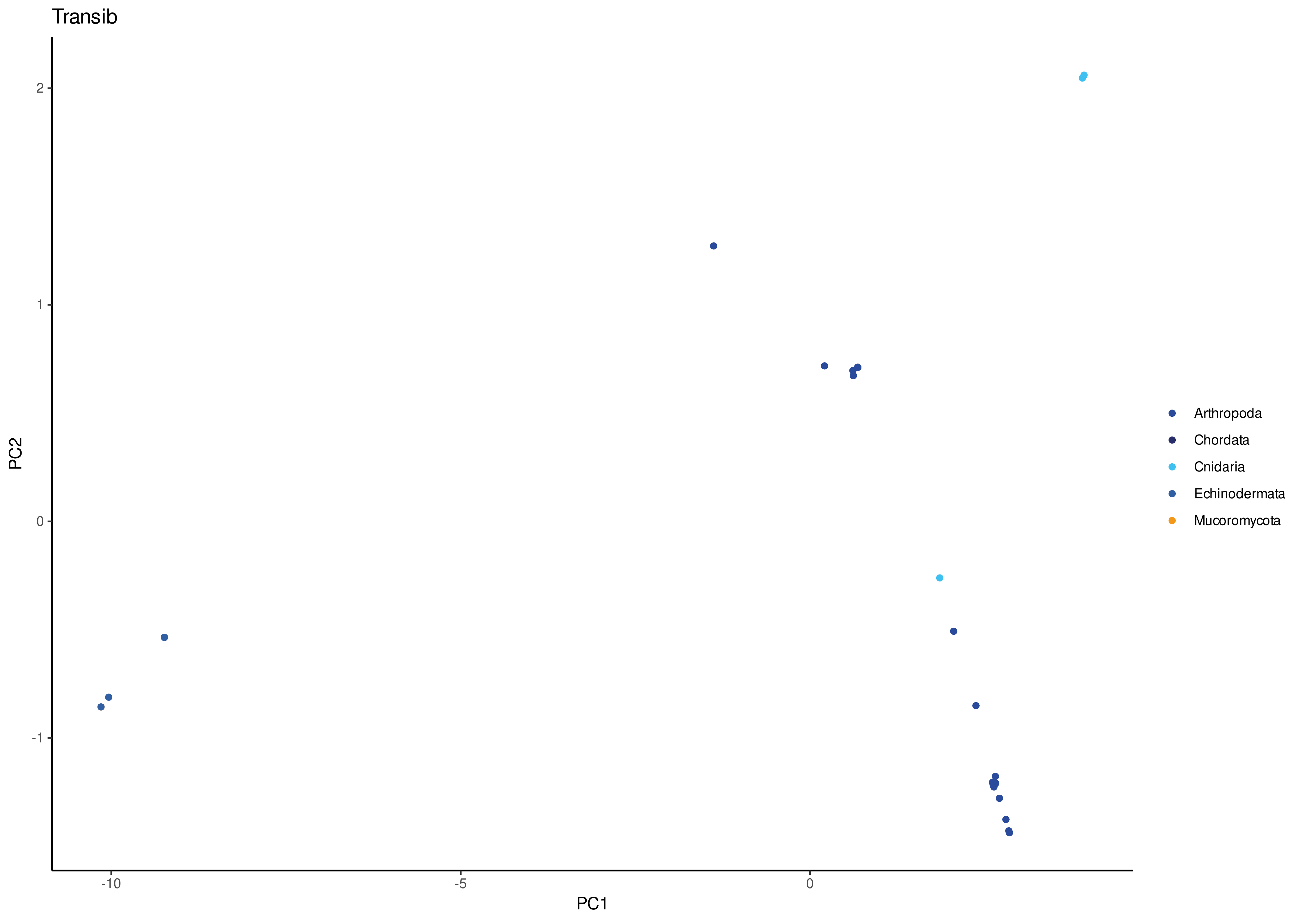
Transib
PC2
PC1
2
1
0
-1
-10
-5
0
Arthropoda
Chordata
Cnidaria
Echinodermata
Mucoromycota

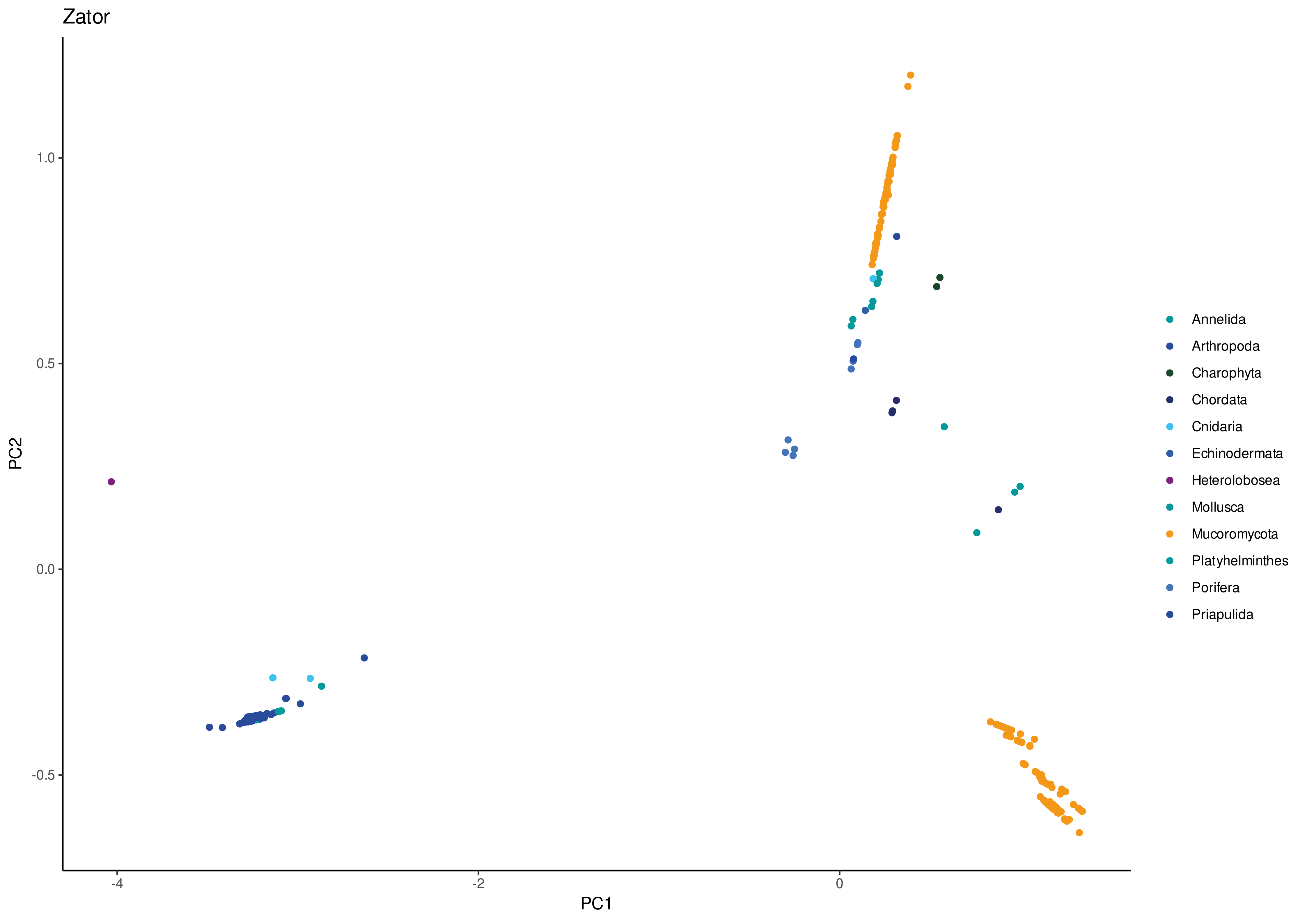
Zator
PC1
PC2
-4
-2
0
1.0
0.5
0.0
-0.5
Annelida
Arthropoda
Charophyta
Chordata
Cnidaria
Echinodermata
Heterolobosea
Mollusca
Mucoromycota
Platyhelminthes
Porifera
Priapulida

Table S1. Number of transposases recovered for each DDE TE superfamily

| DDE TE superfamily | No. transposases |
|---|---|
| Academ | 234 |
| CMC | 1085 |
| Ginger | 1079 |
| GingerRoot | 1186 |
| hAT | 4714 |
| Kolobok | 809 |
| Merlin | 1509 |
| Mutator | 4536 |
| Novosib | 10 |
| P | 193 |
| PHIS | 5738 |
| PiggyBac | 1351 |
| Pogo | 1775 |
| Sola1 | 760 |
| Sola2 | 272 |
| Sola3 | 167 |
| Tc1mar | 3052 |
| Transib | 35 |
| Zator | 368 |
| Sum | 28873 |

Table S2. Mean tree height for each DDE TE superfamily

| DDE TE superfamily | Tree height |
|---|---|
| Academ | 5.301 |
| CMC | 4.385 |
| Ginger | 3.568 |
| GingerRoot | 6.176 |
| hAT | 4.941 |
| Kolobok | 4.837 |
| Merlin | 3.887 |
| Mutator | 9.199 |
| Novosib | 2.343 |
| P | 3.696 |
| PHIS | 4.052 |
| PiggyBac | 9.631 |
| Pogo | 7.118 |
| Sola1 | 7.766 |
| Sola2 | 3.535 |
| Sola3 | 7.169 |
| Tc1mar | 3.469 |
| Transib | 7.355 |
| Zator | 2.710 |
| Mean | 5.323 |

Table S3. Species diversity of major host groups and corresponding numbers of transposases recovered

| Host group | Estimated species diversity | Transposases recovered |
|---|---|---|
| Excavata | 600 | 311 |
| Amoebozoa | 13300 | 33 |
| HaptophytaSAR | 29907 | 1160 |
| Holomycota | 97101 | 3330 |
| Archaeplastida | 416400 | 9505 |
| Holozoa | 1274280 | 14531 |
| | 1831588 | |
| | 3663176 | |

Based on the following taxonomic breakdown:

| Host subgroup | Estimated number of species | References |
|---|---|---|
| Amoebozoa | 13,300 to 22,600 | Adl et al. 2007 |
| Ciliophora | > 8,000 | Shazib et al. 2019 |
| Perkinsozoa | At least 11 but poorly known | Jeon et al. 2019 |
| Heterolobosea | About 150 | Panek et al. 2016 |
| Parabasalia | About 450 | Cepicka et al. 2016 |
| Ascomycota | > 64,000 | Wijayawardene et al. 2017 |
| Basidiomycota | > 31,000 | Taylor et al. 2015 |
| Blastocladiomycota | About 183 | Wijayawardene et al. 2018 |
| Microsporidia | About 1400 | Han & Weiss 2017 |
| Mucoromycota | About 312 | Wijayawardene et al. 2018 |
| Zoopagomycota | About 206 | Wijayawardene et al. 2018 |
| Annelida | > 21,000 | Weigert et al. 2016 |
| Arthropoda | About 1,020,000 described to 81.6 millions | Giribet & Edgecombe 2019 and Larsen et al. 2017 |
| Brachiopoda | About 350 | Herper et al. 2017 |

| | | |
|---|---|---|
| Chordata | 70,000 to 80,000 | Larsen et al. 2017 |
| Cnidaria | > 14,350 | Okamura & Gruhl 2020 |
| Echinodermata | About 7000 | Russell 2013 |
| Hemichordata | > 130 | Tassia et al. 2016 |
| Mollusca | About 70,000 to 200,000 estimated | Rosenberg 2014 |
| Nematoda | 30,000 described to 40.8 millions | Smythe et al. 2019 and Larsen et al. 2017 |
| Placozoa | About 100 | Schierwater & DeSalle 2018 |
| Platyhelminthes | > 20,000 | Adell et al. 2015 |
| Porifera | > 8,000 | Manconi & Pronzato |
| Priapulida | 20 described | Schmidt-Rhaesa et al. 2017 |
| Rotifera | About 2,000 | Wallace et al. 2015 |
| Tardigrada | > 1,200 | Nelson et al. 2015 |
| Chlorophyta | > 5,300 | Rioux & Turgeon 2015 |
| Rhodophyta | > 7,100 | Takaishi et al. 2016 |
| Streptophyta | About 404,000 | Lughadha et al. 2016 |
| Choanoflagellata | > 125 | King et al. 2008 |
| Filozoa | 5 described | Southworth et al. 2018 |
| Haptista | > 96 (genus level), > 500 Haptophyta | Adl et al. 2019, Guiry 2012 |
| Bacillariophyta | About 21,000 | Guiry 2012 |
| Oomycota | About 800 | Judelson 2017 |

Table S4. Host range at different taxonomic levels for each DDE TE superfamily

| DDE TE superfamily | Families | Orders | Classes | Phyla | Kingdoms | Total |
|---|---|---|---|---|---|---|
| Academ | 34 | 30 | 18 | 9 | 2 | 234 |
| CMC | 108 | 68 | 27 | 15 | 4 | 1085 |
| Ginger | 91 | 61 | 29 | 17 | 4 | 1079 |
| GingerRoot | 83 | 52 | 24 | 13 | 3 | 1186 |
| hATs | 274 | 142 | 34 | 17 | 5 | 4714 |
| Kolobok | 62 | 44 | 24 | 15 | 4 | 809 |
| Merlin | 90 | 50 | 30 | 15 | 5 | 1509 |
| Mutator | 96 | 60 | 26 | 18 | 8 | 4536 |
| Novosib | 3 | 2 | 2 | 2 | 2 | 10 |
| P | 18 | 13 | 10 | 9 | 5 | 193 |
| PHIS | 222 | 118 | 47 | 23 | 6 | 5738 |
| Piggybac | 118 | 68 | 32 | 18 | 7 | 1351 |
| Pogo | 115 | 73 | 38 | 17 | 6 | 1775 |
| Sola1 | 58 | 43 | 26 | 16 | 5 | 760 |
| Sola2 | 57 | 33 | 15 | 7 | 1 | 272 |
| Sola3 | 15 | 12 | 11 | 8 | 4 | 167 |
| Tc1mar | 151 | 83 | 37 | 21 | 7 | 3052 |
| Transib | 13 | 11 | 7 | 5 | 2 | 35 |
| Zator | 29 | 22 | 16 | 12 | 4 | 368 |
| | | | | | Total | 28873 |

Table S5. Number of estimated horizontal transfer events for each DDE TE superfamily

| | Family | Order | Class | Phylum | Kingdom |
|---|---|---|---|---|---|
| Academ | 80 | 73 | 50 | 39 | 2 |
| CMC | 202 | 162 | 55 | 47 | 6 |
| Ginger | 156 | 132 | 65 | 49 | 16 |
| GingerRoot | 403 | 346 | 100 | 89 | 8 |
| hAT | 841 | 615 | 192 | 112 | 18 |
| Kolobok | 107 | 83 | 38 | 28 | 3 |
| Merlin | 195 | 146 | 113 | 81 | 22 |
| Mutator | 432 | 330 | 75 | 41 | 27 |
| Novosib | 3 | 1 | 1 | 1 | 1 |
| P | 30 | 22 | 12 | 10 | 4 |
| PHIS | 1173 | 906 | 334 | 234 | 28 |
| Piggybac | 361 | 216 | 87 | 57 | 10 |
| Pogo | 294 | 219 | 108 | 72 | 16 |
| Sola1 | 175 | 127 | 50 | 42 | 8 |
| Sola2 | 99 | 76 | 29 | 24 | 0 |
| Sola3 | 36 | 31 | 13 | 11 | 3 |
| Tc1mar | 484 | 335 | 137 | 83 | 19 |
| Transib | 13 | 11 | 9 | 6 | 1 |
| Zator | 43 | 38 | 23 | 17 | 4 |

Table S6. Verification checks of transposases labelled as originating from prokaryotes

| TE superfamily | GenInfo identifier | GenBank accession | Host class | Host order | Host family | Host species | Habitat | Flanking region check | BLASTn with nucleotidic sequence | BLASTp | TE from bacteria? |
|---|---|---|---|---|---|---|---|---|---|---|---|
| Academ | 154390699 | RUA06787 | Gammaproteobacteria | - | - | - | Various environmental, phytopathogens, animal pathogens | No TIRs, full sequence 3103 bp (from hydrothermal vent metagenome) | No match | A few sequences from sea anemones but low query cover and %ID values | No - Probably contamination |
| CMC | 159591543 | WP133372327 | Bacilli | Bacillales | Bacillaceae | Bacillus_aciditolerans | Rice paddy soil | Record removed, not annotated anymore | - | - | - |
| CMC | 115017517 | OON91157 | Clostridia | Clostridiales | Lachnospiraceae | Epulopiscium_sp._AS2M-Bin002 | Symbiotic relationship with sturgeonfish (gut) | No TIRs, full sequence 11497 bp from the gut metagenome of *Acanthurus sohal* (red sea sturgeonfish) | No match with >50% of query cover | Quite a lot of sequences from fish species (especially groupers) | No - Probably contamination |
| Ginger | 140139123 | PYY19639 | - | - | - | - | Various environmental | No TIRs (only 45 bp before protein), full sequence 2506 bp from a soil sample (soil metagenome) | No match | One sequence 91% query cover 53% id from a fungus | No - Probably contamination |
| Ginger | 140139253 | PYY20810 | - | - | - | - | Various environmental | No TIRs, full sequence 4357 bp from a soil sample (soil metagenome) | 2 matches (*Gossypium* and a cestode) on 5' flank 56% query cover >90% id | Two sequences 98% query cover 51% id from fungi | No - Probably contamination |
| Ginger | 140139077 | PYY19226 | - | - | - | - | Various environmental | No TIRs, partial sequence 1631 bp from a soil sample (soil metagenome) | No match with >50% of query cover | Two sequences 99% query cover 50% and 59% id from fungi | No - Probably contamination |
| Ginger | 140139151 | PYY19889 | - | - | - | - | Various environmental | No TIRs, full sequence 3608 bp from a soil sample (soil metagenome) | No match | Several sequences >90% query cover >50% id from fungi, plants and mussels | No - Probably contamination |
| Ginger | 156801327 | RYE21344 | Sphingobacteriia | Sphingobacteriales | Sphingobacteriaceae | | Various environmental | No TIRs, partial sequence 2402 bp from phyllosphere metagenome on *Populus alba* | No match | Two sequences 99% query cover 50% from *Citrus clementina* (not bacteria) | No - Probably contamination |
| Ginger | 112543532 | OLD63605 | - | - | - | - | Various environmental | No TIRs, full sequence 10908 bp from a soil sample (soil metagenome) | Matches 3' flank with *V. vinifera* (not bacteria) 76,3% query cover and 97,73% id | No result >50% id | No - Probably contamination |
| Ginger | 154126260 | RTK47865 | Bacilli | Lactobacillales | Enterococcaceae | Enterococcus_faecalis | Symbiotic relationship in mammals (gut) | No TIRs, full sequence 281 bp from human bile | Three matches with *Arachis hypogaea* | Lots of matches from nuts but not suprising giving the short sequence | No - Probably contamination |
| Ginger | 117867333 | WP083183379 | Betaproteobacteria | Burkholderiales | Burkholderiaceae | Candidatus_Glomeribacter_gigasporarum | Obligate endosymbiote of fungus *Gigaspora margarita* | Record removed, not annotated anymore | - | - | - |
| GingerRoot | 118263508 | ORU94823 | Gammaproteobacteria | Thiotrichales | Piscirickettsiaceae | Cycloclasticus_sp | Symbiont_of_Poecilosclerida_sp (sponge), marine sediments | No TIRs, full sequence 4212 bp from *Poecilosclerida* sp. (symbiont) | 1 match 3' flank with *P. caudatus* (not a bacteria) 69,4% query cover and 100% id | A few matches >50% query cover and %ID with Porifera species | No - Probably contamination |
| GingerRoot | 118263526 | ORU94940 | Gammaproteobacteria | Thiotrichales | Piscirickettsiaceae | Cycloclasticus_sp | Symbiont_of_Poecilosclerida_sp (sponge), marine sediments | No TIRs, full sequence 3382 bp from *Poecilosclerida* sp. (symbiont) | No match | A few matches >50% query cover and %ID with Porifera species | No - Probably contamination |
| GingerRoot | 154390431 | RUA04468 | Gammaproteobacteria | Gammaproteobacteria | Gammaproteobacteria | - | Various environmental, phytopathogens, animal pathogens | No TIRs, full sequence 2604 bp from hydrothermal vent (Mariana) | A few matches 3' flank with *N. vectensis* (not a bacteria) 82% query cover and 86,67% id | A few matches >50% query cover and %ID with Cnidaria species | No - Probably contamination |
| GingerRoot | 154390447 | RUA04598 | Gammaproteobacteria | Gammaproteobacteria | Gammaproteobacteria | - | Various environmental, phytopathogens, animal pathogens | Long TIRs of 129 bp but weirdly start 100 bp in the protein, so it could be a palindromic sequence not linked to TIRs. Full sequence 4728 bp from hydrothermal vent | 1 match 5' flank with a cestode 54,7% query cover and 100% id | A few matches >50% query cover and %ID with Porifera species | No - Probably contamination |
| hAT | 154554031 | RUM29574 | Aquificae | Aquificales | Aquificaceae | Aquifex_sp. | Hot springs | No TIRs, full sequence 3096 bp from hydrothermal vent | No match | A few matches >50% query cover and %ID with an amphibian species | No - Probably contamination |
| hAT | 156801552 | RYE23048 | Sphingobacteriia | Sphingobacteriales | Sphingobacteriaceae | Sphingobacteriaceae | Various environmental, pathogens | No TIRs, full sequence 3076 bp from phyllosphere metagenome on *Populus alba* (silver poplar) | Matches with *Ipomoea sp.* (one TE *Tip100*, not a bacteria) 73% query cover and about 76% id | Matches with different plants on 99% of the query and 59-69% id | No - Probably contamination |
| hAT | 154390475 | RUA04851 | Gammaproteobacteria | - | - | - | Various environmental, phytopathogens, animal pathogens | Potential TIRs of 25 bp, full sequence 4261 bp from hydrothermal vent | No matches >50% query cover and %id | Matches with *S. pistillata* (one transposon BS, not a bacteria) about 70-90% query cover and %id | No - Probably contamination |
| Merlin | 57540448 | ETW92248 | - | - | - | Candidatus_Entotheonella_factor | Symbiont of Theonella swinhoei (sponge) | No TIRs, full sequence 826 bp from *Theonella swinhoei* (symbiont) | No matches | Matches with other Porifera >80% query cover and >50% id | No - Probably contamination |
| Merlin | 118263555 | ORU95108 | Gammaproteobacteria | Thiotrichales | Piscirickettsiaceae | Cycloclasticus_sp. | Marine sediment | No TIRs, full sequence 1901 bp from *Poecilosclerida* sp. (sponge symbiont) | No matches | A few matches with other Porifera >90% query cover and >50% id | No - Probably contamination |
| Merlin | 106853862 | WP069591921 | Gammaproteobacteria | Vibrionales | Vibrionaceae | Vibrio_cholerae | Marine and freshwater animal symbiont | This sequence is only available in amino acid (NR record) and is 94 aa | - | A lot of matches with *Rhizopus delemar* (fungus, not a bacteria) but the sequence is short, 77% query cover and >80% id | No - Probably contamination |
| PHIS | 154553933 | RUM28700 | Aquificae | Aquificales | Aquificaceae | Aquifex_sp. | Hot springs | No TIRs, full sequence 3411 bp from hydrothermal vent | No matches | A few matches from different bacteria and *C. gigas* (Pacific oyster), >95% query covers and >50% id | No - Probably contamination |
| PHIS | 154554190 | RUM31001 | Aquificae | Aquificales | Aquificaceae | Aquifex_sp. | Hot springs | No TIRs, full sequence 3925 bp from hydrothermal vent | No matches | No matches >50% id | No |
| PHIS | 107375700 | SCV66534 | Alphaproteobacteria | Rickettsiales | Anaplasmataceae | Anaplasma_phagocytophilum | Pathogen of animals | No TIRs, full sequence 1756 bp from *A. phagocytophilum* | No matches >50% query cover | Matches from black-legged tick >97% query cover and 50-98% id | No - Probably contamination |
| PHIS | 154555749 | RUM44591 | Deltaproteobacteria | Desulfobacterales | Desulfobulbaceae | Desulfocapsa_sp. | Marine and freshwater habitats | No TIRs, full sequence 3500 bp from hydrothermal vent | No matches | Matches from molluscs and echinoderms >84% query cover and ~55% id | No - Probably contamination |
| PiggyBac | 154554292 | RUM31931 | Aquificae | Aquificales | Aquificaceae | Aquifex_sp. | Hot springs | No TIRs, full sequence 2624 bp from hydrothermal vent | No matches | Only 2 matches with fish 71 and 80% query cover and 50% id | No - Probably contamination |
| PiggyBac | 154390459 | RUA04709 | Gammaproteobacteria | - | - | - | Various environmental, phytopathogens, animal pathogens | No TIRs, partial sequence 3012 bp from hydrothermal vent metagenome | No matches | Matches with cnidarians (often *PiggyBac* TE) >95% query cover and ~55% id | No - Probably contamination |
| Pogo | 68245121 | EAN27256 | Actinobacteria | Micrococcales | Microbacteriaceae | Microbacterium_laevaniformans | Various environmental, phytopathogens, animal pathogens | Record removed, superseeded by complete genome | - | - | - |
| Pogo | 157611348 | RZK36297 | Cytophagia | Cytophagales | Hymenobacteraceae | Hymenobacter_sp. | Tree symbiont (leaves) | No TIRs, full sequence 1972 bp from phyllosphere metagenome (*Populus alba*) | No matches >50% query cover | Matches with ascomycetes 86-100% query cover and 64-83,64% id | No - Probably contamination |
| Pogo | 118140973 | WP084705286 | Alphaproteobacteria | Rhizobiales | Beijerinckiaceae | Beijerinckia_mobilis | Soil | Record removed, not annotated anymore | - | - | - |
| Pogo | 118259600 | ORU81480 | Gammaproteobacteria | Thiotrichales | Francisellaceae | Francisella_tularensis | Soil, mammal pathogen | No TIRs, partial sequence 1135 bp from natural spring water | A few matches with *Verticillium dahliae* (fungus) ~75% query cover and 73% id | A lot of matches with fungi >95% query cover and >59% id | No - Probably contamination |
| Sola1 | 149130375 | RLI68114 | Candidatus Heimdallarchaeota | - | - | Candidatus_Heimdallarchaeota_archaeon | Hydrothermal sediment | No TIRs, full sequence 4433 bp from deep-sea hydrothermal vent sediments | No matches | A few matches with bacterial species (but not Archeae) ~80-90% query cover and 60-80% id | No - Probably contamination |
| Sola1 | 140139201 | PYY20342 | - | - | - | - | Various environmental | No TIRs, full sequence 3259 bp from soil metagenome | No matches | Matches with *Rhizophagus irregularis* (fungus) 99% query cover and ~60% id | No - Probably contamination |
| Sola1 | 57178417 | ETR66427 | Deltaproteobacteria | Desulfobacterales | Desulfobacteraceae | Candidatus_Magnetoglobus_multicellularis | Hypersaline lagoon, multicellular | No TIRs, full sequence 1573 bp from sediment and water | No matches | Matches with other *Magnetobulus* bacteria and from limpet ~70% query cover and ~53% id | No - low % id and query cover |
| Sola1 | 57178632 | ETR67790 | Deltaproteobacteria | Desulfobacterales | Desulfobacteraceae | Candidatus_Magnetoglobus_multicellularis | Hypersaline lagoon, multicellular | No TIRs, full sequence 2760 bp from sediment and water | No matches | No match >50% id | No |
| Sola1 | 92767425 | KPA10833 | Deltaproteobacteria | Desulfobacterales | Desulfobacteraceae | Candidatus_Magnetomorum_sp._HK-1 | Coastal tidal sand flats | No TIRs, partial sequence 677 bp from sea sediments | No matches | No match >50% id | No |
| Sola1 | 57178442 | ETR66575 | Deltaproteobacteria | Desulfobacterales | Desulfobacteraceae | Candidatus_Magnetoglobus_multicellularis | Hypersaline lagoon, multicellular | No TIRs, partial chaperonin sequence 1649 bp from sediment and water | No matches | Several matches with other bacteria, 66-99% query cover and 50-64% id | No - other bacterial species and low %id |
| Sola1 | 57178476 | ETR66776 | Deltaproteobacteria | Desulfobacterales | Desulfobacteraceae | Candidatus_Magnetoglobus_multicellularis | Hypersaline lagoon, multicellular | No TIRs, chaperonin sequence 1824 bp from sediment and water | No matches | Several matches with other bacteria, 66-99% query cover and 50-70% id | No - other bacterial species and low %id |
| Sola1 | 118058364 | WP084069893 | Deltaproteobacteria | Desulfobacterales | Desulfobacteraceae | Desulfobacterium_vacuolatum | Marine sediment | Nucleotidic corresponding sequence not available | Nothing with tBLASTn | A few matches with other bacterial species with 68-92% query cover and 74-89%id | No - other bacterial species |
| Sola1 | 107285359 | OEU55806 | Deltaproteobacteria | Desulfobacterales | Desulfobacterales | Desulfobacterales | Marine sediment | 35 bp TIRs, full sequence 2213 bp from marine sediment metagenome | No matches | A few sequences from other bacteria 61-98% query cover and 50-81% id | No - other bacterial species |
| Sola1 | 108536325 | OGV34713 | Gammaproteobacteria | Legionellales | Legionellales | Legionellales_bacterium_RIFCSPHIGHO2_12_FULL_35_11 | Various environmental, pathogens | 39 bp TIRs, full sequence 3566 bp from subsurface metagenome | No match >50% query cover (around 2-4% only) | A few sequences from other bacteria >95% query cover and 50-57% id | No - other bacterial species and low %id |
| Sola1 | 118263510 | ORU94837 | Gammaproteobacteria | Thiotrichales | Piscirickettsiaceae | Cycloclasticus_sp._symbiont_of_Poecilosclerida_sp._M | Symbiont_of_Poecilosclerida_sp (sponge), marine sediments | No TIRs, full sequence 4104 bp from *Poecilosclerida* sp. (symbiont) | No matches | No match >50% id | No |
| Sola1 | 118263529 | ORU94956 | Gammaproteobacteria | Thiotrichales | Piscirickettsiaceae | Cycloclasticus_sp._symbiont_of_Poecilosclerida_sp._M | Symbiont_of_Poecilosclerida_sp (sponge), marine sediments | No TIRs, full sequence 3314 bp from *Poecilosclerida* sp. (symbiont) | No matches | Matches with other sponges and a few other invertebrates but ~70% query cover and 50-55% id | No - Probably contamination |
| Sola1 | 15207018 | EDN68038 | Gammaproteobacteria | Thiotrichales | Thiotrichaceae | Beggiatoa_sp._PS | Sulfur-rich environments | 23 bp TIRs, full sequence 2156 bp from purified filament at 4m deep sediments in Baltic sea | No matches | Several matches from other bacteria 58-100% query cover and 50-75% id | No - other bacterial species |
| Sola1 | 15207465 | EDN71489 | Gammaproteobacteria | Thiotrichales | Thiotrichaceae | Beggiatoa_sp._PS | Sulfur-rich environments | 31 bp TIRs, full sequence 13628 bp from purified filament at 4m deep sediments in Baltic sea | No matches | Several matches from other bacteria >95% query cover and 52-80% id | No - other bacterial species |
| Sola1 | 102631477 | OAD19029 | Gammaproteobacteria | Thiotrichales | Thiotrichaceae | Candidatus_Thiomargarita_nelsonii | Hydrogen sulfide rich environments | No TIRs, full sequence 1939 bp from single cell marine sediments | No matches | only one match ~50% query cover and id from Candidatus *Magnetoglobus* | No - low query cover |
| Sola1 | 72328351 | KHD04931 | Gammaproteobacteria | Thiotrichales | Thiotrichaceae | Candidatus_Thiomargarita_nelsonii | Hydrogen sulfide rich environments | 75 bp TIRs, full sequence 3672 bp from *Provanna laevis* (symbiont) | No matches | A few matches with other bacteria 83-99% query cover and ~50% id | No - other bacterial species |
| Sola1 | 72328268 | KHD04346 | Gammaproteobacteria | Thiotrichales | Thiotrichaceae | Candidatus_Thiomargarita_nelsonii | Hydrogen sulfide rich environments | Record removed (mastered by WGS sequencing) | - | - | - |
| Sola2 | 156656929 | RYA67237 | Gammaproteobacteria | Enterobacterales | Enterobacteriaceae | Enterobacter_cloacae_complex_sp._4DZ3-17B2 | Mammal gut | No TIRs, partial sequence 362 bp from cloaca swabs in chicken farm | No match >50% query cover | Several matches with ant species 54-100% query cover and 50-81M id | No - Probably contamination |
| Sola3 | 154390470 | RUA04805 | Gammaproteobacteria | - | - | - | Various environmental, phytopathogens, animal pathogens | No TIRs, full sequence 6605 bp from hydrothermal vent metagenome | No matches with either of the flanks | A few matches with cnidarians 80-89% query cover and 50-69% id | No - Probably contamination |
| Sola3 | 154390464 | RUA04752 | Gammaproteobacteria | - | - | - | Various environmental, phytopathogens, animal pathogens | No TIRs, full sequence 4416 bp from hydrothermal vent metagenome | No matches with either of the flanks | Matches with cnidarians 60-98% query cover and 50-68% id | No - Probably contamination |
| Tc1/Mar | 119431244 | OUB07123 | Bacilli | Bacillales | Bacillaceae | Bacillus thuringiensis | Soil, pathogen of invertebrates | 59 bp TIRs, full sequence 1503 bp from *Bacillus thuringiensis* soil sample | A lot of matches with Tc1 transposon in fish species, >80% query cover and id | A few matches with insects, fish and cnidarians 50-99% query cover and 50-100% id | No - Probably contamination |
| Tc1/Mar | 120765629 | WP088017964 | Bacilli | Bacillales | Bacillaceae | Bacillus thuringiensis | Soil, pathogen of invertebrates | Protein record suppressed, no longer annotated on any genome | - | - | - |
| Tc1/Mar | 159591561 | WP133372499 | Bacilli | Bacillales | Bacillaceae | Bacillus aciditolerans | Soil, pathogen of invertebrates | This sequence is only available in amino acid (NR record) and is 155 aa | tBLASTn retrieved no results | A lot of matches with *Myotis myotis* (not a bacteria), then a few arthropod species >80% query cover and 50-66% id | No - Probably contamination |
| Tc1/Mar | 140485463 | PZO92957 | Bacilli | Lactobacillales | Streptococcaceae | Streptococcus pyogenes | Pathogen of mammals | 21 bp truncated TIRs, full sequence 1368 bp from indoor metagenome (hospital surfaces) | No matches | No match > 50% query cover | No |
| Tc1/Mar | 156133523 | RXF71996 | Deltaproteobacteria | Desulfovibrionales | Desulfovibrionaceae | Desulfovibrio sp. DS-1 | Marine sediment | No TIRs, full sequence 1092 bp from soil | Matches with insects 51-74% query cover and >85% id | A lot of matches with other bacteria >90% query cover and id | No - other bacterial species and insects with low query cover |
| Tc1/Mar | 156133650 | RXF73229 | Deltaproteobacteria | Desulfovibrionales | Desulfovibrionaceae | Desulfovibrio sp. DS-1 | Marine sediment | 19 bp TIRs, ful sequence 2366 bp from soil | No matches | A lot of matches with other bacteria and insects, >70% query cover and >50% id | No - other bacterial species and insects with low percent id |
| Tc1/Mar | 113089697 | WP075666983 | Gammaproteobacteria | Enterobacterales | Enterobacteriaceae | Klebsiella pneumoniae | Mammal symbiont | Protein record suppressed, no longer annotated on any genome | - | - | - |
| Tc1/Mar | 156867418 | WP129321181 | Gammaproteobacteria | Enterobacterales | Enterobacteriaceae | Enterobacter cloacae complex sp. 4DZ3-17B2 | Mammal gut | This sequence is only available in amino acid (NR record) and is 354 aa | tBLASTn: matches with insects (mostly ants), >50% query cover and 50-71% id | Matches with insects (mostly ants), >50% query cover and 50-86% id | No - Probably contamination |
| Tc1/Mar | 64763022 | WP025903236 | Gammaproteobacteria | Enterobacterales | Erwiniaceae | Tatumella sp. UCD-D suzukii | Drosophila symbiont | Protein record suppressed, no longer annotated on any genome | - | - | - |
| Tc1/Mar | 109853523 | WP071226877 | Gammaproteobacteria | Pseudomonadales | Moraxellaceae | Acinetobacter baumannii | Various environmental, mammal pathogen | This sequence is only available in amino acid (NR record) and is 134 aa | tBLASTn: matches with moths maT/mariner-like TEs, >90% query cover and id | A lot of matches with other bacteria, then a few from lepidopterans, 60-90% query cover and ~94% id | No - Probably contamination |
| Tc1/Mar | 109843414 | WP071212576 | Gammaproteobacteria | Pseudomonadales | Moraxellaceae | Acinetobacter baumannii | Various environmental, mammal pathogen | Protein record suppressed, no longer annotated on any genome | - | - | - |
| Tc1/Mar | 109871405 | WP071290505 | Gammaproteobacteria | Pseudomonadales | Moraxellaceae | Acinetobacter baumannii | Various environmental, mammal pathogen | This sequence is only available in amino acid (NR record) and is 346 aa | tBLASTn: matches with *Bombyx* Bmmar TEs, 100% query cover and >80% id | A lot of matches with other bacteria, then from insects, >90% query cover and id | No - Probably contamination |
| Tc1/Mar | 117102800 | WP080578147 | Gammaproteobacteria | Pseudomonadales | Pseudomonadaceae | Pseudomonas monteilii | Various environmental, mammal pathogen | This sequence is only available in amino acid (NR record) and is 216 aa | tBLASTn: matches with insects and mammals, >85% query cover and >65% id | A lot of matches with other bacteria, then from insects, >90% query cover and id | No - Probably contamination |
| Tc1/Mar | 133495367 | WP103100911 | Gammaproteobacteria | Xanthomonadales | Xanthomonadaceae | Xanthomonas citri | Phytopathogen | This transposase sequence is only available in amino acid (NR record) and is 350 aa | tBLASTn: matches with insects SETmar-like, >80% query cover and >50% id | A lot of matches with insects, then from other bacteria, >80% query cover and >50% id | No - Probably contamination |

Table S7. Numbers of stop codons present in transposases for each DDE transposon superfamily

| Superfamily | No. transposases | No. transposases with a stop | No. stops | Mean stops | Proportion with stop | Percentage with stop |
|---|---|---|---|---|---|---|
| Academ | 234 | 23 | 122 | 5.3 | 0.098290598 | 9.8 |
| CMC | 1085 | 20 | 41 | 2.1 | 0.01843318 | 1.8 |
| Ginger | 1079 | 9 | 13 | 1.4 | 0.008341057 | 0.8 |
| GingerRoot | 1186 | 36 | 40 | 1.1 | 0.030354132 | 3.0 |
| hATs | 4714 | 113 | 399 | 3.5 | 0.02397115 | 2.4 |
| Kolobok | 809 | 2 | 4 | 2.0 | 0.002472188 | 0.2 |
| Merlin | 1509 | 11 | 16 | 1.5 | 0.007289596 | 0.7 |
| Mutator | 4536 | 179 | 420 | 2.3 | 0.039462081 | 3.9 |
| P | 193 | 7 | 26 | 3.7 | 0.03626943 | 3.6 |
| PHIS | 5738 | 130 | 255 | 2.0 | 0.022655978 | 2.3 |
| Piggybac | 1351 | 29 | 52 | 1.8 | 0.021465581 | 2.1 |
| Pogo | 1775 | 21 | 64 | 3.0 | 0.011830986 | 1.2 |
| Sola1 | 760 | 4 | 9 | 2.3 | 0.005263158 | 0.5 |
| Sola2 | 272 | 4 | 7 | 1.8 | 0.014705882 | 1.5 |
| Sola3 | 167 | 47 | 80 | 1.7 | 0.281437126 | 28.1 |
| Tc1mar | 3052 | 28 | 40 | 1.4 | 0.009174312 | 0.9 |
| Zator | 368 | 0 | 0 | - | 0 | 0.0 |
| Transib | 35 | 0 | 0 | - | 0 | 0.0 |
| Novosib | 10 | 0 | 0 | - | 0 | 0.0 |
| | 28873 | 663 | 1588 | 2.3 | 0.033232444 | 3.3 |
